# Free energy spectroscopy reveals the mechanistic landscape of chromatin compaction

**DOI:** 10.1101/2025.08.13.670082

**Authors:** Kalven Bonin, Yuchen Wang, Ivan Riveros, Ruo-Wen Chen, Bin Zhang, Ralf Bundschuh, Carlos E. Castro, Michael G. Poirier

**Author notes:** Correspondence (C.E.C.), (M.G.P).

## Abstract

Eukaryotic genomic DNA is repeatedly wrapped into nucleosome spools: the basic building block of chromatin. This organization regulates the physical accessibility of the genome to gene transcription, replication, and repair regulatory factors. Chromatin compaction is controlled by multivalent weak interactions, resulting in a complicated conformational landscape that remains challenging to characterize. This work reports a method for characterizing chromatin compaction, Free Energy Spectroscopy (FES), which is based on DNA nanotechnology and transmission electron microscopy. This method experimentally determines the chromatin compaction free energy landscape in terms of end-to-end distance and nucleosome stacking interactions. By deconvolving the free energy landscapes of partially and fully compact tetranucleosomes, FES revealed three separate mechanisms by which linker histones reshape the compaction energetics to condense chromatin. This study establishes FES as a method with the potential to help answer a broad range of mechanistic questions about genome and epigenome function.

## INTRODUCTION

Chromatin structural dynamics is a key regulator of accessibility for gene transcription, repair, and replication. For example, transcription regulatory complexes access DNA within open euchromatin but are excluded from compact heterochromatin^1,2^. An important level of chromatin compaction is clusters of four nucleosomes, i.e. tetranucleosomes. They form *in vivo* to help form chromatin clutches that regulate transcription^3,4^ and *in vitro* within longer nucleosome arrays^5–7^. This tetranucleosome unit of chromatin is compacted by magnesium ions^8^ and the linker histone H1, an abundant chromatin architectural protein that binds near the nucleosome dyad region^9^. H1 regulates tetranucleosome compaction by influencing the extent to which DNA is wrapped into nucleosomes^10,11^ and the stacking between nucleosomes^12,13^, which are two essential differences between euchromatin and heterochromatin.

The folded structure of the tetranucleosome has been investigated *in vitro* by x-ray crystallography^14^, cryo-electron microscopy (cryo-EM)^5^, cysteine crosslinking^12^, and ensemble fluorescence^15^. These studies provide high-resolution information about static structures and have revealed that the primary folded structure consists of a zig-zag pattern with two nucleosome-nucleosome stacking interactions (NSI) between nucleosomes 1-3 and 2-4^14^. Stopped flow^16^ and single-molecule^6^ fluorescence studies have provided kinetic information about the time to transition between different compacted states, while single-molecule force spectroscopy^17–19^ has provided information about discrete free energy differences between distinct compacted states.

Importantly, chromatin is a continuum of conformational states, where the relative probability of open and compact chromatin states is determined by the free energy difference between each of the states through the Boltzmann distribution. A comprehensive description of chromatin compaction is a full free energy landscape describing the continuum of states, which has currently only been predicted by computational modeling^20–24^. The ability to experimentally determine chromatin compaction free energy landscapes would open a new level of understanding of how chromatin controls genome accessibility. DNA nanotechnology has the potential to provide an approach to probe tetranucleosome structure and dynamics, including measuring the free energies associated with compaction, while being imaged by transmission electron microscopy (TEM) to elucidate the corresponding tetranucleosome conformation^25,26^.

Scaffolded DNA origami is a powerful approach in the field of DNA nanotechnology, which utilizes a long single-stranded DNA (ssDNA) scaffold and numerous short ssDNA oligonucleotides that base-pair together to construct functional nanostructures^27–29^ that can be designed to be stable under physiological conditions^30^. DNA-based nanostructures have been used as probes to detect and quantify many biomolecular interactions^25^. For example, DNA-based nanomechanical devices have quantified DNA bending^31^, DNA shearing^32^, DNA-protein interactions^33^, histone octamer stacking^34^, and nucleosome unwrapping^30,35^

In this work, we use the recently established set of nanocalipers: nanoscale DNA Force Spectrometers (nDFSs)^31,32^, to study tetranucleosome compaction. These nDFS devices consist of two caliper arms, which are approximately 50 nm in length, and have an 8x3 cross-section of double-stranded DNA (dsDNA) helices with some helices removed in the middle layer. The two stiff arms of the nDFS nanocalipers are connected by both short (2 bases) and long (70 bases) ssDNA loops. By introducing additional DNA base pairing within the long ssDNA loops, the angular free energy landscape can be modulated. After incorporating a tetranucleosome between the two nanocaliper arms, the end-to-end distance, *ℓ*, can be used as a natural “reaction coordinate” for the compaction free energy landscape, which relates to the angular free energy landscape through the arm length of the nanocaliper. Three different nanocaliper designs are used here: nDFS.A, nDFS.B, and nDFS.C^31^, allowing for a wide range of tetranucleosome end-to-end distances to be probed.

By using these nDFS nanocalipers, we experimentally determine the compaction free energy landscapes of tetranucleosomes as a function of end-to-end distance. We find that the free energy landscape of Mg^2+^ compacted tetranucleosomes has a minimum free energy plateau over a broad range of end-to-end distances. This indicates Mg^2+^ compacted tetranucleosomes have significant conformational freedom. Above about 30 nm, the free energy increases linearly by about 4 k_B_T as the end-to-end distance is doubled, revealing the energetic cost for extending tetranucleosomes. The human linker histone, H1.0, converts the free energy plateau to a 3 k_B_T free energy well with a free energy minimum at an end-to-end distance of about 5 nm, revealing how H1.0 alters the compaction free energy landscape to regulate tetranucleosome compaction. Because the free energy landscapes are determined from TEM images, the number of nucleosome stacking interactions (NSIs) can also be determined for each particle. This allows for determining a two-dimensional free energy landscape in terms of the end-to-end distance and the stacking state. This multidimensional analysis reveals the energetic cost and changes in tetranucleosome conformational freedom as one nucleosome stacking interaction is disrupted. These results provide a new mechanistic picture of how tetranucleosomes are “opened” for chromatin regulatory factors. Overall, this approach for determining tetranucleosome compaction free energy landscapes, which we refer to as Free Energy Spectroscopy (FES), is positioned to provide important insight into the mechanisms of a wide range of chromatin compaction regulatory factors and more generally into conformational transitions of biomolecular complexes at the 10-100nm scale.

## DESIGN

FES is an experimental approach for determining the conformational free energy landscape of large macromolecular complexes, such as tetranucleosomes, which is the focus of this study (**Figure 1**). The approach relies on DNA origami nanocalipers to define the conformational reaction coordinate, which here is the tetranucleosome end-to-end distance, *ℓ*. The conformations of the nanocalipers with and without single tetranucleosomes are quantified from TEM images, allowing facile determination of tetranucleosome end-to-end distance *ℓ* free energy landscapes. By using a set of nanocalipers with various angular distributions, different regions of the end-to-end distance *ℓ* free energy landscape are quantified. These landscapes are then combined to determine the free energy landscape over a wide range of end-to-end distances *ℓ*. The number of nucleosome stacking interactions NSIs are also identified from the same TEM images, allowing a second reaction coordinate to be determined. This enables FES to measure the shape of multidimensional tetranucleosome compaction free energy landscapes. These measurements can then be done with chromatin regulator factors such as the linker histone H1.0, which is a focus of this study, to determine how the factor reshapes the tetranucleosome conformational landscape. Given that these DNA nanocalipers are straightforward to prepare and the wide availability of TEM instrumentation, FES is positioned to investigate mechanisms of how a wide range of regulators function to control the conformational landscapes of not only chromatin, but of large macromolecular complexes in general such as ribonucleoprotein complexes and even protein condensates.

**Fig 1:**
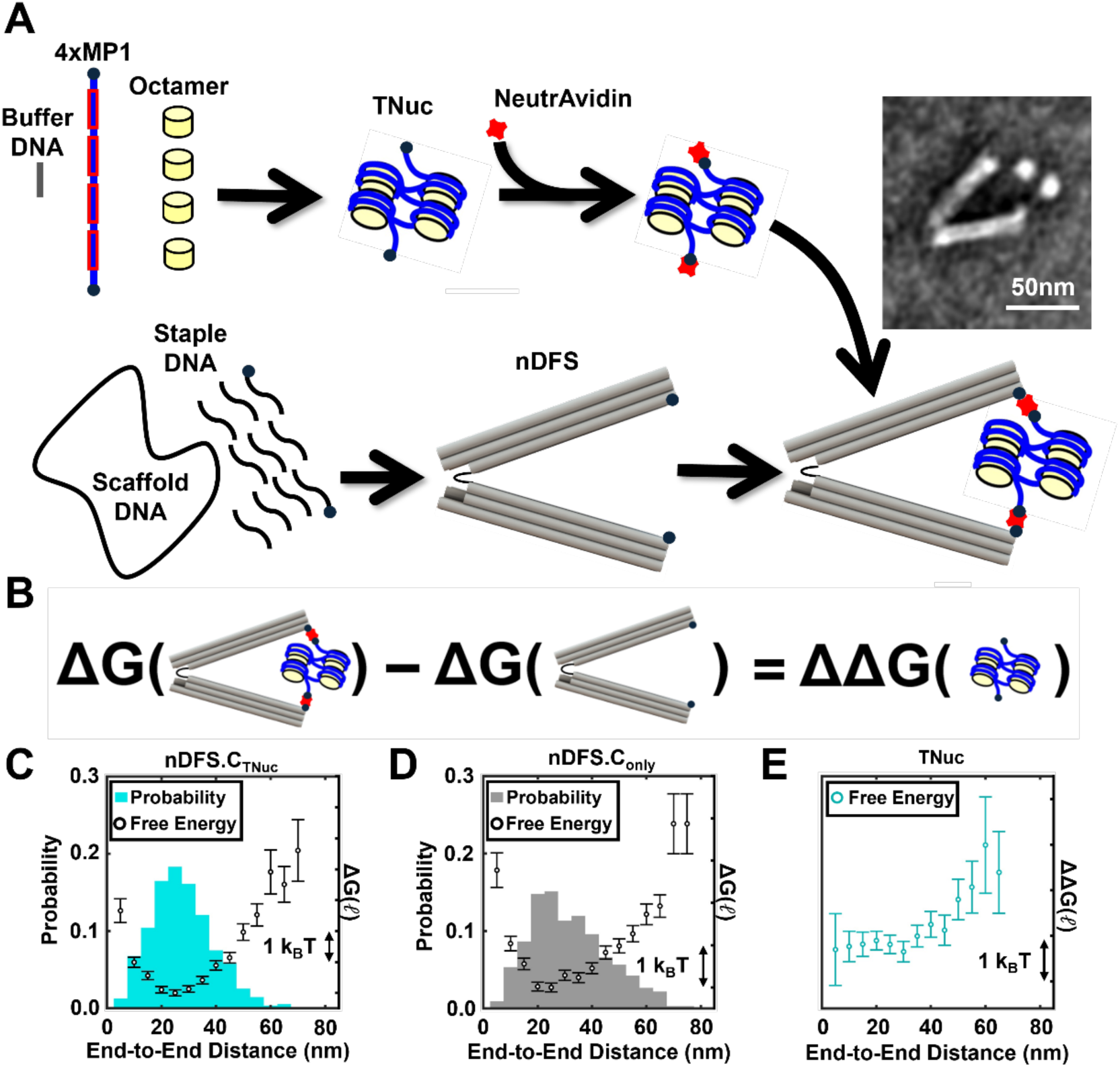
Free energy measurement strategy. (**A**) Schematic of tetranucleosome incorporation into the nDFS devices, including DNA origami nanocaliper folding and tetranucleosome reconstitution. (Inset) TEM image of a properly incorporated Mg^2+^ compacted tetranucleosome into a nanocaliper. (**B**) Schematic representation showing the process for computing the tetranucleosome free energy as a function of end-to-end distance from free energy measurements. (**C**-**D**) End-to-end probability distributions (bars) and corresponding free energy landscapes (open circles) from nanocalipers with tetranucleosomes (nDFS.C_TNuc_) and (D) free nanocalipers (nDFS.C_only_). (**E**) End-to-end tetranucleosome free energy landscape. The error bars of the free energy values were propagated from the uncertainties of the probability measurements, which were estimated by the square root of the number of events in each bin.

## RESULTS

### DNA nanocalipers directly measure the compaction free energy landscape of tetranucleosome compaction

To investigate tetranucleosome free energy landscapes, we developed a DNA nanotechnology approach (**Figure 1**), building on previous measurements of single nucleosomes^30,35^ and histone octamers^34^. We used a set of nDFS nanocalipers: nDFS.A, nDFS.B, and nDFS.C, which were previously developed to study dsDNA bending and DNA wrapping into nucleosomes^31^. Both the nDFS nanocalipers^31^ and tetranucleosome arrays^16,36^ were prepared and validated as previously described. The tetranucleosome DNA template contained a 4x repeat of a high-affinity nucleosome positioning sequence (NPS), mp1^36^, which is a variant of the Widom 601 NPS^37^, which were spaced by 30 bp of linker DNA and flanked by 50 bp of DNA. Biotins were included at each 5-prime end of the tetranucleosome DNA and the end of each nanocaliper arm. NeutrAvidin was then used to attach a single tetranucleosome between the two nanocaliper arms (**Figure 1**).

For the initial proof-of-principle measurement, we used nDFS.C because its end-to-end distance distribution overlapped with the typical size of compacted tetranucleosomes (**Figure S1**), which we anticipated would help incorporation efficiency. Tetranucleosomes were integrated into the nDFS device by combining them at an equimolar concentration at room temperature in physiological ionic conditions (200mM NaCl and 1mM MgCl_2_, **Figure 1, S2**) and then imaged using negative stain TEM. These integration conditions resulted in 68±3% of the nDFS.C devices having a tetranucleosome properly integrated (**Figure S3**). Fixation of samples with conditions used in previous TEM studies of chromatin^38^ prevented the disruption of nucleosome stacking without inducing tetranucleosome compaction (**Figure S4-S5**) or altering the end-to-end distance distribution of the nDFS.C device (**Figure S6)**. Regions of interest within TEM images containing a single nDFS.C device were selected and analyzed with imageJ to quantify the end-to-end distance distribution of the nDFS.C device with and without tetranucleosomes (See Star Methods for details).

The end-to-end distance (*ℓ*), measured as the distance between the inner tip of each caliper arm of each device (**Figure 1A**), was determined for 546 nDFS.C nanocalipers with properly incorporated tetranucleosomes (nDFS.C_TNuc_) and 635 nDFS.C nanocalipers that were not incubated with tetranucleosomes (nDFS.C_only_) (**Table S1)**. The ensemble of *ℓ* values was used to determine the end-to-end distance probability distribution, P(*ℓ*), (**Figure 1C-D**). By assuming that the ensemble is in thermal equilibrium, the free energy landscape as a function of *ℓ* can be calculated using the Boltzmann probability, ΔG = - k_B_T ln (P(*ℓ*) / P_ref_). P(*ℓ*) is the probability for a given end-to-end distance *ℓ*, while P_ref_ is the probability of the reference state (**Figure 1C-D**). Since we are only interested in differences in free energy, the choice of P_ref_ does not influence the shape of the landscape. We therefore plot ΔG without indicating ΔG=0 where the graph’s free energy tick marks indicate a ΔG=1 k_B_T.

When the tetranucleosome is properly attached to a nanocaliper, visual inspection of the images reveals that the nucleosomes are not in direct contact with the nanocaliper beyond the biotin-Neutravidin-biotin interactions (**Figure S2**). This implies that free energy landscapes of the tetranucleosome alone and the nanocaliper alone are additive, where the sum equals the free energy landscape of the nanocaliper-tetranucleosome complex. Therefore, the free energy landscape of the tetranucleosome alone can be calculated from the difference between the free energies of the nanocaliper-tetranucleosome complex and the nanocaliper alone, i.e. ΔΔG(*ℓ*)_TNuc_ = ΔG(*ℓ*)_nDFS+TNuc_ - ΔG(*ℓ*)_nDFS_ (**Figure 1B**). Using this approach to determine the tetranucleosome compaction free energy landscape ΔΔG(*ℓ*)_TNuc_, we find that the tetranucleosome free energy does not change significantly for end-to-end distances *ℓ* less than about 30 nm, while above 30 nm the free energy rises linearly (**Figure 1C-E**). These results demonstrate that DNA origami nanocalipers can be used as a probe to measure the continuous free energy landscape of the tetranucleosome in terms of its end-to-end distance *ℓ*. To our knowledge, this is the first experimental measurement of a chromatin compaction free energy landscape.

### Combined measurements with different DNA nanocalipers significantly improve the precision and dynamic range of the tetranucleosome free energy landscape

While the nDFS.C nanocaliper enabled the measurement of the free energy landscape of tetranucleosome compaction, the range of end-to-end distances for which the free energy can be accurately measured is limited (**Figure 1E**). This is because only end-to-end distances that are thermally accessible can be observed and quantified. We are using about 500 conformation measurements to quantify a free energy landscape with a single device and require at least 3 events in an end-to-end distance bin to be included in the probability distribution. Therefore, detectable free energy differences are limited to about k_B_T ln(500/3) ∼ 5 k_B_T. To expand the end-to-end distance range of the measurement, we used multiple nDFS devices, each with a different free energy landscape, to bias the tetranucleosome to varying ranges of conformations. This approach is analogous to umbrella sampling, an approach widely used in molecular dynamics simulations of bimolecular complexes^39,40^, where a known energy potential (typically harmonic) is added to bias conformational distributions. This allows for low probability states to be sufficiently populated for quantification. The applied energy potential is then removed after the simulation. By applying multiple energy potentials where each bias over a different range of states, free energy landscapes over a wider range of conformations can be quantitatively studied and reconstructed.

Here, we used 2 additional nDFS devices, nDFS.A and nDFS.B, which were designed with larger mean opening angles^31^ by modifications to the vertex within the nDFS nanocaliper design (**Figure S7**). The nDFS.A and nDFS.B free energy landscapes have minima at *ℓ* = 55nm and 65 nm, respectively, which are larger than 25nm, the end-to-end distance *ℓ* for the nDFS.C free energy minimum (**Figure 2A-C**). This should allow nDFS.A and nDFS.B to probe tetranucleosome conformations corresponding to larger end-to-end distances. We carried out free energy measurements of tetranucleosomes with both nDFS.A and nDFS.B, as we did with nDFS.C. We found the percentage of nDFS.A and nDFS.B devices that contained properly integrated tetranucleosomes were 45±2% and 27±2%, respectively (**Figure S3**). These are lower than the integration efficiency into nDFS.C, which is expected since the tetranucleosome end-to-end distance *ℓ* is less likely to match the end-to-end distance *ℓ* of either nDFS.A and nDFS.B than of nDFS.C.

**Fig 2:**
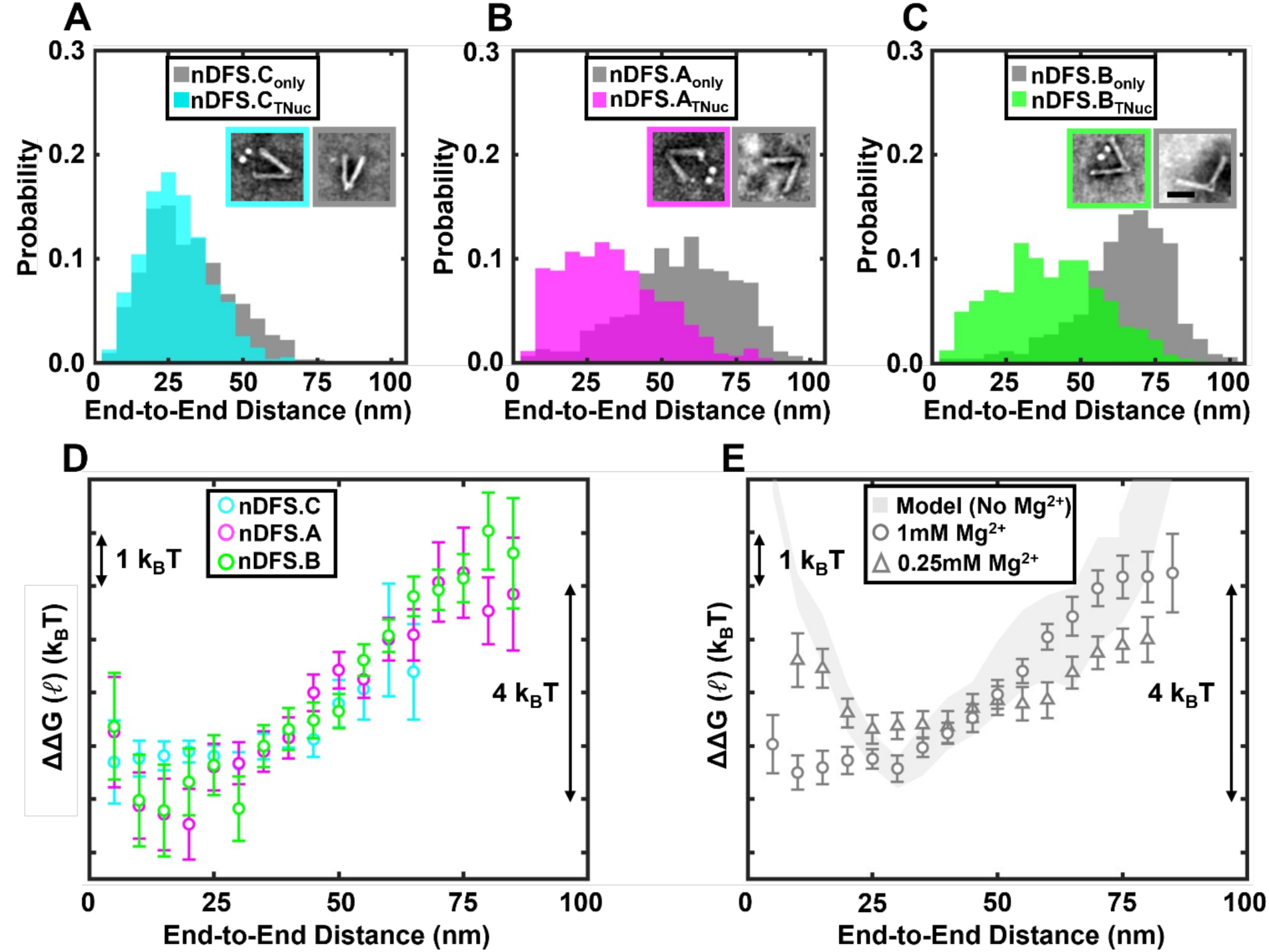
The application of multiple nanocaliper designs increases the precision and dynamic range of the free energy measurement. (**A**-**C**) End-to-end distance probability distributions for nanocalipers are shown in grey for each nanocaliper design (nDFS.C_only_, nDFS.A_only_, nDFS.B_only_). End-to-end distance probability distributions of nanocalipers with tetranucleosomes (nDFS.C_TNuc_, nDFS.A_TNuc_, nDFS.B_TNuc_) are shown in (A) cyan, (B) magenta, and (C) green, respectively. Insets show examples of images used to determine each probability distribution. The black scale bar in (C) is 50nm. (**D**) End-to-end distance tetranucleosome free energy landscapes from each nanocaliper design. The error bars of the free energy values were propagated from the uncertainties of the probability measurements, which were estimated by the square root of the number of events in each bin. (**E**) End-to-end distance tetranucleosome free energy landscapes determined from the weighted average of the three independent free energy measurements using the three different nanocalipers in 1mM Mg^2+^ (open grey circles) and 0.25 mM Mg^2+^ (open grey triangles). The error bars of the free energies were determined as described in the Star Methods section. The free energy landscape derived from molecular dynamics simulations of the coarse-grained model is plotted as the shaded grey area, which represents the 95% confidence interval.

We collected TEM images of the nDFS.A and nDFS.B nanocalipers alone (nDFS.A_only_ and nDFS.B_only_) and with tetranucleosomes (nDFS.A_TNuc_ and nDFS.B_TNuc_) (**Table S1**). Regions of interest containing individual nanodevices were selected (**Figure S8-S11**) and then analyzed to determine end-to-end distance probability distributions P(*ℓ*) for each device (**Figure 2A-C**, insets) as described above. Visual inspection of each probability distribution P(*ℓ*) reveals the integration of tetranucleosomes into nDFS.A and nDFS.B resulted in a shift in tetranucleosome conformations corresponding to larger end-to-end distances *ℓ* compared to nDFS.C. Furthermore, the probability distributions P(*ℓ*) with nDFS.A_TNuc_ and nDFS.B_TNuc_ spans a much wider range of end-to-end distances *ℓ* than the probability distributions P(*ℓ*) with nDFS.C_only_, which is consistent with wider angle distributions of nDFS.A and nDFS.B. These observations indicate that nDFS.A and nDFS.B extend the tetranucleosome end-to-end distances *ℓ* more than nDFS.C.

We separately determined the free energy landscape ΔΔG(*ℓ*)_TNuc_ using nDFS.A and nDFS.B, as previously described for nDFS.C (**Figure 2D**). As overall offsets to the free energy landscape are arbitrary, a constant free energy offset needs to be subtracted from each landscape to compare the three free energy landscape ΔΔG(*ℓ*)_TNuc_ measurements from the three different nanocalipers. To determine the offset, a loss function with two constant offsets for ΔΔG(*ℓ*)_TNuc_ from nDFS.A and nDFS.B was minimized, where the uncertainty of the measurements for each device was taken into account (see Star Methods for details). The three independent free energy landscape measurements show good agreement within the overlapping ranges of end-to-end distance *ℓ*, indicating that this method can reliably measure the free energy landscape ΔΔG(*ℓ*)_TNuc_.

Now that we have established that the free energy landscape ΔΔG(*ℓ*)_TNuc_ can be measured by three distinct nanocalipers, we combined these three measurements (**Figure 2E**) by computing the weighted average of ΔΔG(*ℓ*)_TNuc_ at each value of *ℓ* (see Star Methods for details). This increased the range of end-to-end distances *ℓ* to extend from 5 nm to 85 nm, while having a free energy precision of about 1 k_B_T over a range of about 4 k_B_T. The resolution of the end-to-end distance is set by our bin size of 5 nm. This bin size was used as a good tradeoff between spatial resolution and maintaining a smooth probability distribution given the total number of measurements. This bin size is larger than the precision of an individual end-to-end distance measurement of a nanocaliper, which we estimate to be about 1 nm. (**Figure S12**)

With these improvements, distinct regions within the free energy landscape ΔΔG(*ℓ*)_TNuc_ are better resolved. At small values of end-to-end distance *ℓ*, 5-30 nm, the free energy landscape ΔΔG(*ℓ*)_TNuc_ is constant, implying a free energy plateau. This indicates that within this range of *ℓ*, there is no free energy difference between these levels of tetranucleosome compaction, implying it can fluctuate freely over a range of compaction. At larger values of *ℓ*, >30 nm, the free energy increases by 4 k_B_T, implying that it is energetically costly to extend the tetranucleosomes over these larger end-to-end distances.

Overall, these results demonstrate that this DNA nanotechnology-based method can be used to determine a continuous free energy landscape, which is an approach for characterizing tetranucleosome compaction. The approach leverages the tunability of the DNA origami nanocalipers where measurements with different vertex designs are combined to provide significantly greater precision and dynamic range. This technique directly measures a spectrum of free energies through probabilities, which is why we refer to this approach as Free Energy Spectroscopy (FES). The measured free energy landscapes are rich with mechanistic details, including revealing that tetranucleosomes have distinct regions of compaction, which is important for understanding how chromatin compaction regulates genome accessibility. Finally, these results provide a critical baseline for measurements of chromatin regulatory factors and how they function to regulate chromatin compaction.

### The measured compaction free energy landscape is consistent with an independently determined coarse-grained computational model

To validate our measured free energy landscape, we computed the free energy profiles of the tetranucleosome using umbrella sampling simulations based on coarse-grained protein-DNA models. We employed the 3SPN.2C DNA model^41^, which represents each nucleotide with three interaction sites, and the SMOG2 protein model^42^ to describe the histone components. Additional protein-DNA interactions were incorporated through electrostatic and Lennard-Jones potentials. This modeling framework has been previously validated in studies of nucleosome DNA unwrapping^43^, the force-extension behavior of chromatin fibers^44^, and the folding pathways of tetra-nucleosome assemblies^21,24^.

Our simulations exhibit good agreement with the magnesium compacted tetranucleosome free energy landscape for end-to-end distances exceeding 25 nm (**Figure 2E**). However, at distances of less than 25 nm, the computed free energy profile diverges. One plausible explanation for this divergence is the implicit treatment of divalent ions in our simulations. Under compact configurations, increased interactions of divalent ions with chromatin may further attenuate electrostatic repulsion among DNA segments, thereby reducing the energetic barrier^23^.

We therefore determined the tetranucleosome free energy landscape with 4-fold less Mg^2+^ (0.25 mM). Since Mg^2+^ helps stabilize the nDFS structures, we confirmed by TEM imaging that the nDFS structures remained well folded during the experiment in 0.25 mM Mg^2+^ (**Figure S13**). We then used nDFS.C, nDFS.A and nDFS.B to determine the tetranucleosome free energy landscape at 0.25 mM Mg^2+^ (**Figure 2E, S14-S16**). We find that at distances of less than 25 nm the free energy landscape increases similarly to the simulation results. Given that the model parameters were derived independently of the experimental data, this agreement strongly supports the quantitative reliability of our approach in probing the energy landscape of chromatin folding.

### The linker histone H1.0 compacts tetranucleosomes by locally biasing the free energy landscape to the higher compacted states through converting the short-range plateau to an energy well

After establishing the FES method for investigating magnesium-mediated tetranucleosome compaction, we used FES to investigate the mechanisms by which the linker histone H1 compacts chromatin. We focused on the human H1.0^45^ because it is an extensively studied linker histone isoform^46^. H1.0 binds near the nucleosome dyad^47^, which increases DNA wrapping into the nucleosome^48^ and promotes chromatin compaction^48–50^. This allows H1.0 to function as a key regulator of transcription by limiting genome accessibility in compact chromatin and suppressing binding of transcription regulatory factors^51,52^. While single-molecule force spectroscopy studies have investigated linker histone-mediated chromatin compaction^18,19^, the impact of linker histones on the energetics of chromatin compaction remains enigmatic.

To determine the impact of H1.0 on the compaction free energy landscape ΔΔG(*ℓ*)_TNuc_, nDFS.C was used initially because it has the highest resolution at small *ℓ*, and H1.0 is anticipated to impact the free energy landscape at small *ℓ* since it compacts chromatin. We carried out FES measurements with tetranucleosomes incubated for 30 minutes at room temperature with increasing concentrations of H1.0 (5, 10, 20, 40, and 80nM) **(Figure S18-S22, Table S2)**. We measured the probability distribution P(*ℓ*) of nDFS.C_TNuc_ at each H1.0 concentration (**Figure 3**). There is a noticeable shift towards smaller *ℓ*, implying that H1.0 converts tetranucleosomes toward more compacted states. We then determined the compaction free energy landscape ΔΔG(*ℓ*)_TNuc_ for increasing H1.0 concentrations and observed the formation of a free energy well over a range of *ℓ* from 5 nm to 25 nm, with the minimum free energy at the 5 nm bin. The formation of this free energy well appears at 10 nM of H1.0 and saturates at about 40 nM where the free energy well is ∼3 k_B_T deep. These results show that H1.0 increases the compaction of tetranucleosomes by converting the 5-20 nm range of the free energy plateau into a free energy well with a minimum at an end-to-end distance *ℓ* around 5 nm.

**Fig 3:**
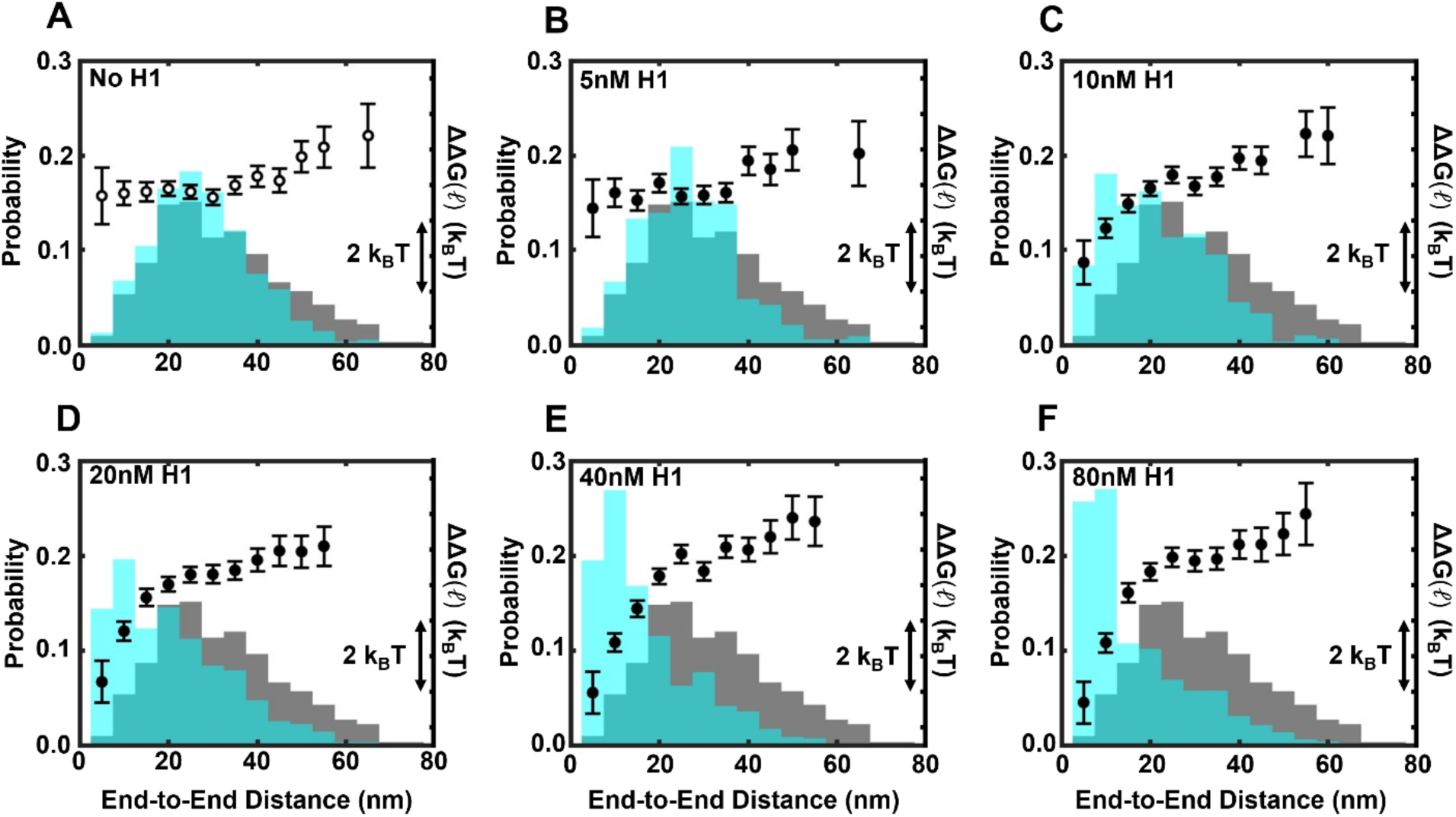
H1 reshapes the tetranucleosome compaction free energy landscape by converting the short end-to-end distance plateau to an energy well. (**A**-**F**) End-to-end distance probability distributions of nDFS.C nanocalipers alone (nDFS.C_only_) and with tetranucleosomes (nDFS.C_TNuc,H1_) are shown in grey and cyan, respectively. The H1.0 concentrations (0, 5, 10, 20, 40, and 80 nM) used in the measurements are listed in the upper left of each plot. Resulting free energy curves are plotted in black closed circles. The error bars of the free energy values were propagated from the uncertainties of the probability measurements, which were estimated by the square root of the number of events in each bin.

While the FES measurements with nDFS.C revealed the impact of H1.0 on tetranucleosome compaction free energies at small end-to-end distances *ℓ*, states above *ℓ* ∼ 60 nm were so rarely populated that free energies could not be determined. Therefore, to measure the compaction free energy landscape ΔΔG(*ℓ*)_TNuc_ over a wider range of end-to-end distances *ℓ*, we repeated the FES measurements with nDFS.A and nDFS.B focusing on the subsaturating and saturating H1.0 concentrations of 10 nM and 40 nM, respectively (**Figure 4A-C, S23-S26, S27A-C, Table S2**). Similar to nDFS.C, both 10nM and 40nM of H1.0 significantly compacted the tetranucleosome where the nDFS.A_TNuc_ and nDFS.B_TNuc_ states occupied lower values of *ℓ*. The compaction free energy landscape ΔΔG(*ℓ*)_TNuc_ was determined for each nDFS (**Figure 4D, S27D**), which extended the free energy measurements out to a value of *ℓ* ∼ 85 nm. We then combined the three measurements of tetranucleosomes with 10 nM and 40nM of H1.0 as was done without H1.0 (**Figure 2E, S27E**) to determine a final compaction free energy landscape ΔΔG(*ℓ*)_TNuc_ with improved free energy resolution and increased range of *ℓ* (**Figure 4E**). Comparing the compaction free energy landscapes ΔΔG(*ℓ*)_TNuc_ with 10 nM and 40 nM of H1.0 (**Figure 4E**, black) to no H1.0 (**Figure 4E**, grey) confirms that as H1.0 is increased, the free energy plateau converts to a free energy well, while for larger end-to-end distances H1.0 has less of an impact on ΔΔG(*ℓ*)_TNuc_. The free energy well has a depth of 3 k_B_T, which increases the probability of these localized states by about 10-fold. Interestingly, this change in free energy is much smaller than the binding free energy of H1.0 to nucleosomes^53^. Overall, these results reveal that H1.0 compacts the tetranucleosome by converting the free energy plateau that already prefers compacted configurations into a free energy well that further localizes the energetically favorable compacted states with end-to-end distances of about 5 nm.

**Fig 4:**
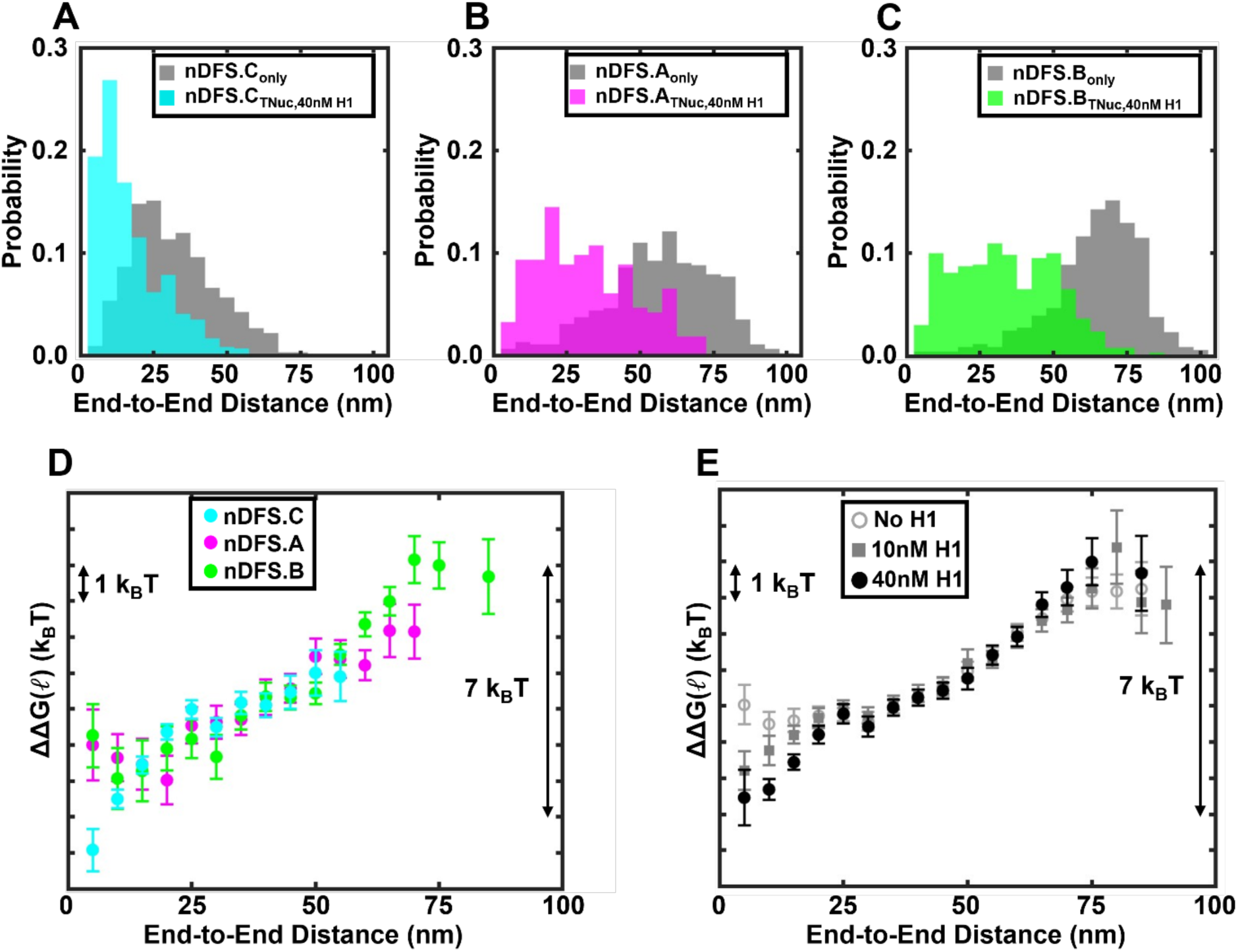
The application of FES with three separate nanocalipers allows for the impact of H1.0 on the tetranucleosome compaction free energy landscape to be probed over a range of 80 nm. (**A**-**C**) End-to-end probability distributions for free nanocalipers are shown in grey for each nanocaliper design (nDFS.C_only_, nDFS.A_only_, nDFS.B_only_). Nanocalipers with tetranucleosomes and 40 nM H1.0: nDFS.C_TNuc,40nM H1_, nDFS.A_TNuc,30nM H1_, nDFS.B_TNuc,40nM H1_, are shown in (A) cyan, (B) magenta, and (C) green, respectively. (C) End-to-end distance tetranucleosome free energy landscapes from each nanocaliper design. The error bars of the free energy values were propagated from the uncertainty of the probability measurements, which were estimated by the square root of the number of events in each bin. (**E**) End-to-end distance tetranucleosome free energy landscapes determined from the weighted average of the three independent free energy measurements using the three different nanocalipers in 1 mM Mg^2+^ with 0 nM H1.0 (open grey circles), 10 nM H1.0 (closed dark grey square), and 40 nM H1.0 (closed black filled circles). The error bars of the free energies were determined as described in the Star Methods section.

### Nucleosome-nucleosome stacking is a second reaction coordinate that is stabilized by H1.0 and disrupted by the DNA nanocaliper

Up to this point, these measurements have considered the full ensemble of tetranucleosome conformations as a function of one parameter, the end-to-end distance *ℓ*. Each value of *ℓ* includes an ensemble of different tetranucleosome conformations. Visual inspection of the TEM images of tetranucleosomes alone and within each of the nDFS devices revealed that each tetranucleosome can be classified as 1-, 2-, 3-, or 4-dot states (**Figure 5A, S28**). The number of dots is related to the number of nucleosome-nucleosome stacking interactions since two stacked nucleosomes appear as a single dot in TEM images, as previously reported^12^. This stacking interaction is mediated through interactions between the histone octamer acidic patch and the H4 N-terminal tail^54,55^, which is critical for the formation of compact chromatin^56,57^.

**Fig 5:**
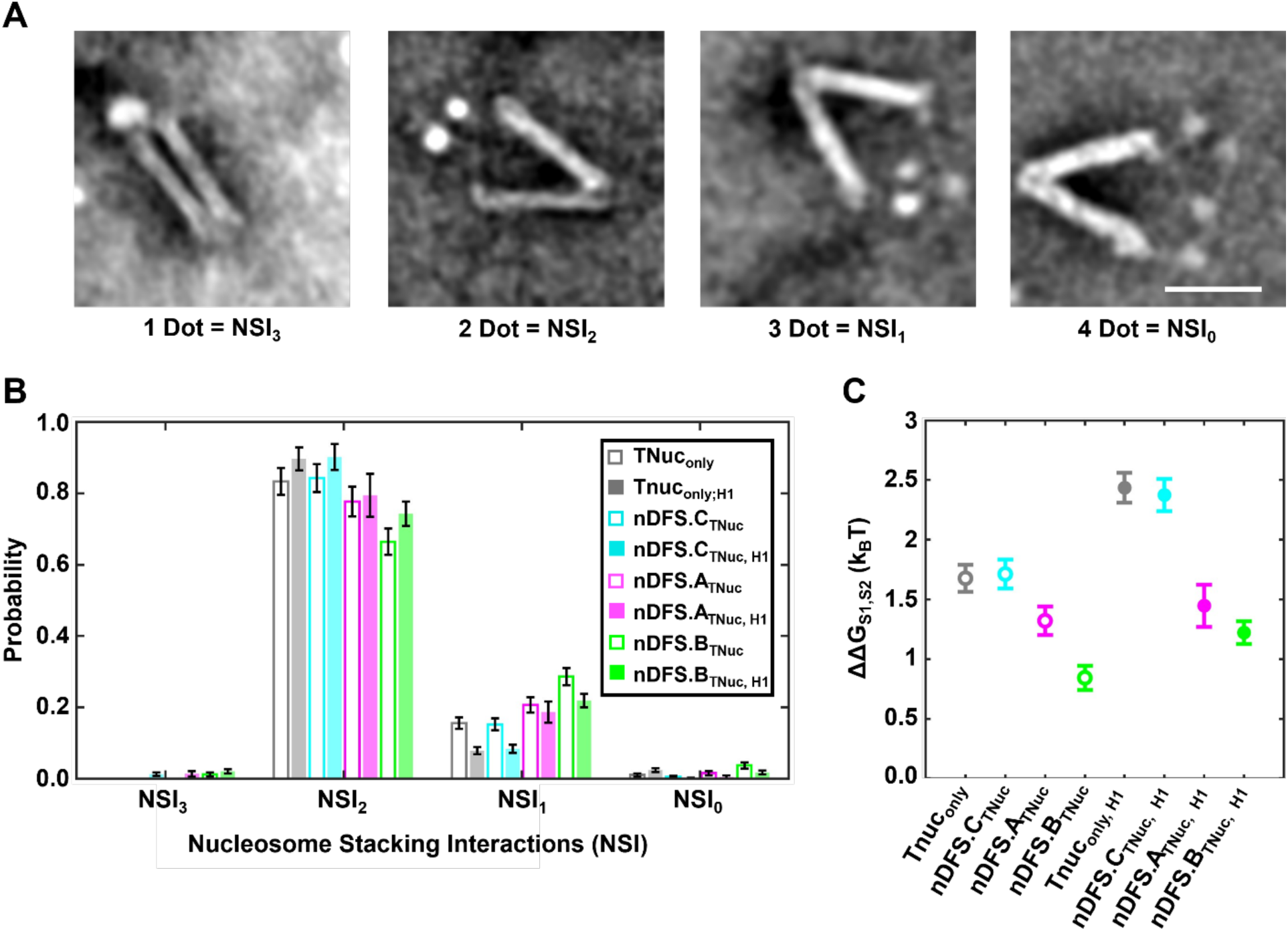
Subclassifying by the number of nucleosome stacking interactions provides insight into the nucleosome stacking free energies. (**A**) Example images of single nanocalipers with a tetranucleosome that has three, two, one, or zero stacking interactions based on the number of observed dots. The scale bar is 50 nm. (**B**) Nucleosome stacking interaction distributions without (open bar plot) and with (closed bar plot) 40 nM H1.0. Tetranucleosomes not incorporated into devices (TNuc_only_) are shown in grey, while tetranucleosomes incorporated into nDFS.C, nDFS.A, and nDFS.B are shown in cyan, magenta, and green, respectively. The error bars are determined from the square root of the number of events in each bin. (**C**) The free energy difference between the state with two nucleosome stacking interactions and the state with 1 nucleosome stacking interaction. The error bars were determined by propagating the uncertainties from the probability measurements in panel (B).

The 4-dot state is consistent with all four nucleosomes being separately visible, implying there are no stacked nucleosomes (**Figure 5A)**. The 3-dot state is consistent with one pair of nucleosomes stacked, which appears as a single dot that is often brighter than the other two single nucleosome dots (**Figure 5A, S28**). Since tetranucleosomes with 30 base pair linker DNA are reported to compact into a 2-start geometry where the n^th^ nucleosome stacks with the n + 2^nd^ nucleosome^12,14^, the 3-dot state likely involves stacking between either the 1^st^ and 3^rd^ nucleosomes or the 2^nd^ and 4^th^ nucleosome, with the other pair being unstacked. The 2-dot state is consistent with two pairs of stacked nucleosomes in the conformation that was solved by x-ray crystallography^14^ and cryo-EM^5^, where the 1^st^ and 3^rd^ nucleosomes stack and the 2^nd^ and 4^th^ nucleosomes stack (**Figure 5A, S28**). Finally, the rarely observed 1-dot state most likely contains three stacking interactions so that there is no space between any of the four nucleosomes (**Figure 5A**). Therefore, we refer to the 4-, 3-, 2-, and 1-dot states with the number of nucleosome stacking interactions, NSI_0_, NSI_1_, NSI_2_, and NSI_3_, respectively.

We classified each tetranucleosome image from four different conditions, without or within an nDFS device and without or with 40nM H1.0 (see Star Methods for details). For each condition, the probability of each stacking state was then determined (**Figure 5B-C**). The NSI_2_ state was dominant for all conditions, with probabilities ranging from 0.67±0.04 to 0.90±0.03 (**Figure 5B, Table S3**). The NSI_1_ state was the second most common, with probabilities ranging from 0.08±0.01 to 0.29±0.02 (**Figure 5B**). Both the NSI_0_ and NSI_3_ states were rare, occurring at probabilities of 0.037±0.009 and 0.240±0.006, respectively, or less (**Figure 5B**, **Table S3**).

The relative probabilities can be converted to free energy differences via the Boltzmann distribution, ΔG = -k_B_T ln [P(NSI_i_) / P(NSI_j_)]. We decided to focus on the NSI_1_ and NSI_2_ states since these were the most probable, providing sufficient sampling. We find that the free energy cost to break one nucleosome stacking interaction, which converts state NSI_2_ to NSI_1_ is 1.6±0.1 k_B_T and 2.4±0.1 k_B_T without and with H1.0, respectively (**Figure 5C, Table S4**). These free energy differences between the NSI_2_ and NSI_1_ states are in line with multiple reports of nucleosome stacking free energy^17,19,34^ and are at the low end of the full range of reported free energies of about 2 to 20 k_B_T^6,18,58,59^. These studies were done over a range of ionic conditions, linker geometries, and nucleosome repeat numbers, which likely explains this order-of-magnitude range of nucleosome stacking free energy values. E.g., a coarse-grained model found the nucleosome stacking free energy to be around 2 k_B_T^21^ when linkers (and thus their mutual repulsion) were included, while the same model determined the stacking between two separate mononucleosomes to be about 15 k_B_T^60^. Interestingly, nDFS.C does not impact these free energy differences (**Figure 5C, Table S4**). However, nDFS.A and nDFS.B reduce the free energy differences between the NSI_1_ and NSI_2_ states, which is expected since both nanocalipers significantly shift the end-to-end distance distributions to larger values (**Figure 2A-C, 4A-C**). Overall, these results reveal that the number of nucleosome stacking interactions is a second reaction coordinate probed by the nDFS experiments, which is complementary to end-to-end distance *ℓ*, and provides additional information about tetranucleosome compaction.

### A multidimensional framework for understanding the energetics of chromatin opening

Now that we have established measurements of the tetranucleosome end-to-end distance free energy landscape ΔΔG(*ℓ*)_TNuc_ and the nucleosome stacking free energy (NSI_1_ vs. NSI_2_), we combined these parameters to provide a multidimensional free energy analysis: ΔΔG(*ℓ*,NSI_i_)_TNuc_. To achieve this, we divided the data into subsets based on the nucleosome stacking state **(Figure S29-S30).** We only included the NSI_1_ and NSI_2_ states because the NSI_0_ and NSI_3_ states did not occur often enough to prepare an energy landscape for these states alone. Importantly, ΔΔG(*ℓ*,NSI_i_)_TNuc_ measurements that compare the NSI_1_ vs. NSI_2_ states are from the same dataset, so differences in free energy can be meaningfully compared.

We first focused on the compaction free energy landscape ΔΔG(*ℓ*,NSI_i_)_TNuc_ in the absence of H1.0 for the two stacking states, NSI_1_ vs. NSI_2_, which we refer to as ΔΔG(*ℓ*,NSI_1_)_TNuc;H1=0nM_ and ΔΔG(*ℓ*,NSI_2_)_TNuc;H1=0nM_, (**Figure 6A, S31**). We find that the NSI_1_ free energy landscape ΔΔG(*ℓ*,NSI_1_)_TNuc;H1=0nM_ is generally higher in free energy as compared to the NSI_2_ free energy landscape ΔΔG(*ℓ*,NSI_2_)_TNuc;H1=0nM_. This is because the probability of the NSI_1_ state is about 5-fold lower than the NSI_2_ state. Since the NSI_2_ state is the most probable, the shape of the NSI_2_ free energy landscape ΔΔG(*ℓ*,NSI_2_)_TNuc;H1=0nM_ (**Figure 6A**, **S31** black circles) is similar to the overall free energy landscape ΔΔG(*ℓ*)_TNuc;H1=0nM_ (**Figure 2E**), where the free energy is constant for *ℓ*<30 nm, then increases for *ℓ*>30 nm. In contrast, the NSI_1_ free energy landscape ΔΔG(*ℓ*,NSI_1_)_TNuc;H1=0nM_ is distinct from the overall free energy landscape ΔΔG(*ℓ*)_TNuc_ in that it has a shallow free energy well for the end-to-end distance range, 30 nm<*ℓ*<45 nm. For *ℓ*<30 nm, the NSI_1_ free energy landscape ΔΔG(*ℓ*,NSI_1_)_TNuc;H1=0nM_ increases, and since the NSI_2_ free energy landscape ΔΔG(*ℓ*,NSI_2_)_TNuc;H1=0nM_ is constant for this range of *ℓ*, the NSI_2_ state is strongly preferred in this range of end-to-end distances *ℓ*. Finally, for *ℓ* >45 nm, the NSI_1_ and NSI_2_ free energy landscapes overlap, i.e. ΔΔG(*ℓ*>45 nm, NSI_1_)_TNuc;H1=0nM_ ∼ ΔΔG(*ℓ*>45 nm, NSI_2_)_TNuc;H1=0nM_. This implies that in the absence of H1.0 the NSI_1_ and NSI_2_ states are equally probable for these large end-to-end distances, and that tetranucleosomes can readily fluctuate between these two states with the available thermal energy.

**Fig 6:**
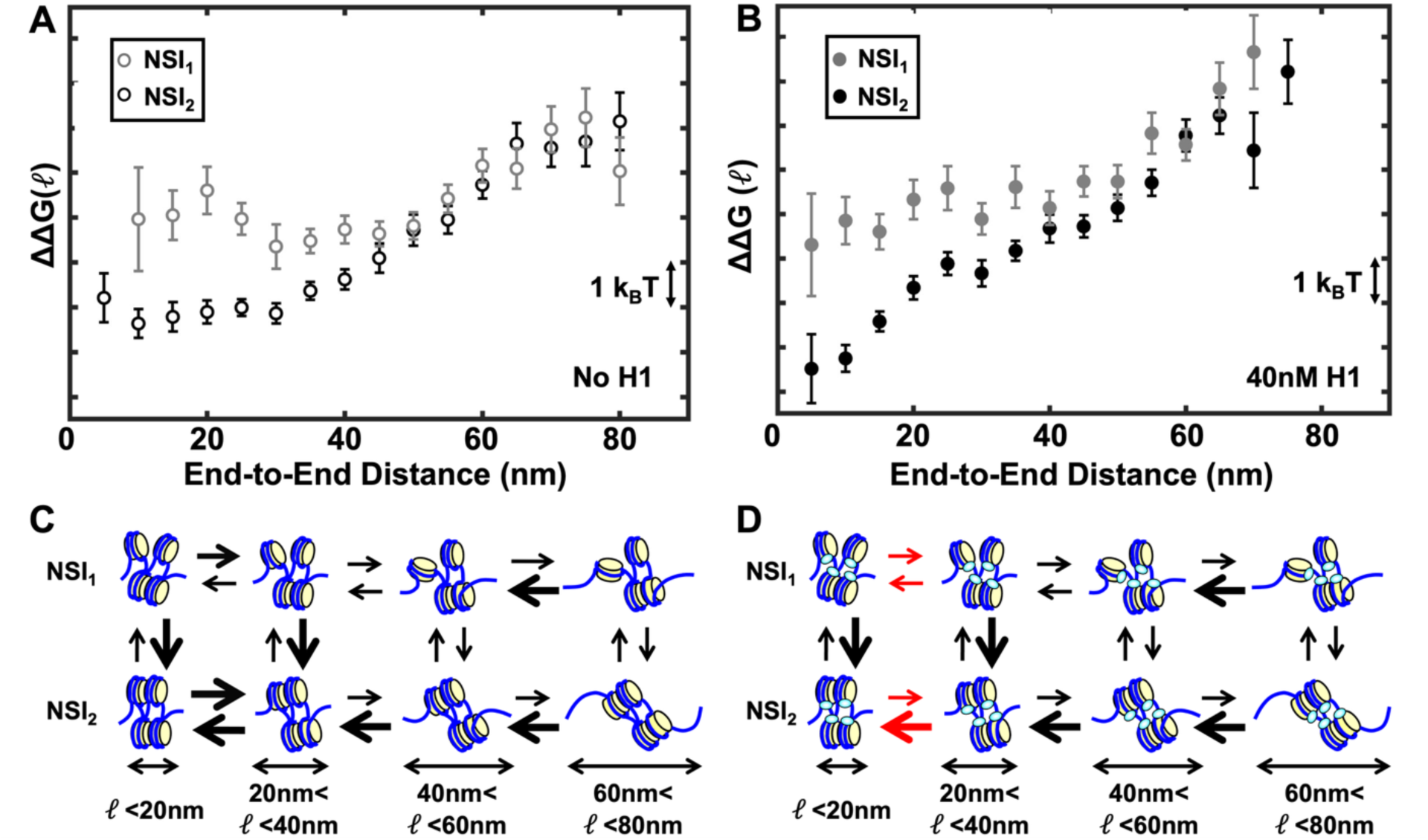
Multidimensional tetranucleosome compaction free energy landscape reveals the energetics for different nucleosome stacking states and how H1.0 reshapes these energy landscapes. (**A**-**B**) End-to-end distance free energy landscapes for partially (NSI_1_, grey) and fully (NSI_2_, black) compacted tetranucleosomes without (A) and with (B) 40 nM H1.0. The error bars of the free energies were determined as described in the Star Methods section. (**C**-**D**) Transition rate model of tetranucleosome compaction based on the multidimensional free energy landscape for tetranucleosomes without (C) and with (D) 40 nM H1.0. The relative size of the arrows for a given transition relates to relative probabilities between states defined by both end-to-end distance ranges and nucleosome stacking interactions. Red arrows in (D) show state transitions that change by H1.0.

We then repeated this analysis with 40 nM H1.0 to determine the impact of this linker histone on the NSI_1_ and NSI_2_ free energy landscapes (**Figure 6B, S32**). As without H1.0, we find that with H1.0 the NSI_1_ free energy landscape ΔΔG(*ℓ*,NSI_1_)_TNuc;H1=40nM_ is generally higher than the NSI_2_ free energy landscape ΔΔG(*ℓ*,NSI_2_)_TNuc;H1=40nM_, which is expected since the NSI_2_ state is about 10-fold more probably than the NSI_1_ state with 40 nm H1.0 (**Figure 5C**). The shape of the NSI_2_ free energy landscape with 40nM H1.0 ΔΔG(*ℓ*,NSI_2_)_TNuc;H1=40nM_ is similar to the overall free energy landscape ΔΔG(*ℓ*)_TNuc_ with 40 nM H1.0 (**Figure 4E**), where H1.0 converts the small end-to-end distance *ℓ* plateau into a free energy well with a minimum at *ℓ* ∼ 5 nm. This is expected since the probability of the NSI_2_ state with 40 nM H1.0 is about 0.9. Interestingly, for *ℓ*<40 nm with 40 nM H1.0, the NSI_1_ free energy landscape ΔΔG(*ℓ*,NSI_1_)_TNuc;H1=40nM_ has a different shape than the NSI_2_ free energy landscape ΔΔG(*ℓ*,NSI_2_)_TNuc;H1=40nM_. In this region, the free energy remains essentially constant, forming a free energy plateau. For *ℓ* > 40 nm, ΔΔG(*ℓ*,NSI_1_)_TNuc;H1=40nM_ and ΔΔG(*ℓ*,NSI_2_)_TNuc;H1=40nM_ overlap, which implies that for this range of end-to-end distances, the NSI_1_ and NSI_2_ states are equally probable and that the tetranucleosome fluctuates between the fully stacked (NSI_2_) and partially stacked (NSI_1_) states.

The impact of H1.0 can be understood by H1 suppressing DNA unwrapping, constraining the tetranucleosome compaction to small end-to-end distances, resulting in the free energy well for the NSI_2_ state. However, when the tetranucleosome partially decompacts into the NSI_1_ state, there is only one stacking interaction. This additional conformational freedom allows for the tetranucleosome to explore a wide range of end-to-end distances with little free energy cost. Overall, these results demonstrate the power of this FES approach, where free energy landscapes of low probability states that are hidden in the overall free energy landscape, but biologically important, can be observed and quantified.

## DISCUSSION

In this work, we utilized a set of DNA origami nanocalipers to implement a methodology, Free Energy Spectroscopy (FES), which experimentally determines multidimensional free energy landscapes of chromatin compaction. This work builds off previous nanocaliper measurements of histone octamer stacking interactions^34^. In particular, this work leverages the ability to tune mechanical properties of DNA origami devices to probe free energies spanning molecular distances over tens of nanometers, and the application of a set of calipers enables quantification of a broader free energy landscape with improved precision. Free energy landscapes provide detailed and mechanistic information about the energetics of chromatin compaction, which currently has only been investigated by computational methods^20–24^. In this study, we determined multidimensional free energy landscapes as a function of both the end-to-end distance *ℓ* and the number of nucleosome stacking interactions NSI. We show that by using multiple nanocalipers with different free energy landscapes, continuous tetranucleosome end-to-end distance free energy landscapes can be determined with a spatial resolution of 5nm over a range of 5 nm to 85 nm and a free energy precision of less than 1 k_B_T over a range of about 7 k_B_T. By including the NSI coordinate, tetranucleosome free energy landscapes with a specific number of nucleosome stacking interactions can also be determined (**Figure 6A-B, Supplementary Figs S31-S32**).

These multidimensional free energy landscapes can be interpreted through the ratio of transition rates between states with different end-to-end distances *ℓ* and nucleosome stacking interactions NSI (**Figure 6C-D**), where higher probability states have larger rates into the state than out of the state. States with one nucleosome stacking interaction NSI_1_ are in the upper row, while states with two nucleosome stacking interactions NSI_2_ are in the lower row. The relative size of the arrows represents the relative state probabilities. For NSI_2_ without H1.0, states with *ℓ*<40 nm have the same free energy and are therefore equally probable. This implies the transition rates between states with different end-to-end distances *ℓ* over this range are the same (**Figure 6C**, *ℓ*<20 nm and 20 nm<*ℓ*<40 nm). For *ℓ*>40 nm, the free energy increases, and the probability decreases, as the end-to-end distance *ℓ* increases. This indicates that the transition rates to smaller end-to-end distances *ℓ* are greater than to larger *ℓ*. In contrast, for NSI_1_ without H1.0, the free energy has a minimum between 30 nm<*ℓ*<50 nm, implying that these are the highest probability states. This indicates that the transition rate to states with *ℓ* between 30 nm and 50 nm is larger than the rates out of this range of *ℓ*. Importantly, the lower free energy of NSI_2_ relative to NSI_1_ for *ℓ*<50 nm, implies that NSI_2_ is more probable for this range of *ℓ* and that transitions from NSI_2_ to NSI_1_ are energetically unfavorable. For *ℓ*>50 nm, the free energies are the same, implying that NSI_1_ and NSI_2_ are equally probable and that the rates between states are essentially the same.

By including H1.0, FES reveals how H1.0 impacts the tetranucleosome compaction free energy landscape of both the NSI_2_ and NSI_1_ states. The transition rates that are impacted by H1.0 are highlighted by the red arrows in **Figure 6D**. H1.0 converts the NSI_2_ landscape with a free energy plateau with end-to-end distances *ℓ* of less than about 30 nm to a free energy well with a minimum at the end-to-end distance *ℓ* of about 5 nm. This implies that H1.0 compacts fully stacked (NSI_2_) tetranucleosomes by localizing the lowest free energy (most probable) states from a wide range of end-to-end distances *ℓ* to a small range of end-to-end distances around 5 nm. Interestingly, this free energy well is about 3 k_B_T, which is significantly smaller than the linker histone binding free energy to nucleosomes^53^, suggesting H1.0 is highly dynamic while bound to tetranucleosomes and that H1.0 does not need to dissociate for the tetranucleosome to decompact. For NSI_1_, the free energy well within the range of 30 nm<*ℓ*<50 nm, is converted to a constant free energy plateau for *ℓ*<50 nm. This implies that H1.0 compacts partially stacked (NSI_1_) tetranucleosomes by converting a free energy well around large end-to-end distances to a plateau where the tetranucleosome can explore a wide range of different compacted states. In addition to these changes in the NSI_1_ and NSI_2_ free energy landscapes, the H1.0 reduces the free energy of the NSI_2_ relative to NSI_1,_ increasing the probability of the NSI_2_ state (**Figure 5**), which further contributes to tetranucleosome compaction. This demonstrates how deconvolving with FES the NSI_1_ and NSI_2_ free energy landscapes, we have detected and quantified three separate mechanisms by which H1.0 compacts tetranucleosomes.

Interestingly, while H1.0 increases the free energy difference between fully and partially stacked tetranucleosomes by about 1 k_B_T, it significantly biases both the fully and partially stacked states to smaller end-to-end distances. This is consistent with H1.0 binding near the nucleosome dyad and is not located to directly impact nucleosome stacking, which is mediated by H4 N-terminal tail interactions with the histone octamer acidic patch^54,55^. H1.0 decreases nucleosome unwrapping fluctuations^51,52^, which is consistent with H1.0 having a limited range of *ℓ* that impacts the free energy landscape and having the largest impact for small values of *ℓ*. We conclude that changes in the NSI_1_ and NSI_2_ free energy landscapes are largely due to an increase in the nucleosome unwrapping free energies. These additional conclusions further highlight how multidimensional FES provides new mechanistic insight into how H1.0 compacts tetranucleosomes by directly disentangling its impact on nucleosome stacking and unwrapping.

FES is well-positioned to be used to investigate the numerous factors that control chromatin compaction and regulate eukaryotic genome transcription, repair, and replication. The factors include histone post-translational modifications, histone variants, linker histone isoforms, nucleosome spacing, histone modifying complexes, ATP-dependent chromatin remodeling complexes, chromatin architectural factors, and transcription factors. Furthermore, FES has the potential to investigate the structural dynamics of other biomolecular complexes, such as RNA folding and its regulation by RNA-binding proteins. Finally, given the ease of obtaining the DNA scaffold and DNA oligonucleotides used to fold nanocalipers, and the general availability of TEM, FES is positioned to become a broadly used method for investigating factors that regulate chromatin states within both euchromatin and heterochromatin.

## LIMITATIONS OF THE STUDY

FES requires a control to determine if a chromatin regulator directly influences the nanocaliper conformational distribution. While we found that the linker histone H1.0 did not impact the nanocaliper distributions, other chromatin regulators could have a direct impact on the nanocalipers. This does not imply the FES cannot be used for such regulators; however, the nanocaliper conformational distribution would need to be determined for each concentration of the chromatin regulator.

While this method determines the relative probabilities of a wide range of conformations, it does not provide kinetic information about the time to transition between different conformational states. Alternative methods will be required to investigate transition kinetics. This set of nanocalipers enabled studies of compacted and partially compacted nucleosomes. Extending these studies to fully decompacted tetranucleosomes or larger chromatin molecules may require nanocaliper design alterations to probe larger end to end distances.

## RESOURCE AVAILABILITY

### Lead Contact

Further information and requests for resources and reagents should be directed and will be fulfilled by the Lead Contact, Michael G. Poirier.

### Materials Availability

Unique and stable reagents generated in this study are available upon request

### Data and Code Availability

- TEM images have been deposited at Mendeley data and are publicly available as of the day of publication.
- The MatLab code for converting the final free energy measurements from three separate probability distributions will be made available on GitHub following publication. The code used for running the molecular dynamics simulations will also be made available on GitHub following publication.
- Any additional information required to reanalyze the data reported in this paper is available from the lead contact upon request.

## ACKNOWLEDGEMENTS

The authors acknowledge the feedback and insights provided by the Poirier, Castro, Bundschuh, and Zhang Lab members. This work was funded by the U.S. National Science Foundation, Division of Molecular and Cellular Biosciences (#2411725 to M.G.P. and C.E.C.; #2042362 to B.Z.), and the National Institutes of Health (R35 GM139564 to M.G.P.; R35 GM133580 to B.Z). We acknowledge resources from the Campus Microscopy and Imaging Facility (RRID:SCR_025078) and the OSU Comprehensive Cancer Center Microscopy Shared Resource, The Ohio State University. This facility is supported in part by grant P30 CA016058, National Cancer Institute, Bethesda, MD. Research reported in this publication was supported by the Office of the Director, National Institutes of Health under award S10 OD023582. This work used Bridges-2 at PSC through allocation BIO240299 from the Advanced Cyberinfrastructure Coordination Ecosystem: Services & Support (ACCESS) program, which is supported by U.S. National Science Foundation grants #2138259, #2138286, #2138307, #2137603, and #2138296.

## AUTHOR CONTRIBUTIONS

K.B., M.G.P, C.E.C., and R.B. conceived the experiments. K.B. and M.G.P. wrote the original draft. C.E.C., R.B., I.R, and B.Z. edited the manuscript. K.B., with help from Y.W., performed the TEM, image analysis, and data analysis. K.B., with help from R.W.C., prepared all samples. I.R. carried out the molecular dynamics simulations with feedback from B.Z. M.G.P., C.E.C., R.B., and B.Z. supervised and obtained funding for the studies.

## STAR METHODS

### Preparation of nDFS device

Methods for designing and fabrication of the nDFS devices (nDFS.A, nDFS.B, and nDFS.C) are described previously and the design of the nDFS was previously shared on the public DNA nanostructure design repository nanobase.org (https://nanobase.org/structures/248).^1,2^ nDFS.C is nDFS.C35 in the previous report^1^. The list of staples for all nDFS designs used was previously reported^1^. For folding of the structure, 20 nM of an M13MP18 8064nt ssDNA scaffold was combined with 200 nM of each staple ssDNA in a ddH_2_O solution containing 5 mM Tris, 5 mM NaCl, 1 mM EDTA and 18 mM MgCl_2_, at pH 8.0. The sequence of each staple was determined using the computer assisted design software caDNAno^3^. The folding reaction was carried out in a thermocycler (Bio-Rad, Hercules, CA) with a 15-minute melting step at 70 °C followed by an annealing step from 63 °C-57 °C at a cooling rate of 1 °C every 3 hours.

### Purification of nDFS device

The nDFS devices were purified using previously described methods.^1^ Briefly, the solution containing newly folded nDFS structures was mixed with equal volume PEG buffer (15% PEG MW8000, 200 mM NaCl and 100 mM Tris) and placed into a standard tabletop centrifuge at 20,000g for 30 minutes. Then, the supernatant was removed, and the purified structures were resuspended into a buffer containing 100 mM HEPES pH 8.0, 1 mM MgCl_2_, and 200 mM NaCl. The concentration of the purified nanocalipers was measured via absorption measurements at 260 nm from a NanoDrop, and the structures were diluted in the same buffer to 10 nM for future experiments.

### Nucleosome template DNA preparation

Tandem repeats of the mp1^4^ nucleosome positioning sequence were cloned into pUC19 and amplified through polymerase chain reaction (PCR) using *Pfu* DNA polymerase which contains an endonuclease proofreading enzyme. The amplified DNA was purified using an anion exchange chromatography column (Gen-Pak™ Fax – Waters Column) using high-performance liquid chromatography (abbreviated as HPLC, Agilent). Purified DNA was buffer-exchanged into 0.5xTE (5mM Tris-HCl pH 8.0, 0.5mM EDTA) by Amicon^®^ Ultra Centrifugal Filters (Mereck Millipore). 147 bp long buffer DNA used for tetranucleosome reconstruction was amplified out of the ampicillin gene of the pUC19 plasmid, and HPLC purified in the same manner as described above.

### Histone octamer preparation

Human histones (H2AK119C, H2B, H3C110A, and H4) were obtained from The Histone Source at Colorado State University. Histone octamer was refolded using a 1.3:1 molecular ratio of H2AK119C and H2B to H3C110A and H4 through double dialysis using unfolding buffer (7 M guanidine hydrochloride, 20 mM Tris-HCl pH 7.5, 10 nM DTT) to refolding buffer (2 M NaCl, 10 mM Tris-HCl pH 7.5, 1 mM EDTA pH 8.0, 5 mM 2-Mercaptoethanol).^5,6^ For gel imaging following native polyacrylamide gel electrophoresis (PAGE) of tetranucleosomes, the H2AK119C was Cy5 labeled at K119C using the maleimide-thiol reaction that was quenched with 10 mM DTT. The refolded histone octamer was purified using Fast Protein Liquid Chromatography (abbreviated FPLC, Cytiva) using a Superdex 200 (Cytiva). The fractions were analyzed by 16% SDS-PAGE gel. Selected fractions were concentrated to a volume of ∼100 μl using 30 K Amicon^®^ Ultra Centrifugal Filters (Merck Millipore).

### Preparation of NeutrAvidin bound tetranucleosomes

Tetranucleosome arrays were reconstituted and purified as previously described^7^. Template tetranucleosome DNA containing a repeat of four mp1^4^ nucleosome positioning sequences and 30 bp linker DNA was combined with buffer DNA in a 1:3 mass ratio with Cy5-labeled histone octamer. A mass ratio of total DNA to histone octamer at 1:0.8 was then prepared in a 50 μl volume in the reconstitution buffer (0.5xTE pH 8.0, 1 mM benzamidine hydrochloride, and 2 M NaCl). Tetranucleosomes were reconstituted by double dialysis where the 50 μl sample was loaded into a small dialysis chamber that was placed in a dialysis chamber containing 80 mL of the reconstitution buffer. This was dialyzed against 4 L of 0.5x TE pH 8.0, 1 mM benzamidine hydrochloride for 6 hours and then against new buffer overnight. The tetranucleosome sample was purified on a 5 to 35% (w/v) sucrose gradient in 0.5x TE at 41000 rpm for 16 hours in a SW41 rotor at 4 °C. Fractions were analyzed by PAGE, and then selected fractions based on the image from a typhoon scanner (Cytiva) were pooled, buffer exchanged and concentrated with 30K amicon filters. The purity of the tetranucleosome sample was then confirmed by 4% PAGE **(Figure S33).**

NeutrAvidin was attached to both biotin-labeled 5 prime ends of the tetranucleosomes by incubating with 40-fold molar excess of NeutraAvidin for 30 minutes at room temperature. Excess Neutravidin was purified away from NeutrAvidin bound tetranucleosomes on a sucrose gradient as described above for the first sucrose gradient purification. The purified Neutravidin labeled tetranucleosomes were analyzed by 4% PAGE **(Figure S34).** Glycerol was added to the sample at a 20% final concentration, which was then aliquoted, flash frozen, and stored at -80 °C

### Optimization of glutaraldehyde crosslinking

Deposition of nucleosome arrays onto TEM grids disrupts nucleosomes and chromatin compaction, so mild glutaraldehyde (GA) crosslinking is used to maintain nucleosomes and chromatin compaction^8–10^. To determine the optimal GA crosslinking conditions for FES, conditions tested were based on previous reports^11^.

To determine the minimal concentration of GA that prevents nucleosome disruption without inducing nucleosome stacking or tetranucleosome aggregation in decompacting conditions, free tetranucleosomes were diluted to 2.5 nM in decompacting buffer conditions (2 mM NaCl, 100 mM HEPES pH 8.0) and crosslinked for 1 minute with 0.025%, 0.05%, 0.1%, 0.2%, 0.5%, 1%, 2%, and 4% glutaraldehyde. **(Supplementary Fig S4, Table S5)**. Tetranucleosomes were then deposited on TEM grids and imaged. Above 0.5% GA, TEM imaging revealed that multiple tetranucleosomes were crosslinked together, while at 0.5% GA and below, individual tetranucleosomes were largely observed. The dot number distributions were then determined for 0.5% GA and below to quantify nucleosome stability and tetranucleosome compaction. At 0.5% GA and below, only 4 dot and 3 dots were observed. However, below 0.2% GA, the fraction of 3 dots increased. This is likely due to the disruption of single nucleosomes since lowering the GA concentrations should not increase nucleosome stacking nor increase the NSI_1_ state relative to the NSI_0_ state. Instead, below 0.2% GA, nucleosomes were disrupted, increasing arrays with only 3 nucleosomes. These TEM measurements under decompacting conditions revealed that GA concentrations between 0.2% and 0.5% stabilize single nucleosomes without significantly inducing tetranucleosome compaction or multimerization.

To determine the range of GA concentrations that maintained tetranucleosome compaction under compacting conditions (200 mM NaCl, 1 mM MgCl_2_, 100 mM HEPES pH 8.0), tetranucleosomes were diluted to 2.5 nM and then GA crosslinked for 1 minute. GA concentrations of 0.02%, 0.04%, 0.06%, 0.08%, 0.1%, 0.2%, and 0.5% were tested. **(Supplementary Figure S5, Table S6)**. 2.5 nM of nanocalipers were included to better replicate experimental conditions. NeutrAvidin was not included, so tetranucleosomes could not incorporate into the nanocalipers. At GA concentrations below 0.08% GA, a significant fraction of NSI_0_ states were observed, which were not expected with 1 mM MgCl_2_, while a similar fraction of NSI_2_ and NSI_1_ states was observed for 0.08% GA and above. Therefore, 0.2% GA is the minimal concentration that is needed to prevent deposition-induced nucleosome disruption and tetranucleosome decompaction, without inducing tetranucleosome compaction or multimerization.

### Preparation of the nDFS device with a tetranucleosome and linker histone H1

Tetranucleosomes and nDFS devices were combined at equal molar ratio (5 nM TNucs and 5 nM nDFS Devices) in a buffer containing 10 mM HEPES pH 8.0, 200 mM NaCl, and 0.25 mM or 1 mM MgCl_2_ and then incubated at room temperature for 2 hours. For linker histone experiments, after the initial 2 hour incubation, H1.0 (NEB) was added at a final concentration of 5, 10, 20, 40, and 80 nM and incubated at room temperature for 30 minutes. The sample was then crosslinked with 0.2% GA for 1 minute by quenching the crosslinking with 100 mM final concentration of Tris-HCl pH 8.0. Each sample was then deposited on a TEM grid for imaging.

### Transmission electron microscopy grid preparation, imaging, and analysis

TEM imaging samples were prepared as previously described^1^. Briefly, 10 μl of the final tetranucleosome sample was deposited on Formvar-coated copper TEM grids, stabilized with evaporated carbon film (Ted Pella; Redding, CA). The sample was incubated on the grid for 10 min. The tetranucleosome sample was then wicked off the grid with filter paper. The sample was stained by applying 6 μl of 2% uranyl formate (SPI, West Chester, PA) twice for 1 and 5 s, respectively. The staining solution was wicked off after each incubation with filter paper. TEM imaging was carried out at the OSU Campus Microscopy and Imaging Facility on an FEI Tecnai G2 Spirit TEM at an acceleration voltage of 80 kV at a magnification of 75,000×.

Single molecules (nDFS_only_, TNuc_only_, and nDFS_TNuc_) were manually selected from the original TEM images at each H1 and Mg^2+^ condition using ImageJ. At 0.25 mM Mg^2+^, the particles were sufficiently spread out that both nDFS_TNuc_ and nDFS_only_ (i.e. nanocalipers with no TNucs incorporated) particles were collected from the same grid for nDFS.A and nDFS.B. For the other nDFS_only_ measurements, separate grids without tetranucleosomes were prepared for particle imaging. Example TEM images are shown in the **Supplementary Material**. Selected 100x100 pixel images containing a single device were filtered using the built-in bandpass filter in ImageJ. This filtered small structures up to 3 pixels (half of the nucleosome diameter) and large structures down to 90 pixels (twice the hinge arm length). After filtering, only images that met the selection criteria (see below) were retained for further analysis. Representative 10x10 image galleries showing 100 randomly selected particles for each condition are shown in the **Supplementary Materials**. The selection criteria include verifying that the hinges are folded properly and deposited on the surface in a clear side view orientation, and contain one tetranucleosome that is bound to both hinge arms. We measured the incorporation efficiency for each device **(Figure S3, Table S7).** After selection, end-to-end distances were measured manually in ImageJ by placing points at the inner edge of each hinge arm.

CSV files from ImageJ containing the location of each point were used to make end-to-end probability distributions, which were in turn used to determine free energy landscapes. Analysis and measurements were carried out in MATLAB. The stacking state of each tetranucleosome was classified manually by visual inspection of each particle. Each dataset was split according to the stacking state and then used to determine the probability distributions and free energy landscapes of each stacking state.

### Free energy measurements and weighted average computation

Free energy landscape measurements for nDFS_only_ and nDFS_TNuc_ were computed using ΔG = -k_B_T ln (P(*ℓ*) / P_max_) where P(*ℓ*) is the probability for a given *ℓ*, while P_max_ is the probability for the value of *ℓ*. Since we report a relative free energy landscape, the offset resulting from the fact that the most likely end-to-end distance is different for nDFS_only_ and nDFS_TNuc_ is not included.

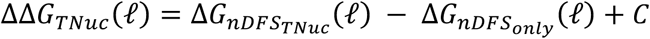

Where C is a constant that can be removed. The free energy landscape of the tetranucleosome was therefore calculated using

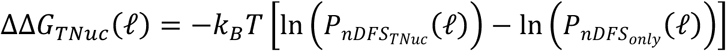

The uncertainty in the probability was computed using the square root of the number of bin counts. Standard uncertainty propagation of the formula above leads to

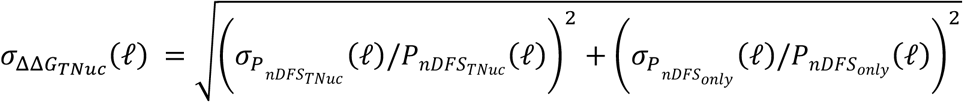

For the H1.0 measurements, we found that H1.0 does not significantly alter the nDFS.C_only_ probability distribution (**Figure S6B**). Therefore, the nDFS.C_only_ probability distribution in the absence of H1.0 was used for determining the compaction free energy landscape ΔΔG(*ℓ*)_TNuc_ at each H1.0 concentration.

Since C varies for each device, there is a constant free energy offset between the free energy measurements from each of the devices. To compute this offset, the following loss function was minimized:

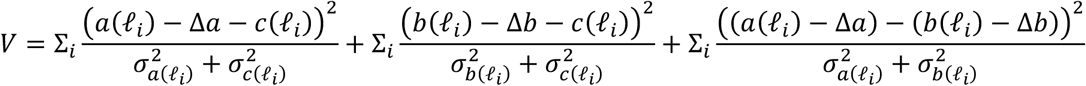

Here *a*(*ℓ*_*i*_) is the measured free energy from nDFS.A at end-to-end distance *ℓ*_*i*_. The purpose of this loss function was to find two constant offsets Δ*a* and Δ*b* that shifted the free energy curves for the measurements from nDFS.A and nDFS.B such that it minimized the difference between datapoints from each nanocaliper measurement, but also considered the uncertainty of each measurement. This offset was determined before the weighted average measurement was computed.

The weighted average mean for each end-to-end distance < ΔΔG(*ℓ*) > from the three independent measurements from each nanodevice was computed with

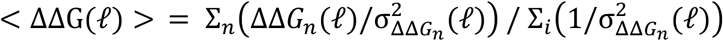

where n represents each nanocaliper design (e.g. nDFS.C). This is weighted by the inverse of the uncertainty for each nanocaliper at each end-to-end distance.

To compute the weighted uncertainty, we added in quadrature the standard deviation (δ_std_) from the three independent measurements at each bin center and the weighted uncertainty 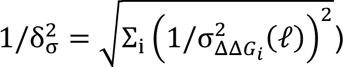 as determined by the individual uncertainties from each of the three measurements:

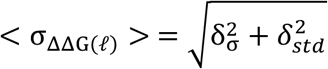

We repeated this same process to determine the weighted end-to-end free energy landscape for NSI_2_ and NSI_1,_ where the free energies were calculated from the probability distributions for tetranuclesomes in each stacking state.

To compute the ensemble free energy difference between NSI_1_ and NSI_2_, we used ΔG_NSI(2-1)_ = -k_B_T ln [P(NSI_1_) / P(NSI_2_)]. Where P(NSI_i_) is the probability that a tetranucleosome has i nucleosome stacking interactions. We used the normalized square root of the counts for each NSI state as the uncertainty σ_NSI_i__ and propagated the uncertainty so that

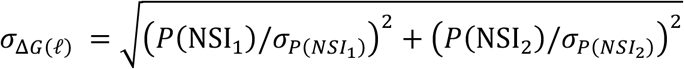

### Coarse grained modeling of free energy landscape

Two-dimensional umbrella sampling simulations were performed. In addition to the end-to-end distance, *ℓ*, which served as a collective variable and facilitates direct comparison with experimental data, a second variable, D_p_, was introduced. This additional coordinate enables sampling of multiple chromatin unfolding pathways was identified in the previous study^12^. Specifically, D_13_ and D_24_ represented the distances between the centers of mass (COMs) of nucleosomes 1 and 3, and nucleosomes 2 and 4, respectively. Their difference, defined as D_p_ = D_13_ − D_24_, served as the second collective variable.

Applying an explicit bias along the D_p_ axis allowed the simulations to selectively probe distinct unfolding mechanisms: one in which the interaction between nucleosomes 1 and 3 was disrupted (D_p_ > 0), another where the contact between nucleosomes 2 and 4 broke (D_p_ < 0), and a symmetric pathway where both interactions were lost simultaneously (D_p_ ∼ 0). This targeted control over pathway selection enhances the overlap between adjacent umbrella windows and improves the convergence of the resulting free energy profile.

A total of 66 GPU-accelerated MD simulations using OpenMM^13^ as the molecular mechanics software along with CG model implementations from OpenABC^14^ were performed. The DNA was prepared with the same sequence as that used by our experimental setup. Each nucleosome in the tetranucleosome had 11 DNA base pairs centered at the dyad and non-disordered regions of the histone core treated as a rigid body to prevent nucleosome sliding and histone dissociation. Electrostatic and solvent interactions, represented by the Debye-Huckel potential, were calculated at 300 K with a 200 mM ionic strength solution. The 66 MD simulations were run in NVT systems with Langevin dynamics via OpenMM’s Langevin Middle Integrator for a total of 2 μs at a 10 fs timestep with collective variables and structures reported every 10 ps. Each simulation had a unique set of CV centers maintained by harmonic restraints. CV centers were selected such that each had a spacing of ∼7.5nm along both CV coordinates, within a set of bounding lines which ensured that at *ℓ* = 2nm, |D_p_| ≤ 5nm and at *ℓ* = 80nm, |D_p_| ≤ 25nm. Prior to running the simulations, each system had a short warm-up simulation where CV restraints were slowly applied over 1 ns. In addition to CV restraints, the warm-up simulation also included restraints to maintain the desired D_13_ and D_24_ distances; these additional restraints are released after the warm-up steps have been completed.

FastMBAR^15^ was used to solve the MBAR equations and “de-bias” each completed simulation, that is, calculate the energy of each state in the full statistical ensemble as generated by the combined set of simulations. With the de-biased energies, the free energy landscape of the tetra-nucleosome with respect to the set of collective variables was computed.

## SUPPLEMENTARY INFORMATION

### Supplementary Figures

**Figure S1:**
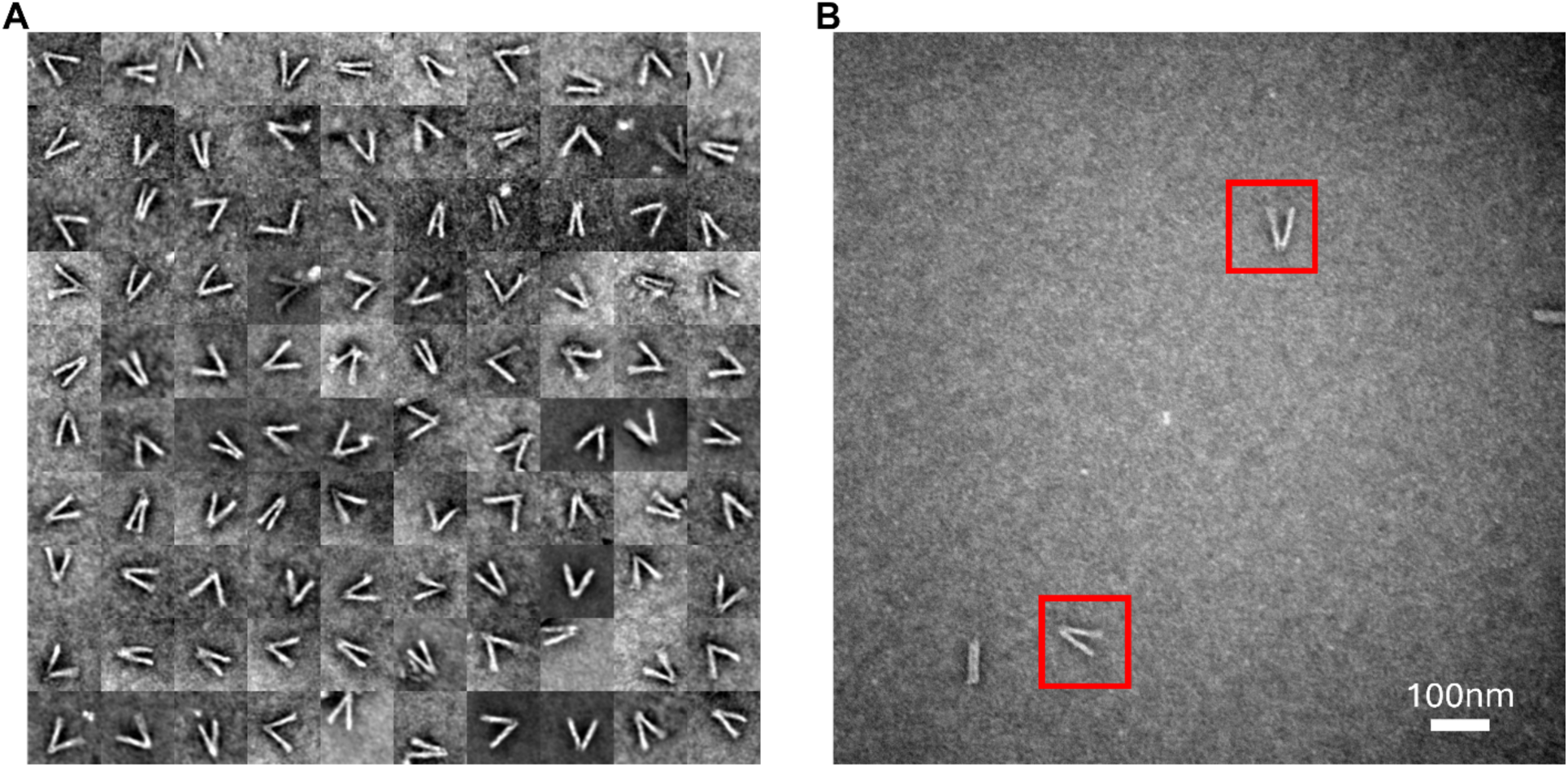
(**A**) Image gallery of 100 nDFS.C_only_ nanocalipers selected at random from the full dataset of 635 particles. (**B**) Example TEM image with example nDFS.C nanocalipers in red boxes.

**Figure S2:**
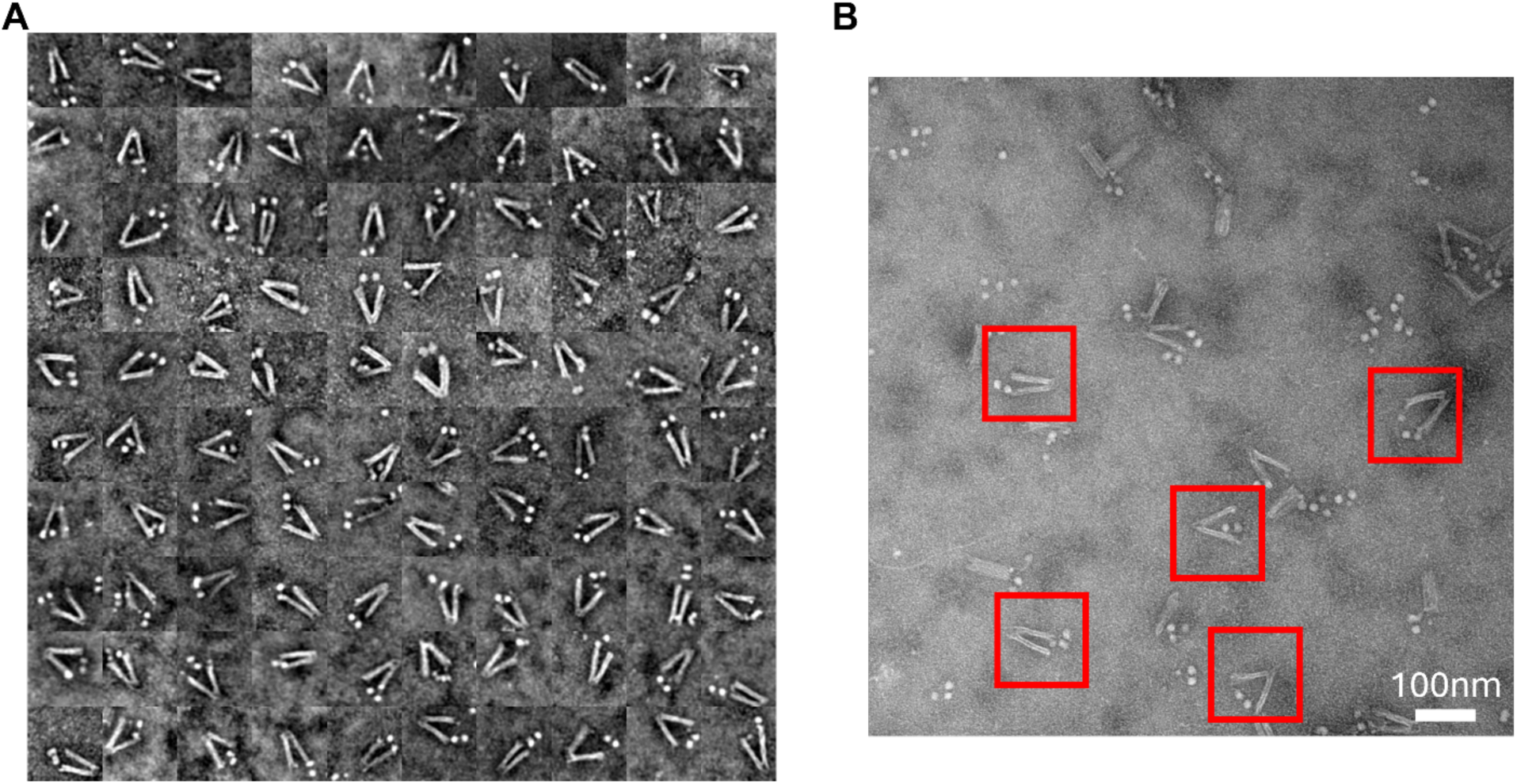
(**A**) Image gallery of 100 nDFS.C_TNuc_ nanocalipers that contain properly integrated tetranucleosomes. They were selected at random from the full dataset of 546 particles. (**B**) Example TEM image with example nDFS.C_TNuc_ in red boxes.

**Figure S3:**
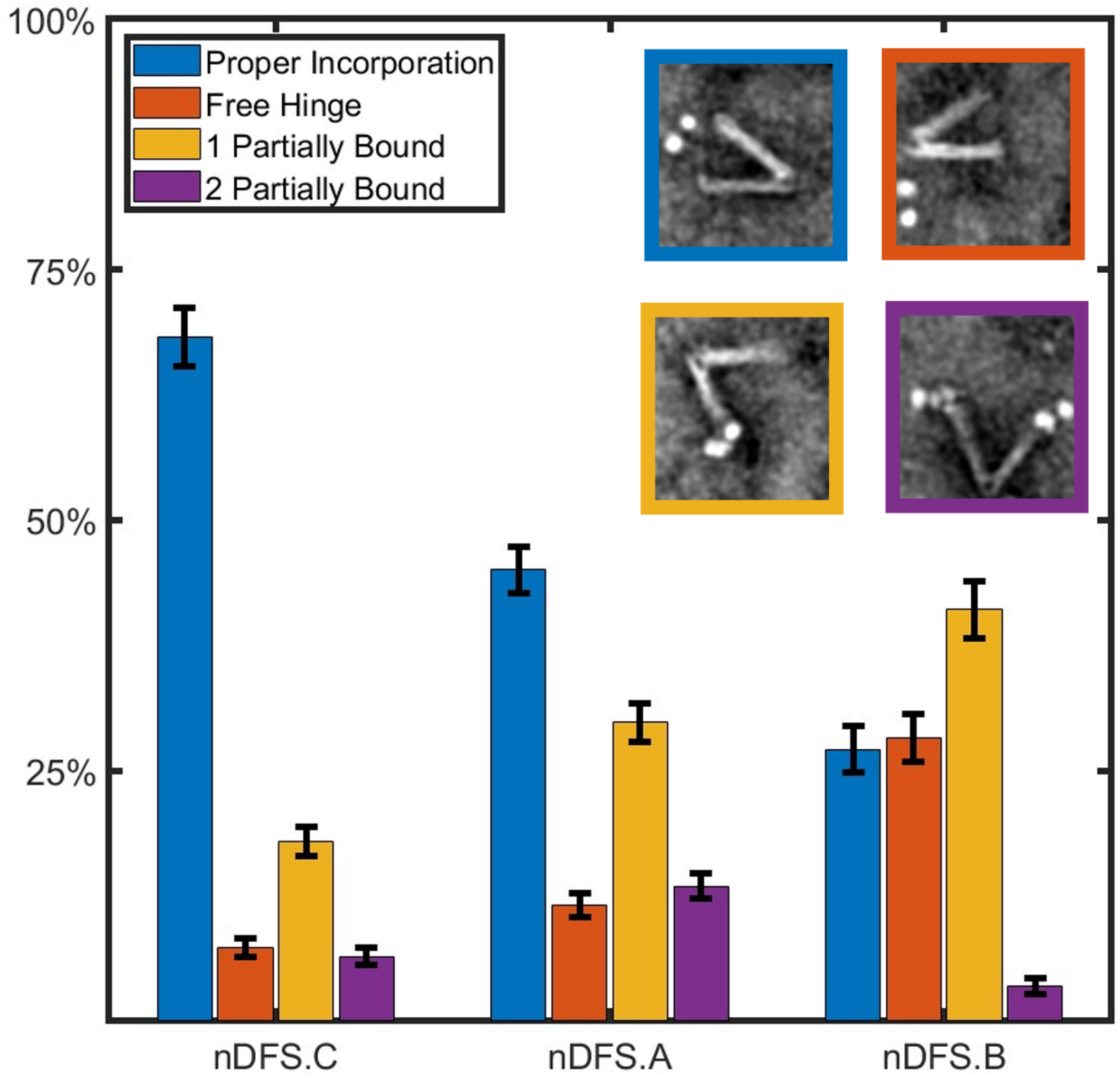
Incorporation efficiency of tetranucleosomes into nanodevices. Bar plot of the probability for each incorporation state for each nanodevice (nDFS.A-C). Blue is the probability of a nanocalilper with a tetranucleosome properly incorporated. Orange is the probability of a nanocaliper without a tetranucleosome. Yellow is the probability of a nanocaliper with one tetranucleosome bound to only one arm. Purple is the probability of a nanocaliper with two tetranucleosomes where each is bound to separate arms. Insets are examples of images for each incorporation state.

**Figure S4:**
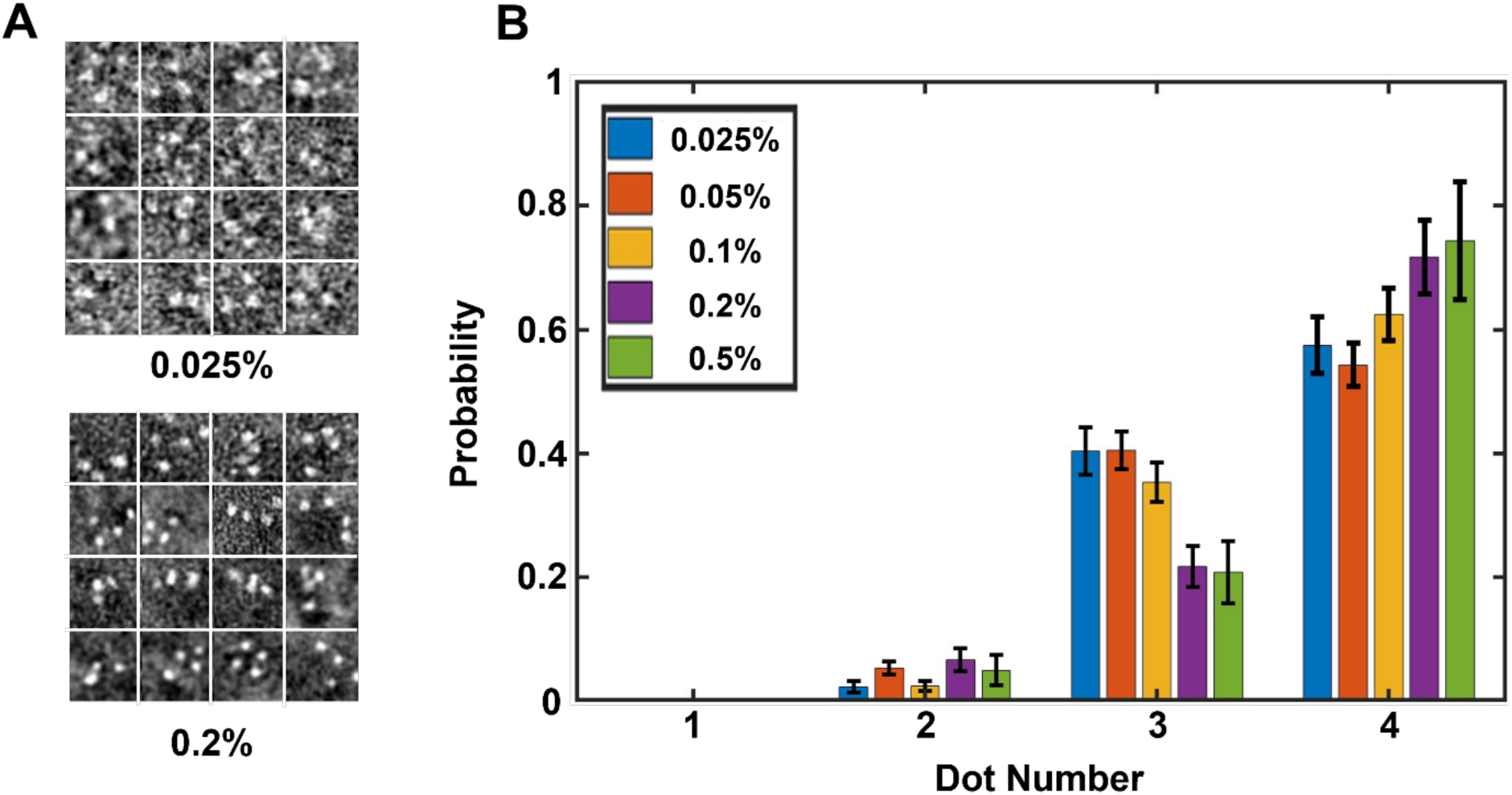
Optimization of glutaraldehyde crosslinking of tetranucleosome in decompacting conditions. **(A)** Galleries of TEM images of tetranucleosomes in a buffer (2 mM NaCl and 100 mM HEPES pH 8.0) that maintains chromatin in an open, decompacted state. The tetranucleosomes were crosslinked with either 0.025 and 0.2% glutaraldehyde (GA) for 1 minute. (**B**) The dot number distribution of individual tetranucleosomes that are crosslinked with 0.025-0.5% GA for 1 minute in decompacting conditions. Above 0.5% GA, the tetranucleosomes multimerized, so a dot number distribution could not be determined.

**Figure S5:**
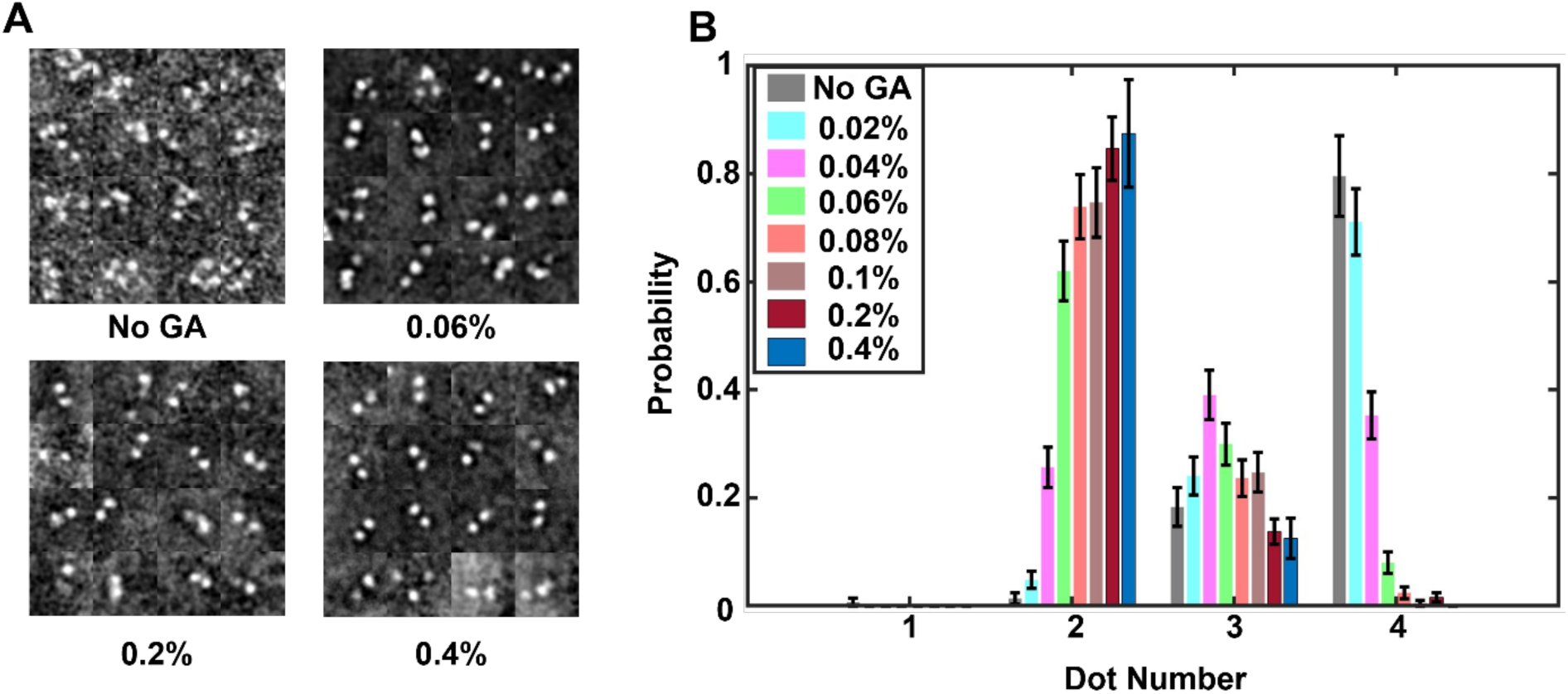
Optimization of glutaraldehyde crosslinking of tetranucleosome in compacting conditions. **(A)** Galleries of TEM images of tetranucleosomes in a buffer (1 mM MgCl_2_, 200 mM NaCl and 100 mM HEPES pH 8.0) that maintains chromatin in a compacted state. The tetranucleosomes were crosslinked with either 0, 0.06%, 0.2%, and 0.4% GA for 1 minute. (**B**) The dot number distribution of individual tetranucleosomes that are crosslinked with 0.025-0.4% GA for 1 minute in compacting conditions.

**Figure S6:**
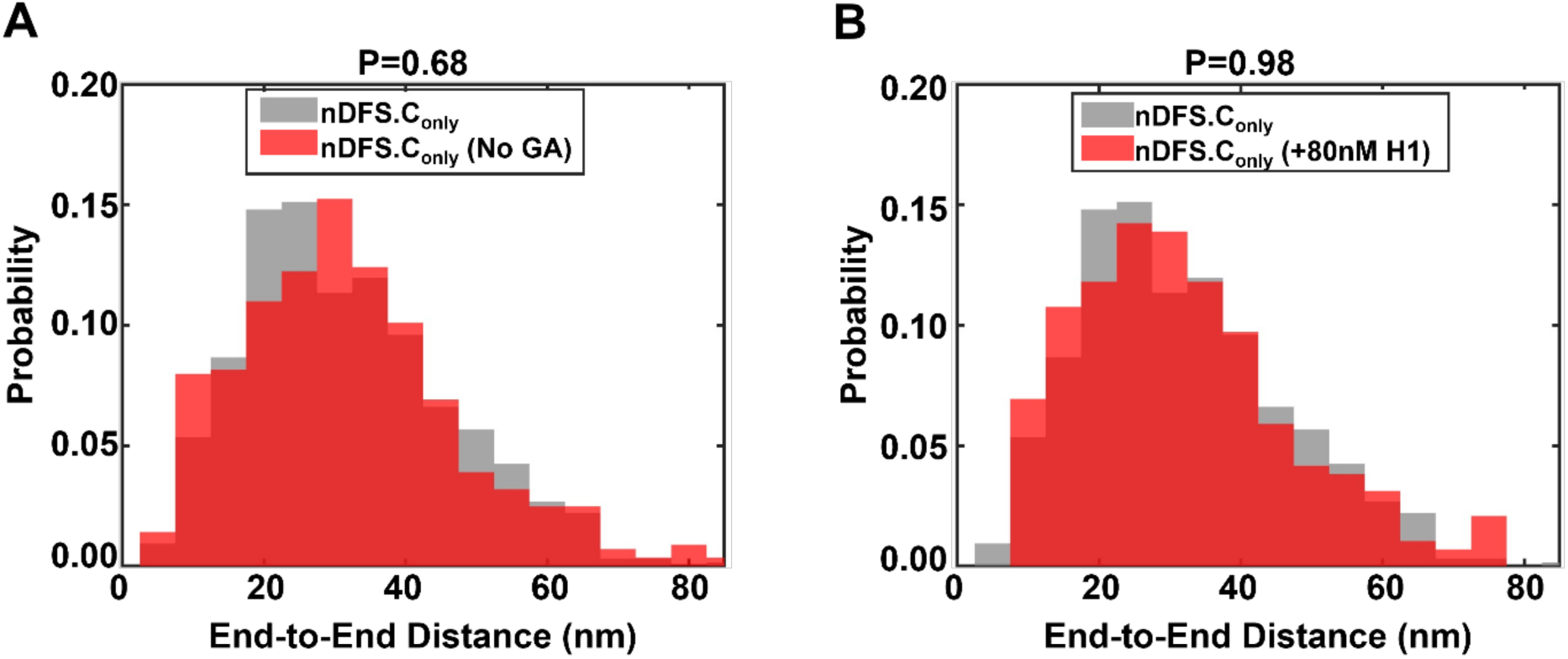
Glutaraldehyde crosslinking and H1.0 do not significantly change the nDFS.C end-to-end distance probability distribution. (**A**) End-to-end distance probability distribution of nDFS.C_only_ without GA crosslinking (grey) and with GA crosslinking at 0.2% for 1 minute (red). (**B**) End-to-end distance probability distribution of nDFS.C without H1.0 (grey) and with 80 nM H1 (red). The KS test was used to compare the distributions. The P values for each are shown in the title of the plot. Each has P>0.05, indicating that crosslinking with 0.2% GA for 1 minute, and the highest H1 concentration used in this study (80 nM) do not significantly impact the nDFS.C_only_ distribution.

**Figure S7:**
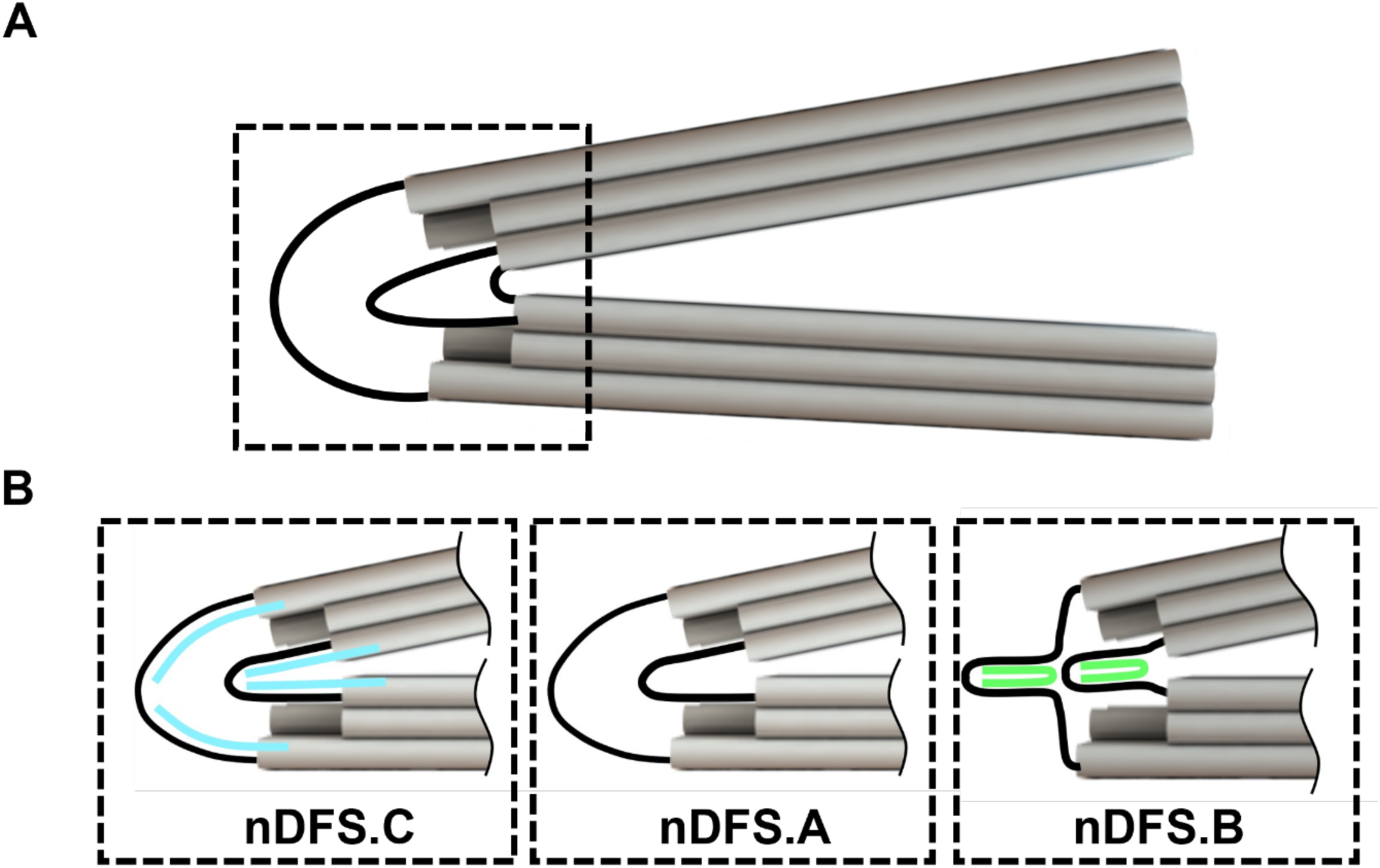
Schematic of vertex modifications which modulate mean nanocaliper angle. (A) Complete schematic representation of the the nanocaliper with grey cylinders representing double helixies of DNA and black curves representing single stranded connections between the nanocaliper arms. The two long connections are 70 base pairs long and the short connection is 2 base pairs long. (B) Schematic of vertex modifications for nDFS.C, nDFS.A, and nDFS.B – the order corresponding to an increasing mean end-to-end distance. The top and bottom nanocaliper arm in nDFS.A is connected fully by single stranded DNA. For the other nanocaliper designs, additional single stranded oligos (cyan for nDFS.C and green for nDFS.B) which are complementary to DNA near the vertex are added to modulate the nanocaliper distribution.

**Figure S8:**
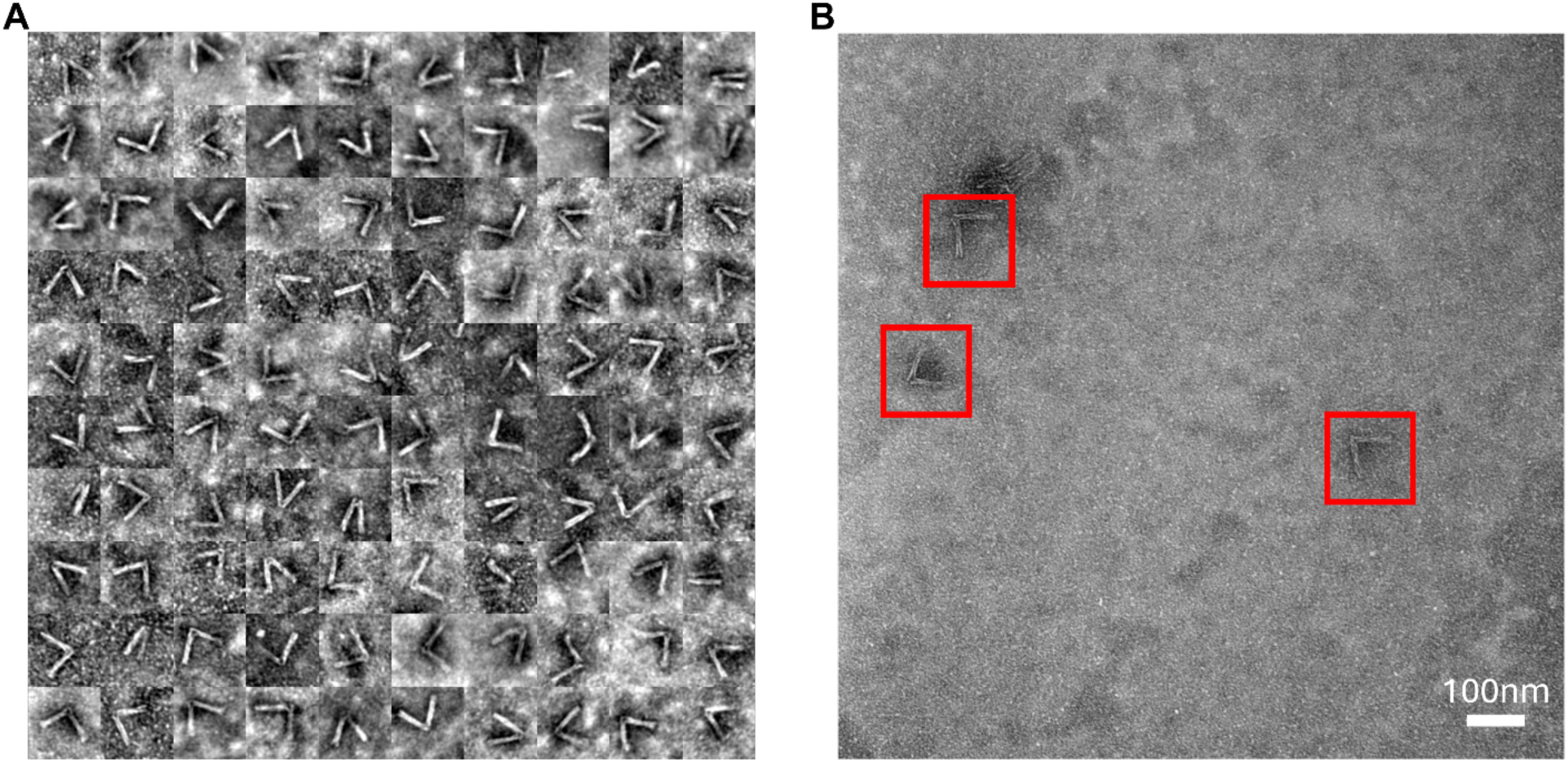
(**A**) Image gallery of 100 nDFS.A nanocalipers selected at random from the full dataset of 545 particles. (**B**) Example TEM image with example nanocalipers in red boxes.

**Figure S9:**
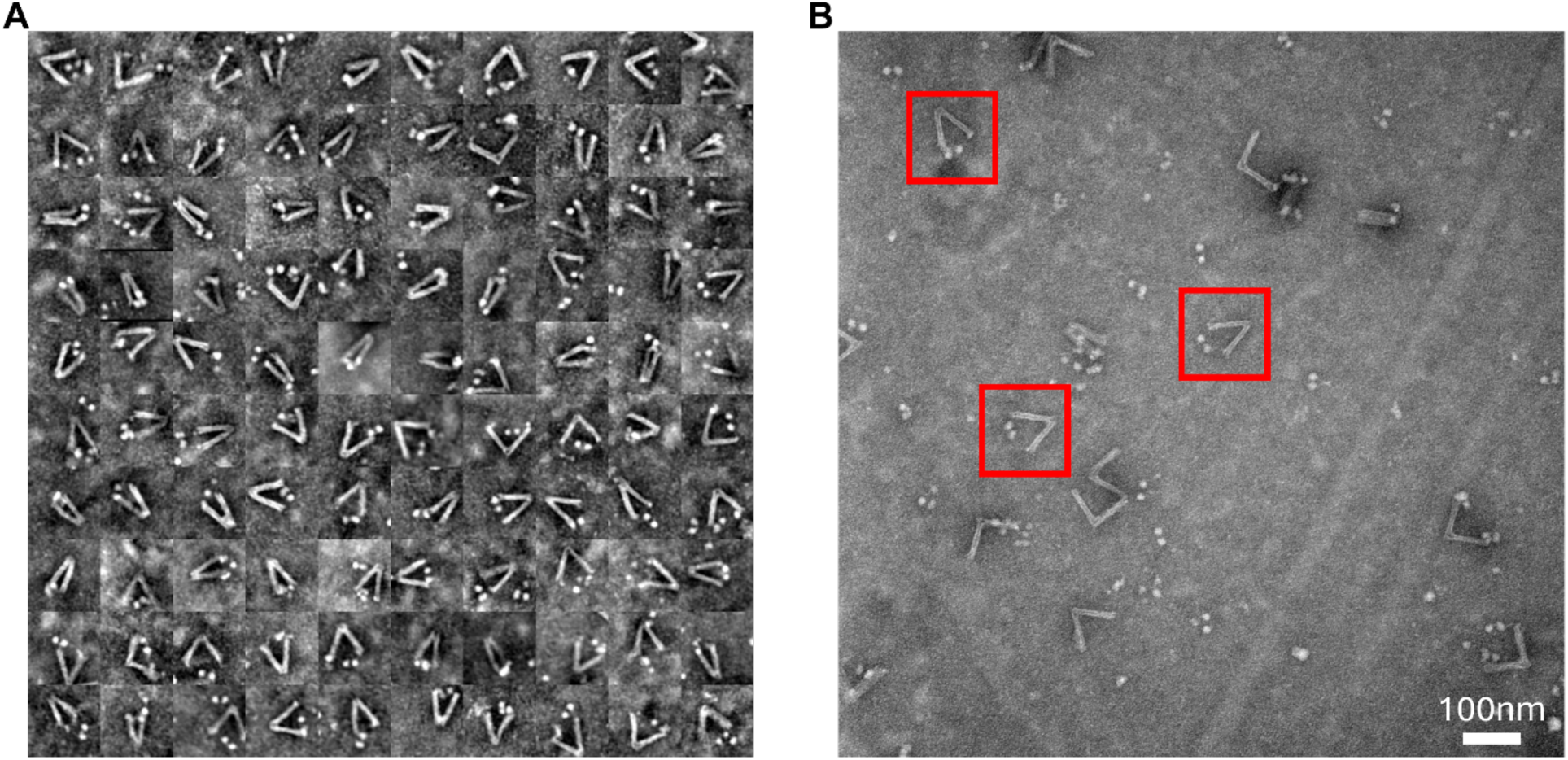
(**A**) Image gallery of 100 nDFS.A_TNuc_ nanocalipers that contain properly integrated tetranucleosomes. They were selected at random from the full dataset of 439 particles. (**B**) Example TEM image with example nDFS.A_TNuc_ in red boxes.

**Figure S10:**
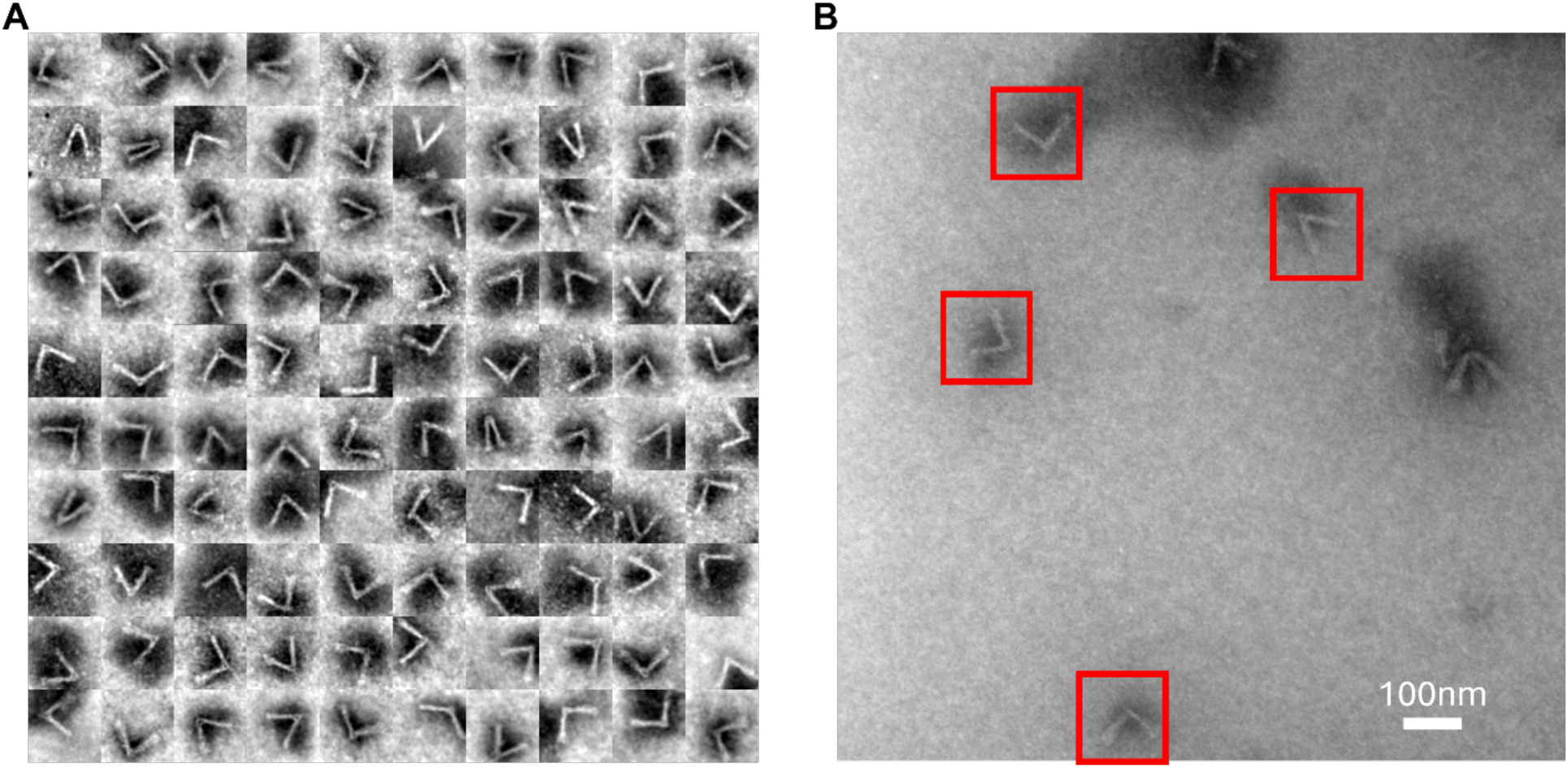
(**A**) Image gallery of 100 nDFS.B nanocalipers selected at random from the full dataset of 687 particles. (**B**) Example TEM image with example nanocalipers in red boxes.

**Figure S11:**
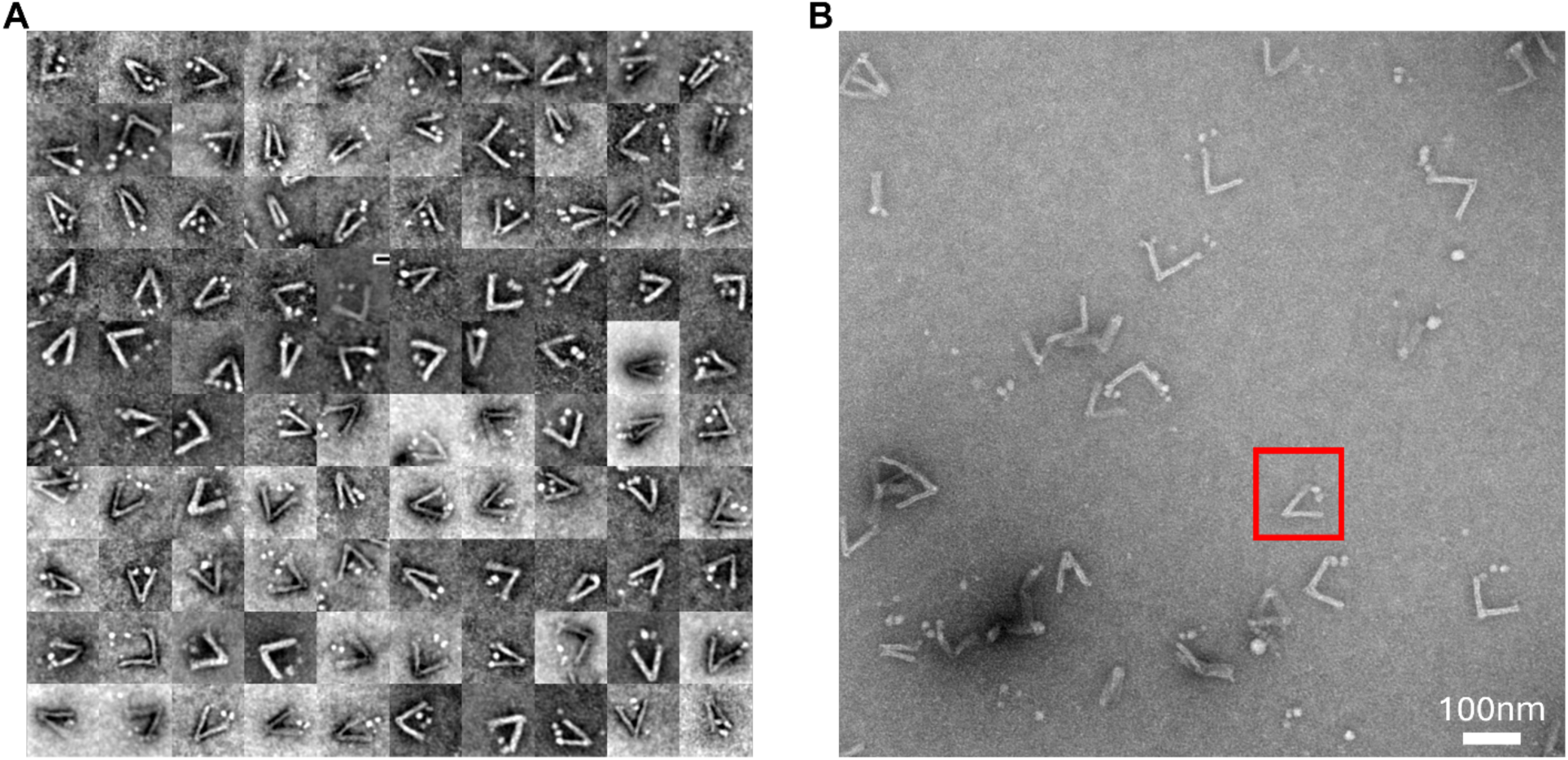
(**A**) Image gallery of 100 nDFS.B_TNuc_ nanocalipers that contain properly integrated tetranucleosomes. They were selected at random from the full dataset of 486 particles. (**B**) Example TEM image with example nDFS.B_TNuc_ in red boxes.

**Figure S12:**
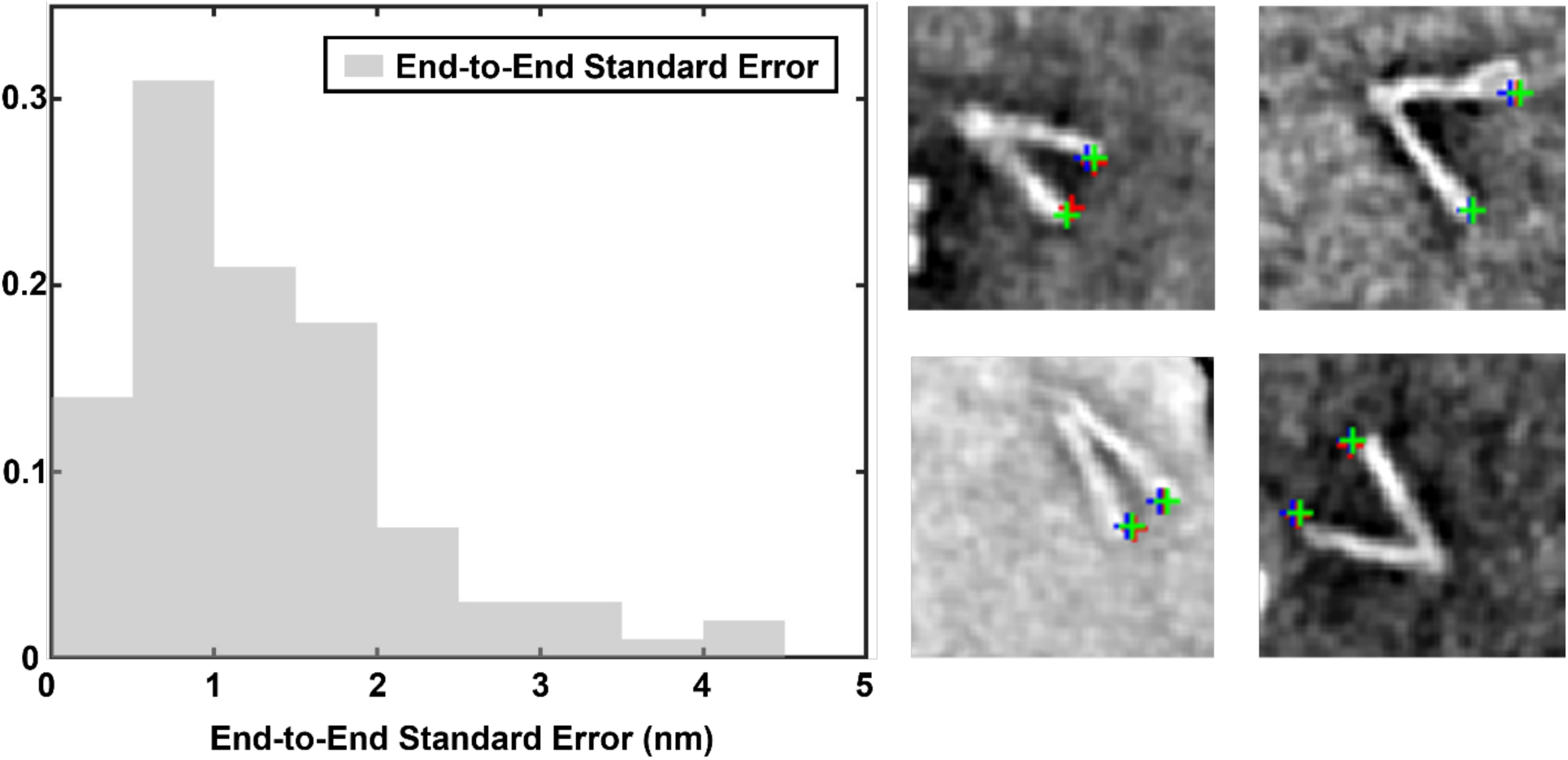
Uncertainty of the nanocaliper end-to-end distance measurement. The end-to-end distance of 100 nDFS.C_only_ was measured manually three times to estimate the uncertainty. The standard error in each measurement was then determined for each individual particle from the three separate measurements. (**A**) The distribution of standard deviations for the end-to-end distance measurement of 100 nanocalipers. The mean standard deviation was 1.3 nm. (**B**) Example images of four nanocalipers showing three different measurements with blue, red, and green crosses. In some cases, all three measurements cannot be seen because the measurements overlap.

**Figure S13:**
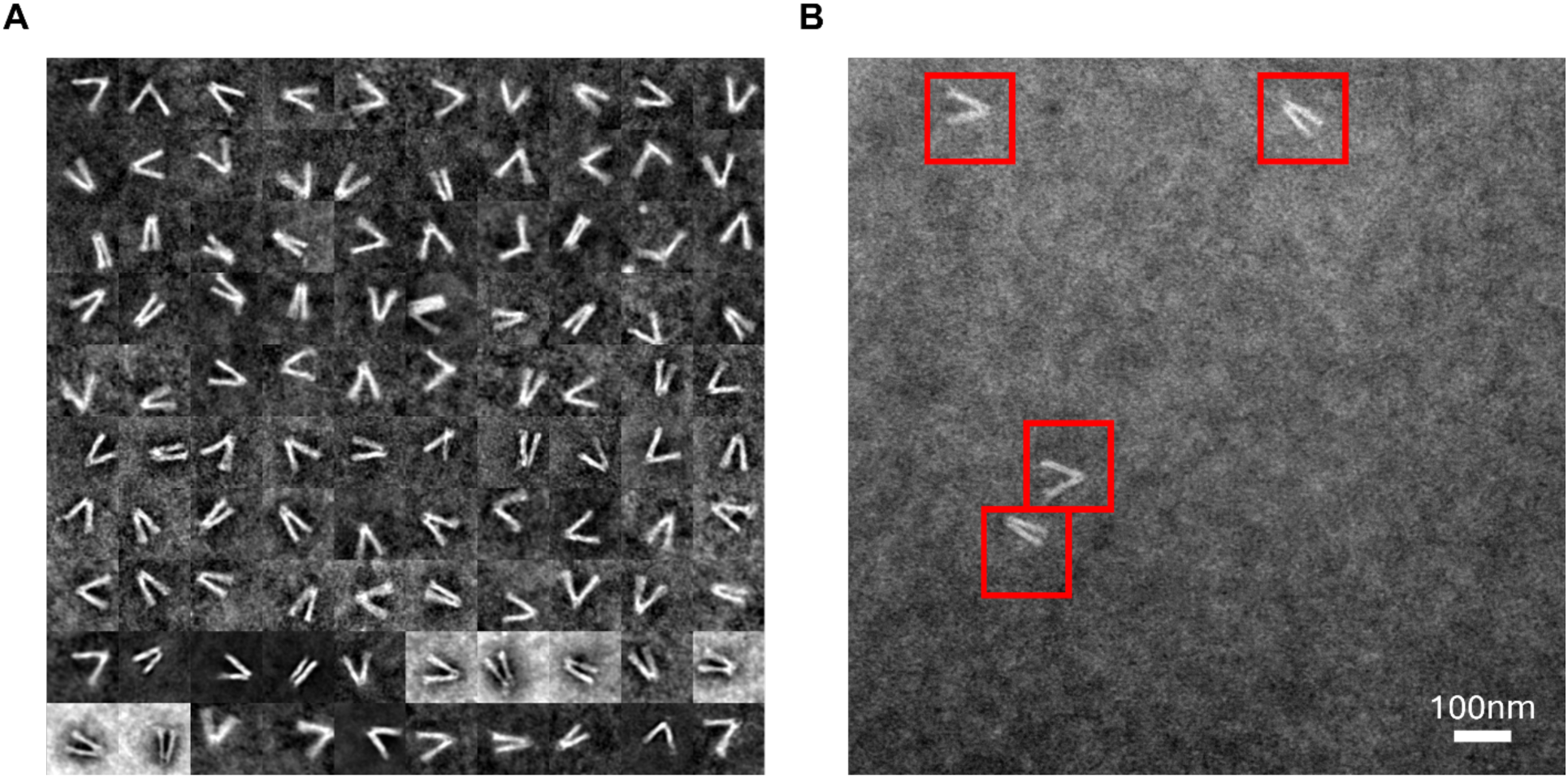
(A) Image gallery of 100 nDFS.C nanocalipers in 0.25 mM Mg^2+^ selected at random from the full dataset of 408 particles. (B) Example TEM image with example nanocalipers in red boxes.

**Figure S14:**
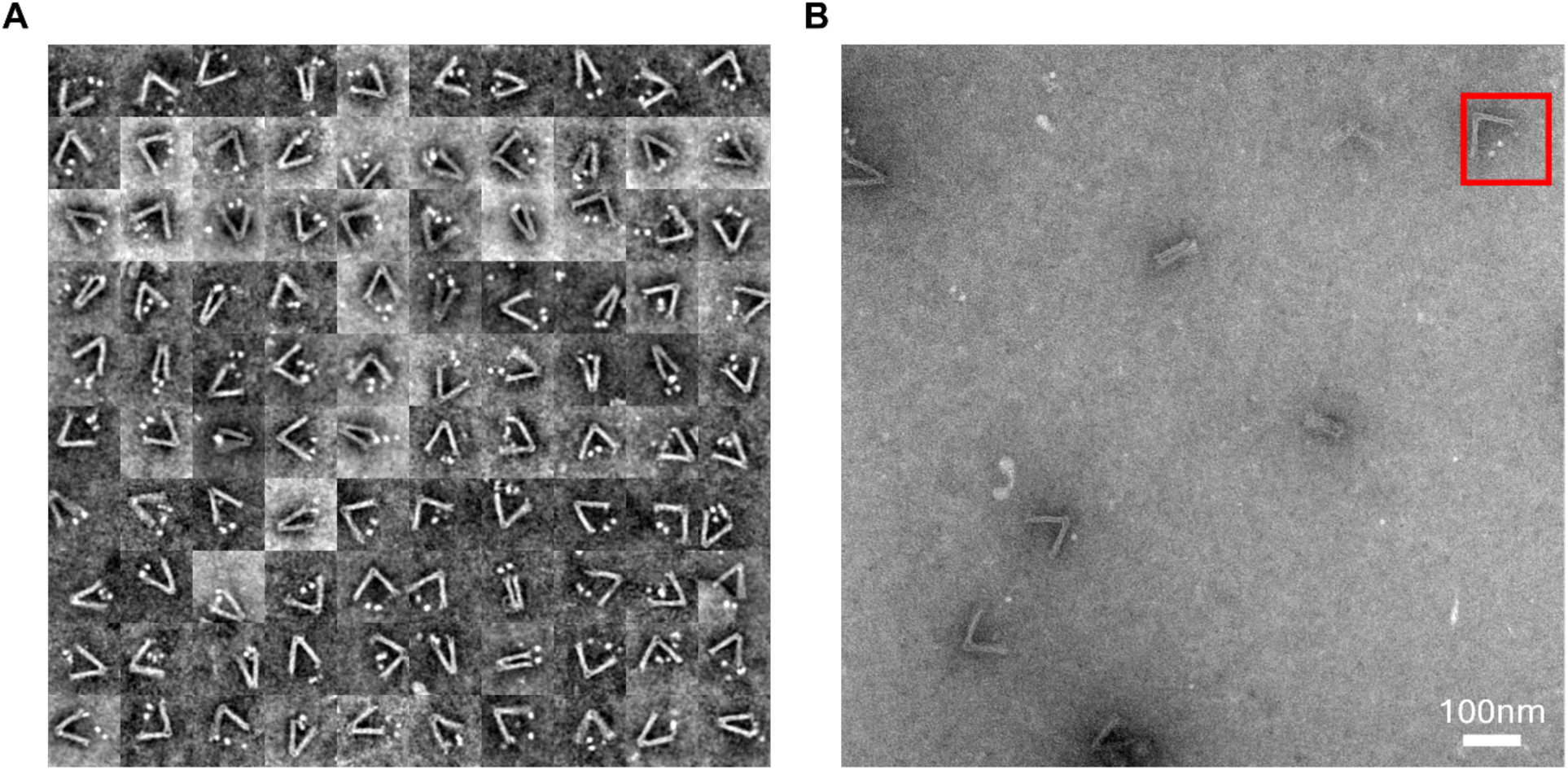
(**A**) Image gallery of 100 nDFS.A_TNuc_ nanocalipers in 0.25 mM Mg^2+^ that contain properly integrated tetranucleosomes. They were selected at random from the full dataset of 246 particles. (**B**) Example TEM image with example nDFS.A_TNuc_ in the red box.

**Figure S15:**
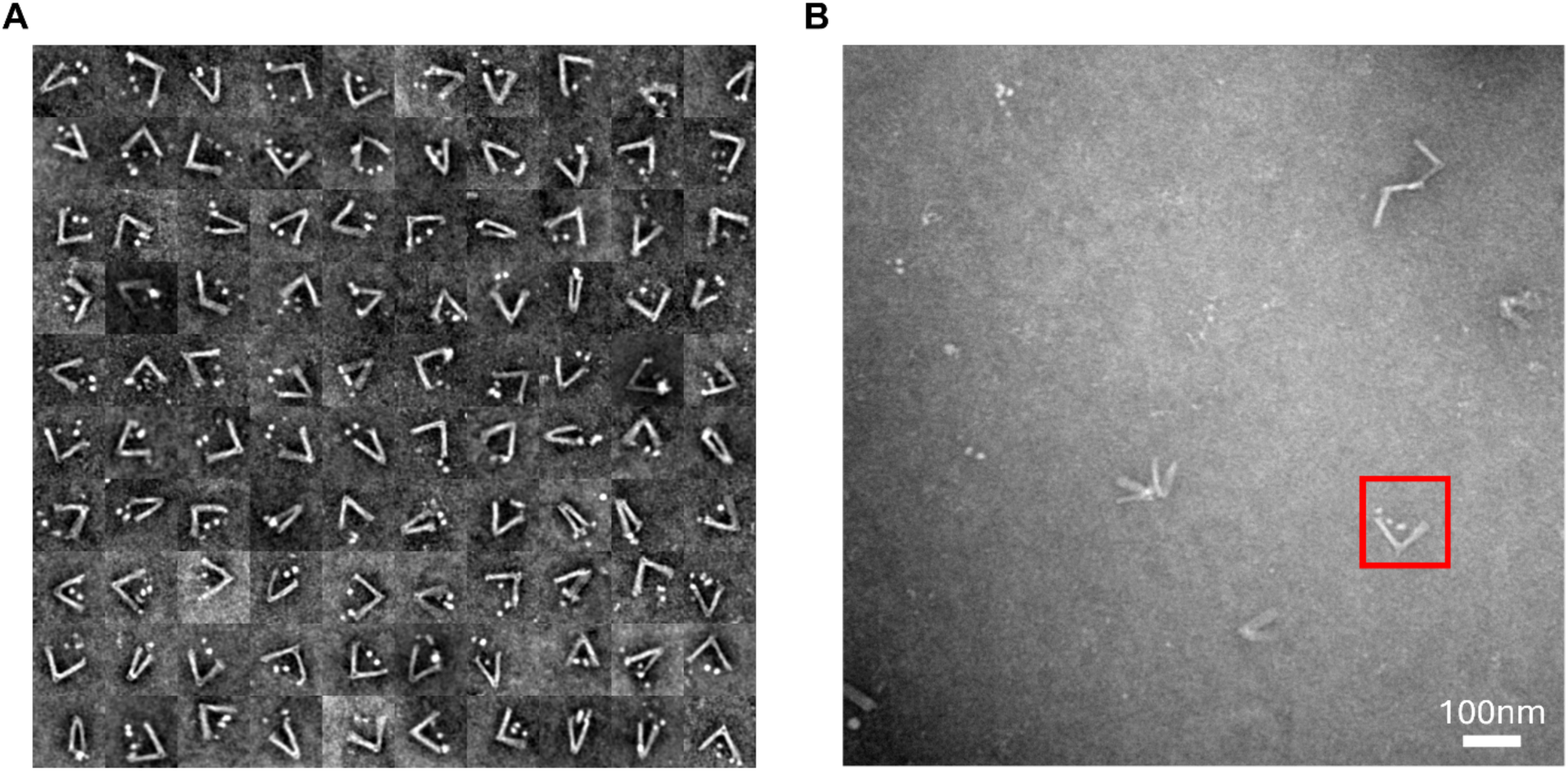
(**A**) Image gallery of 100 nDFS.B_TNuc_ nanocalipers in 0.25 mM Mg^2+^ that contain properly integrated tetranucleosomes. They were selected at random from the full dataset of 266 particles. (**B**) Example TEM image with example nDFS.B_TNuc_ in the red box.

**Figure S16:**
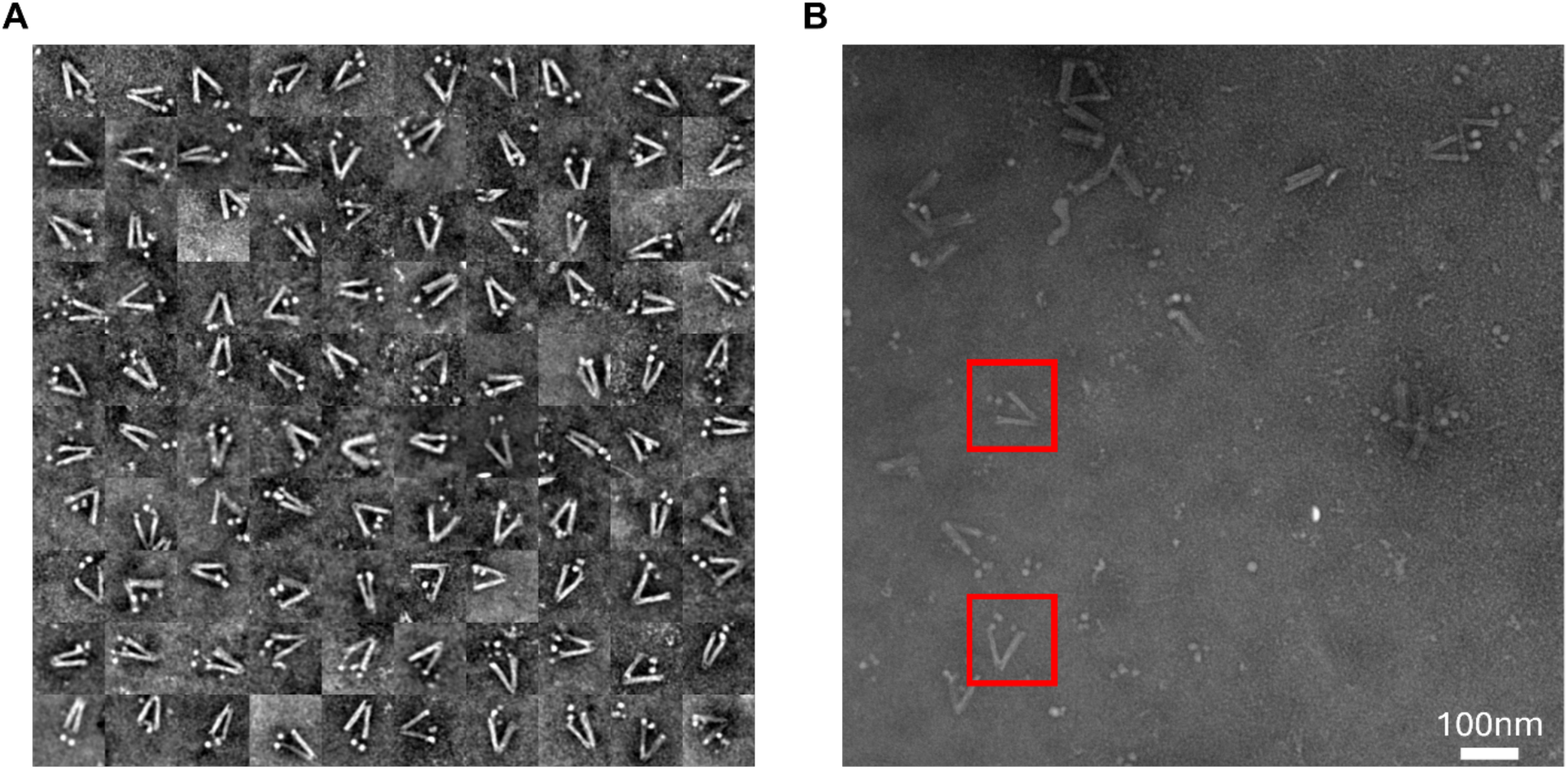
(**A**) Image gallery of 100 nDFS.C_TNuc_ nanocalipers in 0.25 mM Mg^2+^ that contain properly integrated tetranucleosomes. They were selected at random from the full dataset of 388 particles. (**B**) Example TEM image with example nDFS.C_TNuc_ in red boxes.

**Figure S17:**
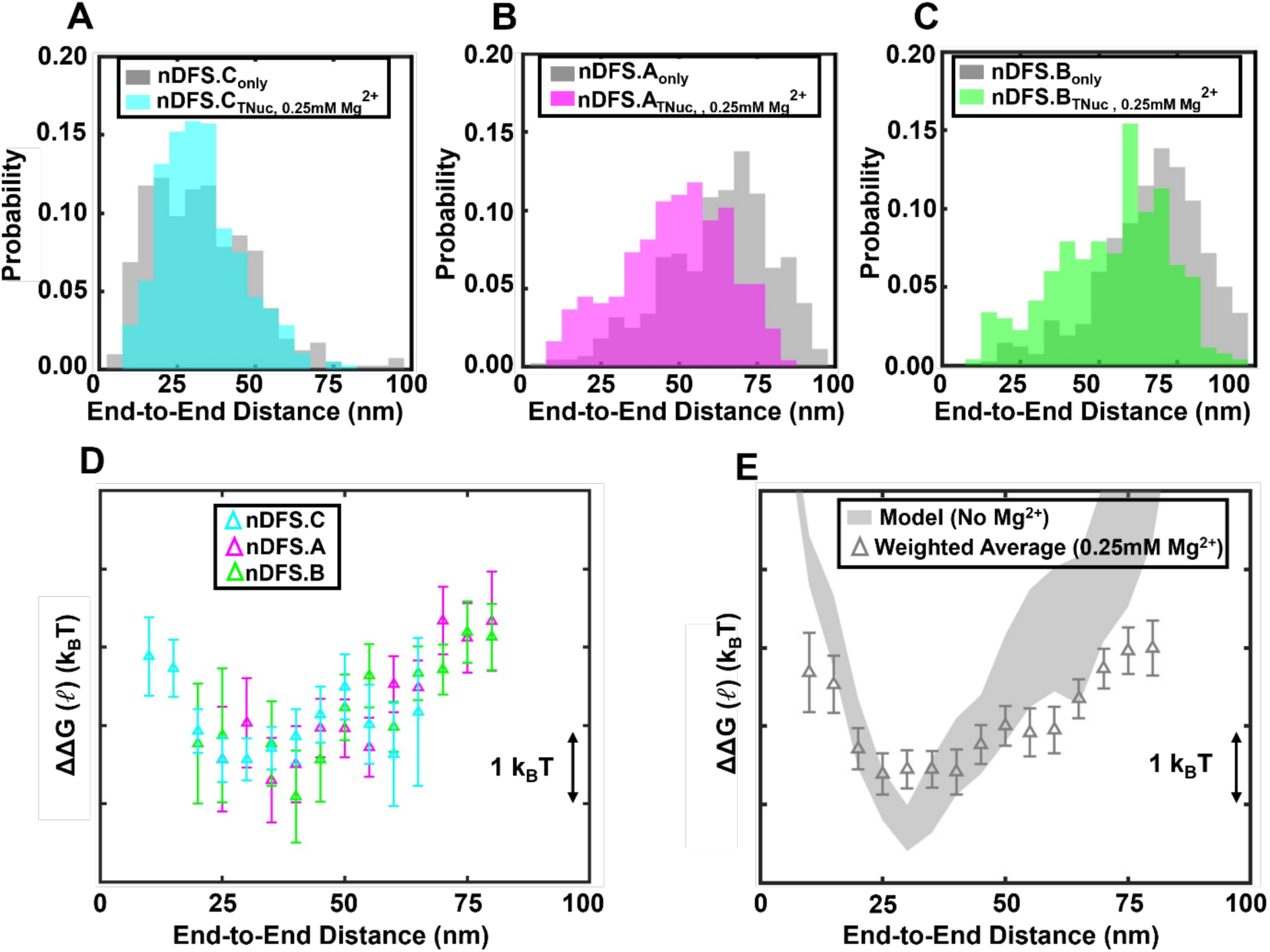
End-to-end free energy at low ionic conditions. (**A**-**C**) End-to-end distance probability distributions for nanocalipers are shown in grey for each nanocaliper design (nDFS.C_only_, nDFS.A_only_, nDFS.B_only_). End-to-end distance probability distributions of nanocalipers with tetranucleosomes (nDFS.C_TNuc_, nDFS.A_TNuc_, nDFS.B_TNuc_) in 0.25 mM Mg^2+^ are shown in (A) cyan, (B) magenta, and (C) green, respectively. (**D**) End-to-end distance tetranucleosome free energy landscapes from each nanocaliper design. The error bars of the free energy values were propagated from the uncertainties of the probability measurements, which were estimated by the square root of the number of events in each bin. (**E**) End-to-end distance tetranucleosome free energy landscapes determined from the weighted average of the three independent free energy measurements using the three different nanocalipers in 0.25 mM Mg^2+^ (open grey triangles). The error bars of the free energies were determined as described in the Methods section. The free energy landscape derived from molecular dynamics simulations of the coarse-grained model is plotted as the shaded grey area, which represents the 95% confidence interval.

**Figure S18:**
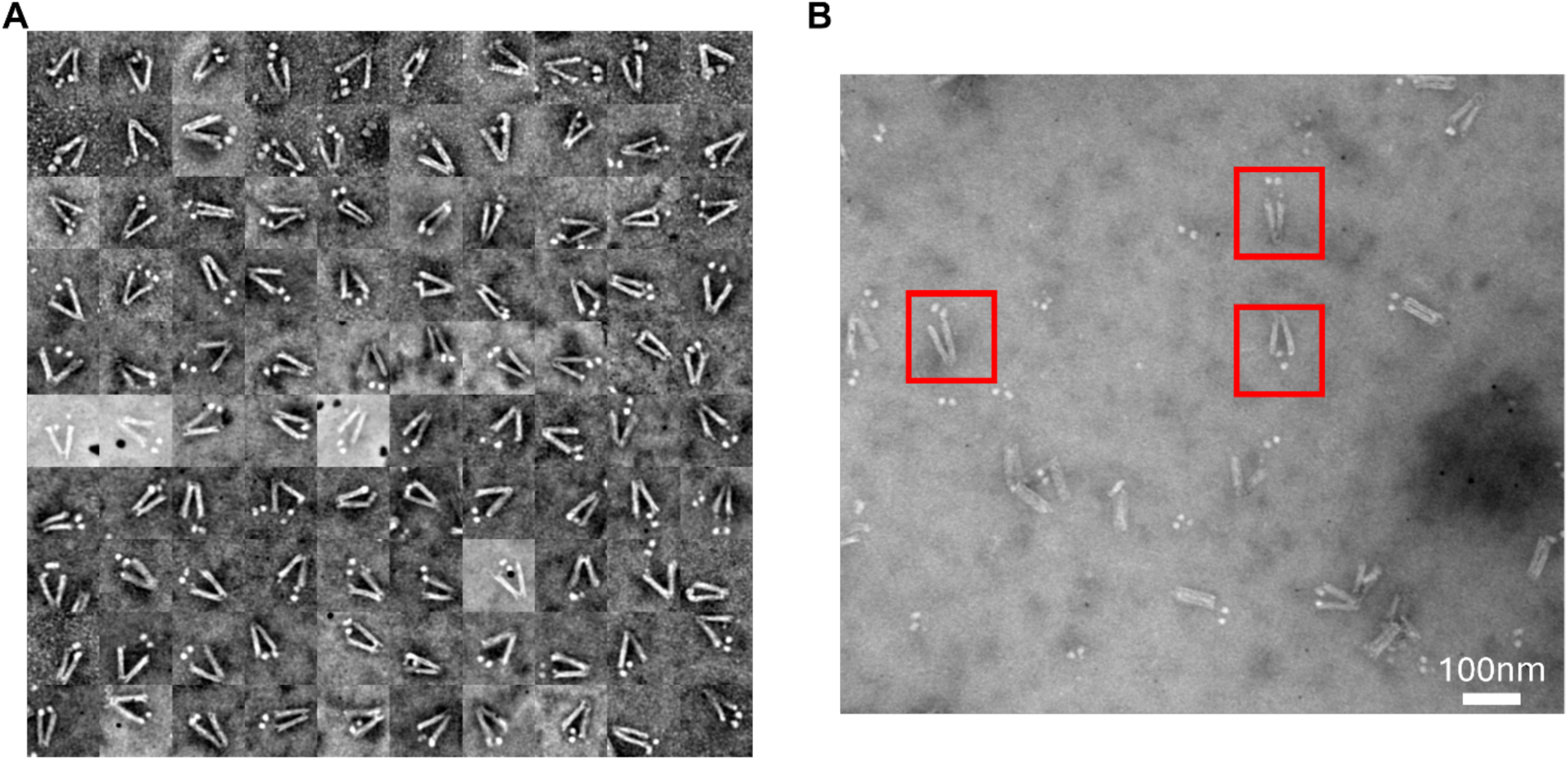
(**A**) Image gallery of 100 nDFS.C_TNuc,H1=5nM_ nanocalipers with 5 nM H1.0 that contain properly integrated tetranucleosomes. They were selected at random from the full dataset of 332 particles. (**B**) Example TEM image with example nDFS.C_TNuc,H1=5nM_ in red boxes.

**Figure S19:**
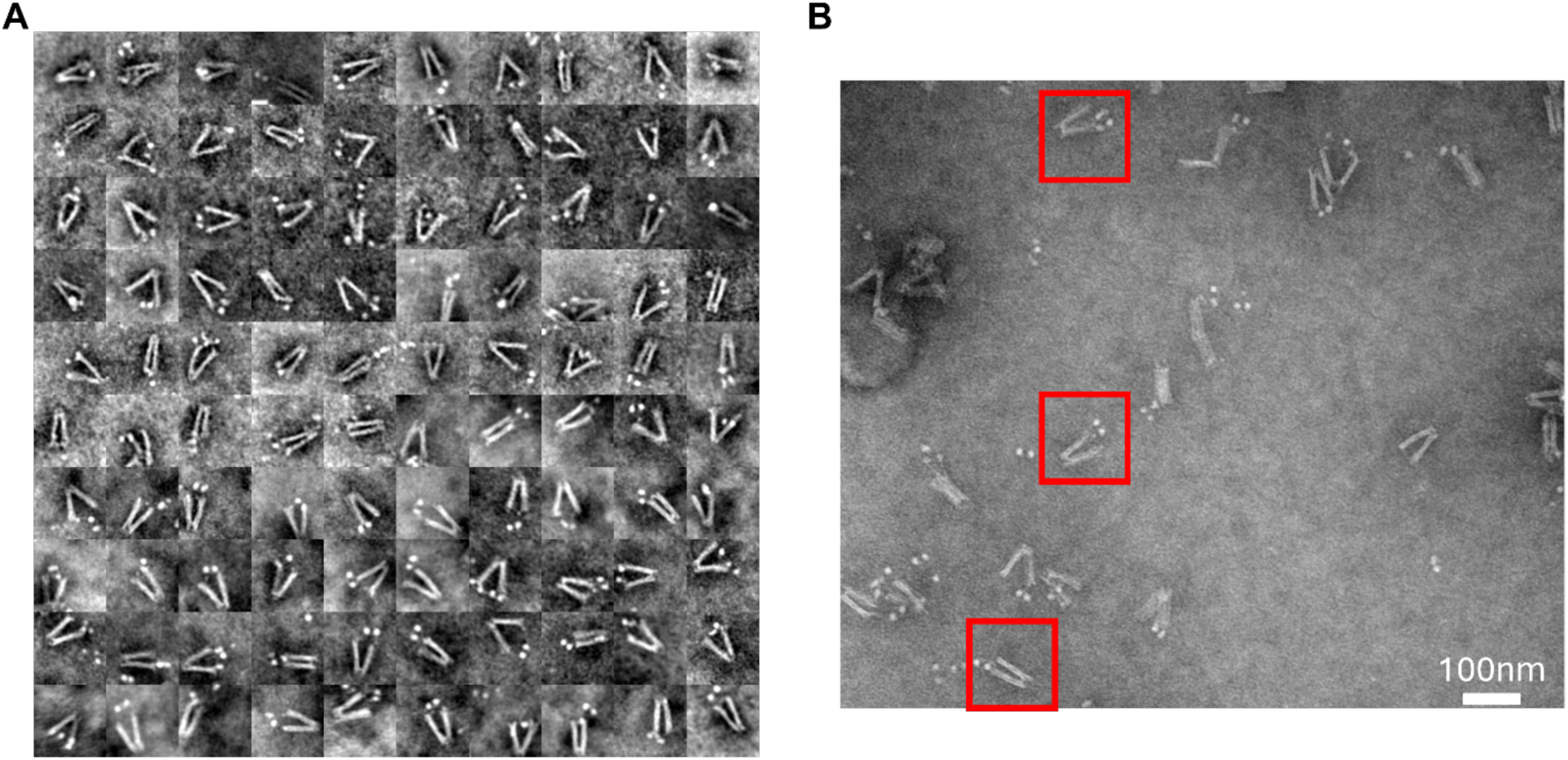
(**A**) Image gallery of 100 nDFS.C_TNuc,H1=10nM_ nanocalipers with 10 nM H1.0 that contain properly integrated tetranucleosomes. They were selected at random from the full dataset of 590 particles. (**B**) Example TEM image with example nDFS.C_TNuc,H1=10nM_ in red boxes.

**Figure S20:**
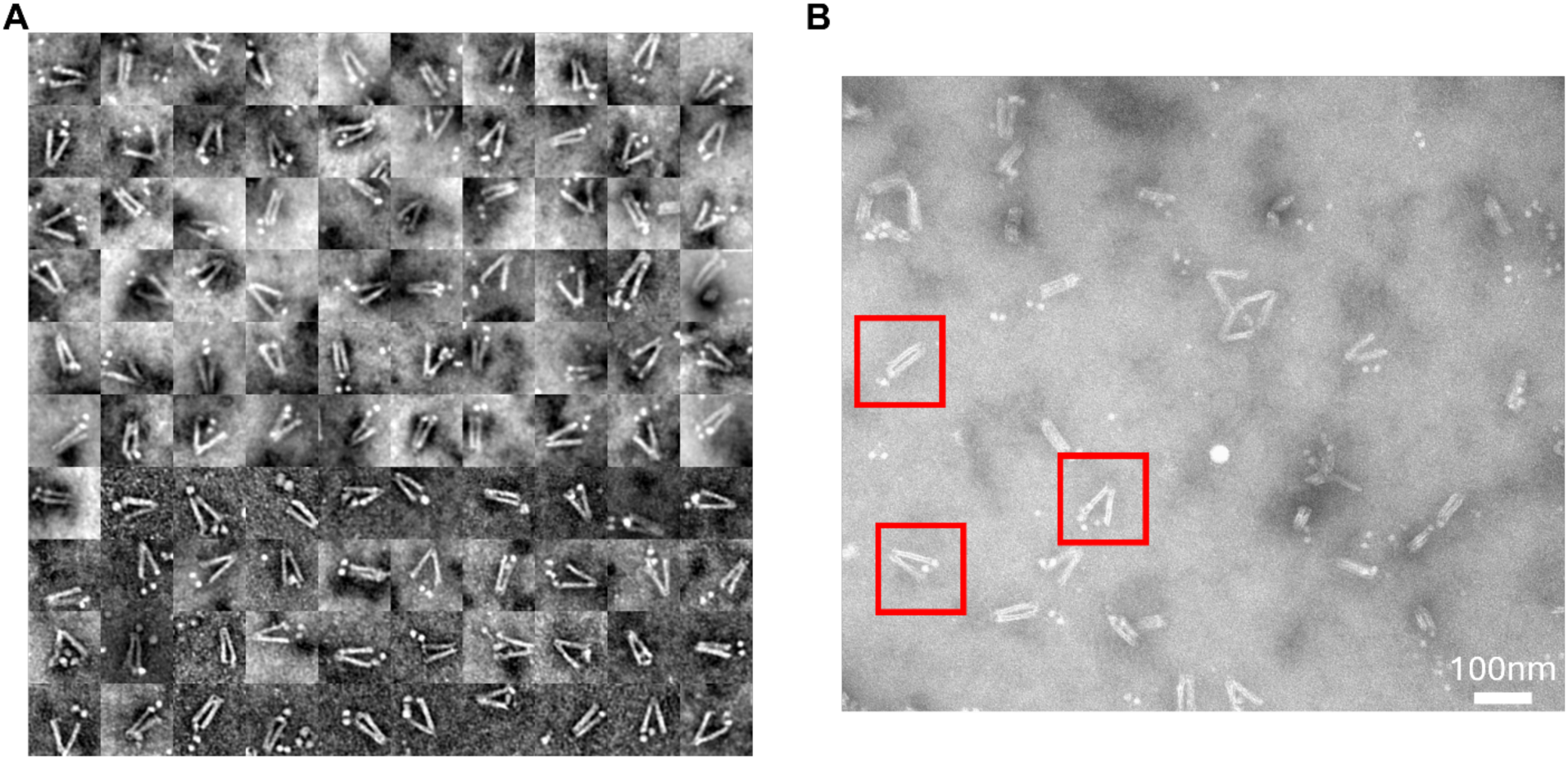
(**A**) Image gallery of 100 nDFS.C_TNuc,H1=20nM_ nanocalipers with 20 nM H1.0 that contain properly integrated tetranucleosomes. They were selected at random from the full dataset of 637 particles. (**B**) Example TEM image with example nDFS.C_TNuc,H1=20nM_ in red boxes.

**Figure S21:**
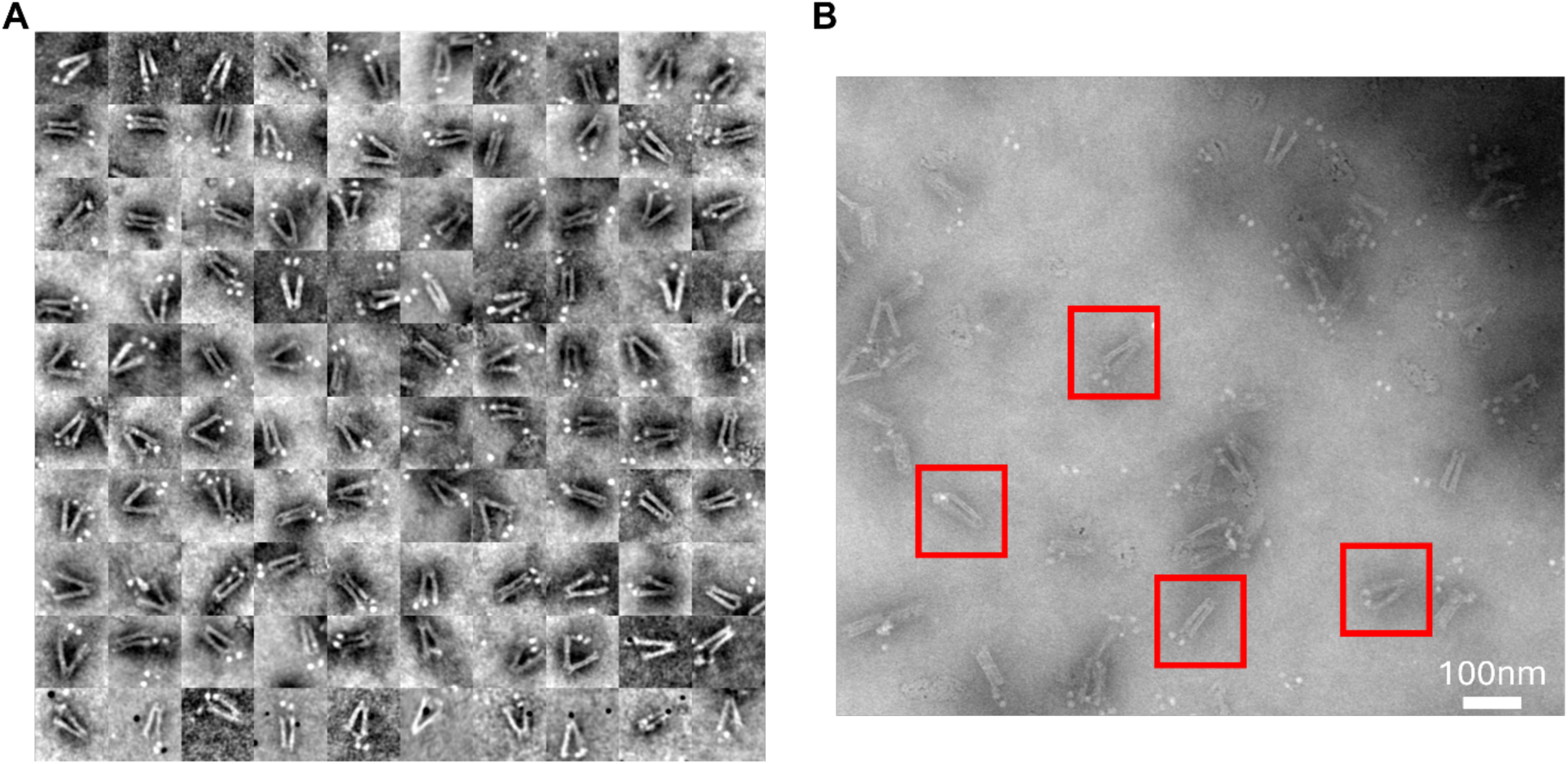
(**A**) Image gallery of 100 nDFS.C_TNuc,H1=40nM_ nanocalipers with 40 nM H1.0 that contain properly integrated tetranucleosomes. They were selected at random from the full dataset of 711 particles. (**B**) Example TEM image with example nDFS.C_TNuc,H1=40nM_ in red boxes.

**Figure S22:**
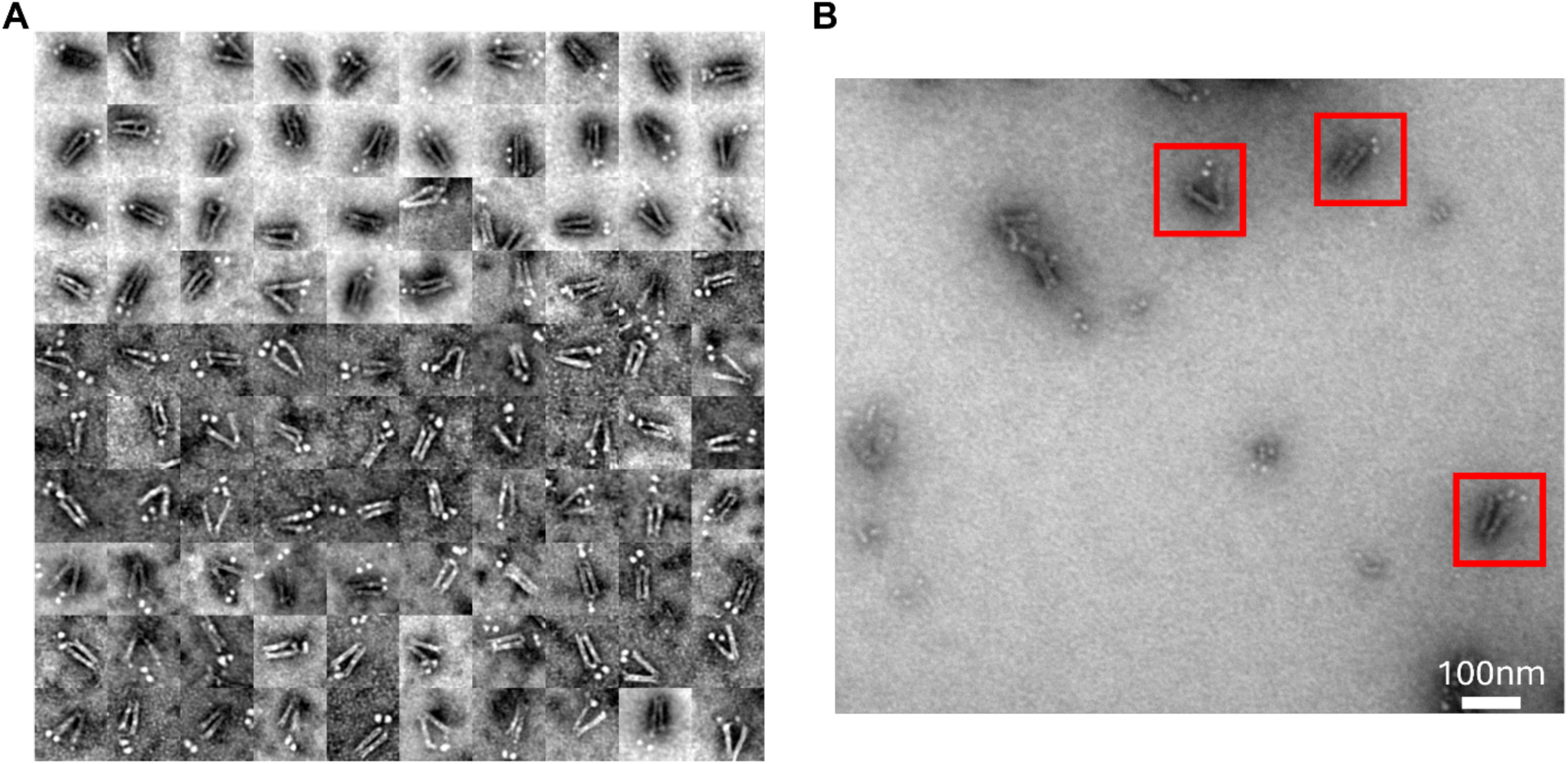
(**A**) Image gallery of 100 nDFS.C_TNuc,H1=80nM_ nanocalipers with 80 nM H1.0 that contain properly integrated tetranucleosomes. They were selected at random from the full dataset of 521 particles. (**B**) Example TEM image with example nDFS.C_TNuc,H1=80nM_ in red boxes.

**Figure S23:**
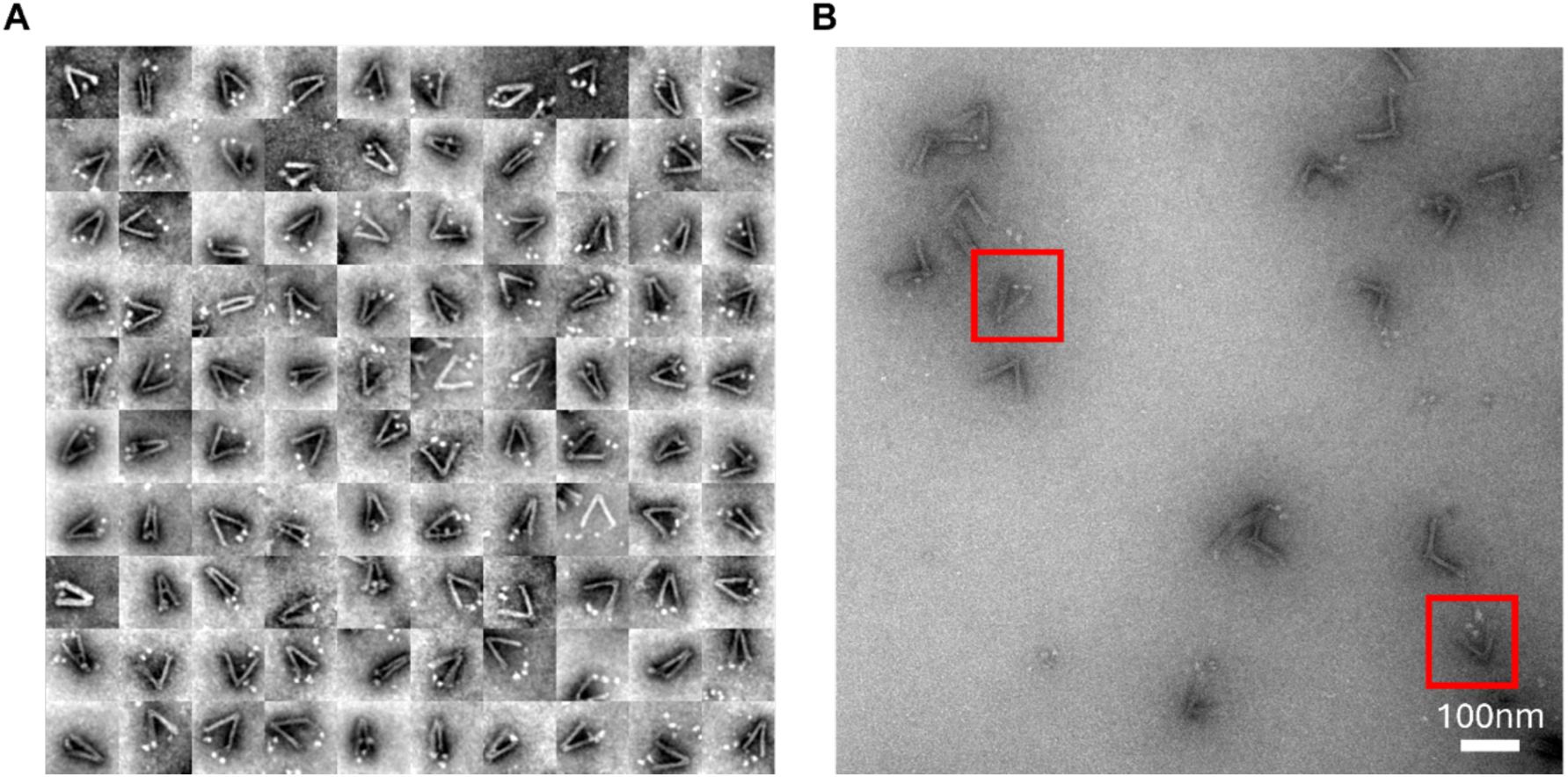
(**A**) Image gallery of 100 nDFS.A_TNuc,H1=10nM_ nanocalipers with 10 nM H1.0 that contain properly integrated tetranucleosomes. They were selected at random from the full dataset of 279 particles. (**B**) Example TEM image with example nDFS.A_TNuc,H1=10nM_ in red boxes.

**Figure S24:**
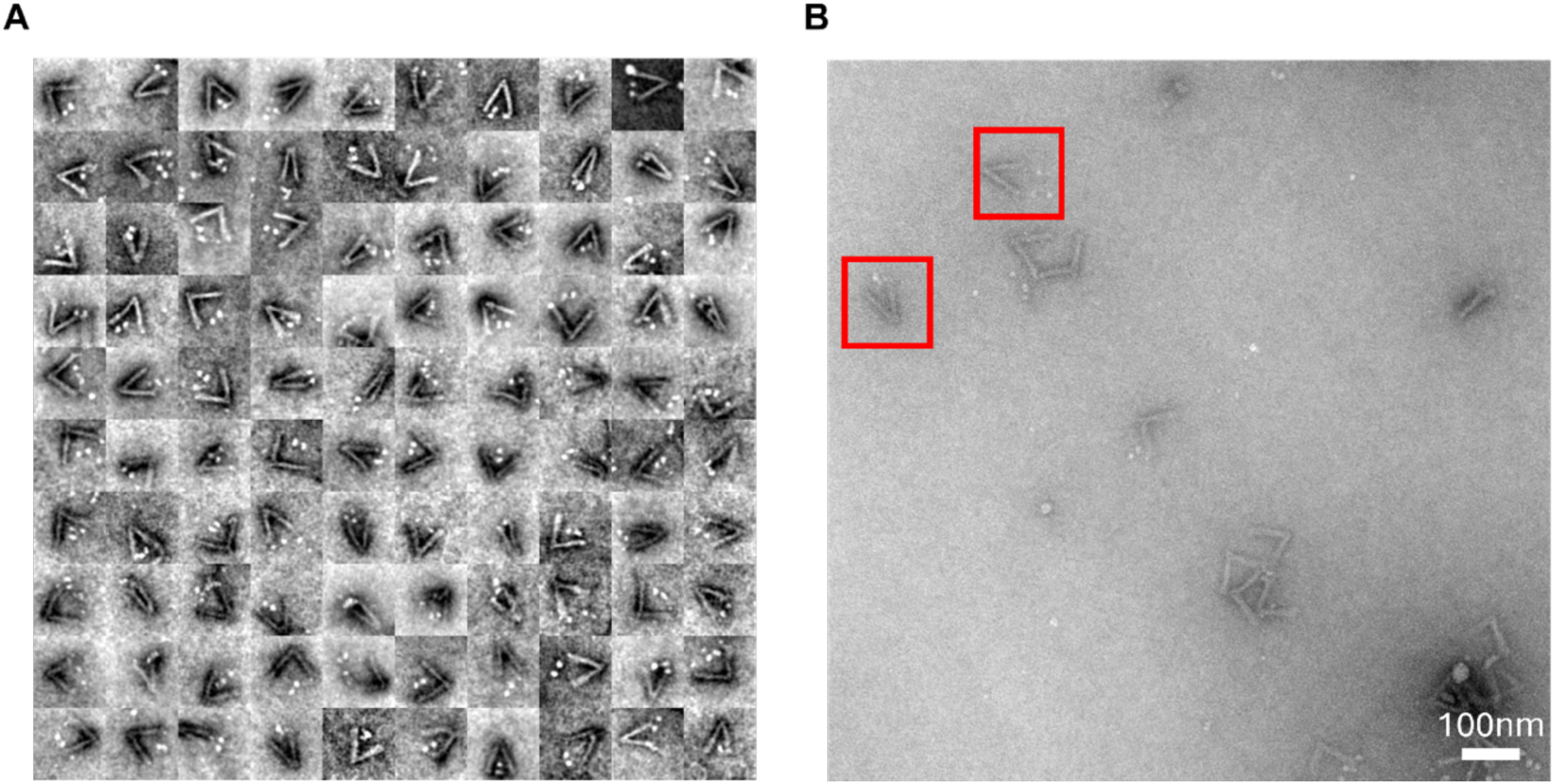
(**A**) Image gallery of 100 nDFS.B_TNuc,H1=10nM_ nanocalipers with 10 nM H1.0 that contain properly integrated tetranucleosomes. They were selected at random from the full dataset of 309 particles. (**B**) Example TEM image with example nDFS.B_TNuc,H1=10nM_ in red boxes.

**Figure S25:**
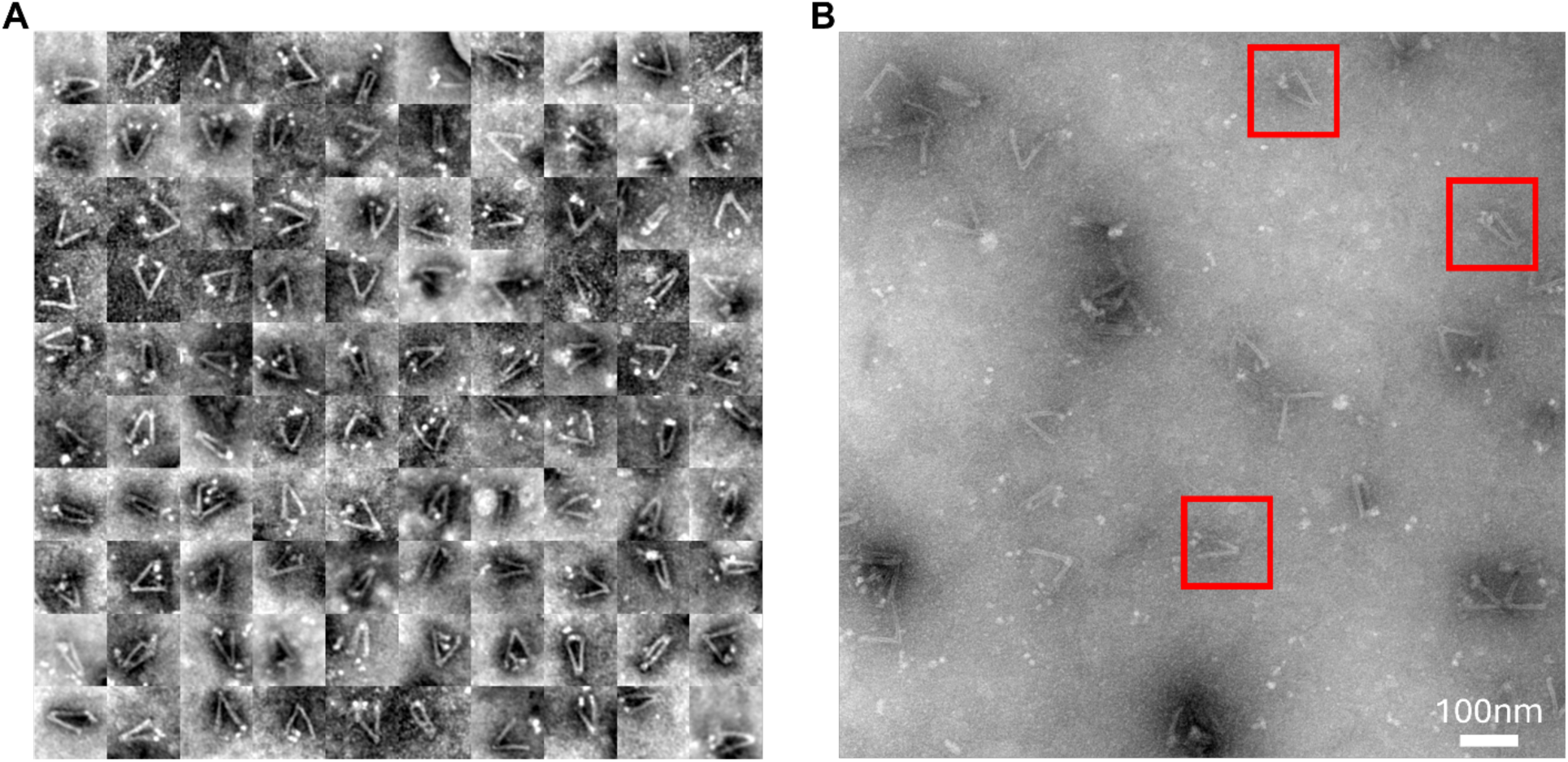
(**A**) Image gallery of 100 nDFS.A_TNuc,H1=40nM_ nanocalipers with 40 nM H1.0 that contain properly integrated tetranucleosomes. They were selected at random from the full dataset of 214 particles. (**B**) Example TEM image with example nDFS.A_TNuc,H1=40nM_ in red boxes.

**Figure S26:**
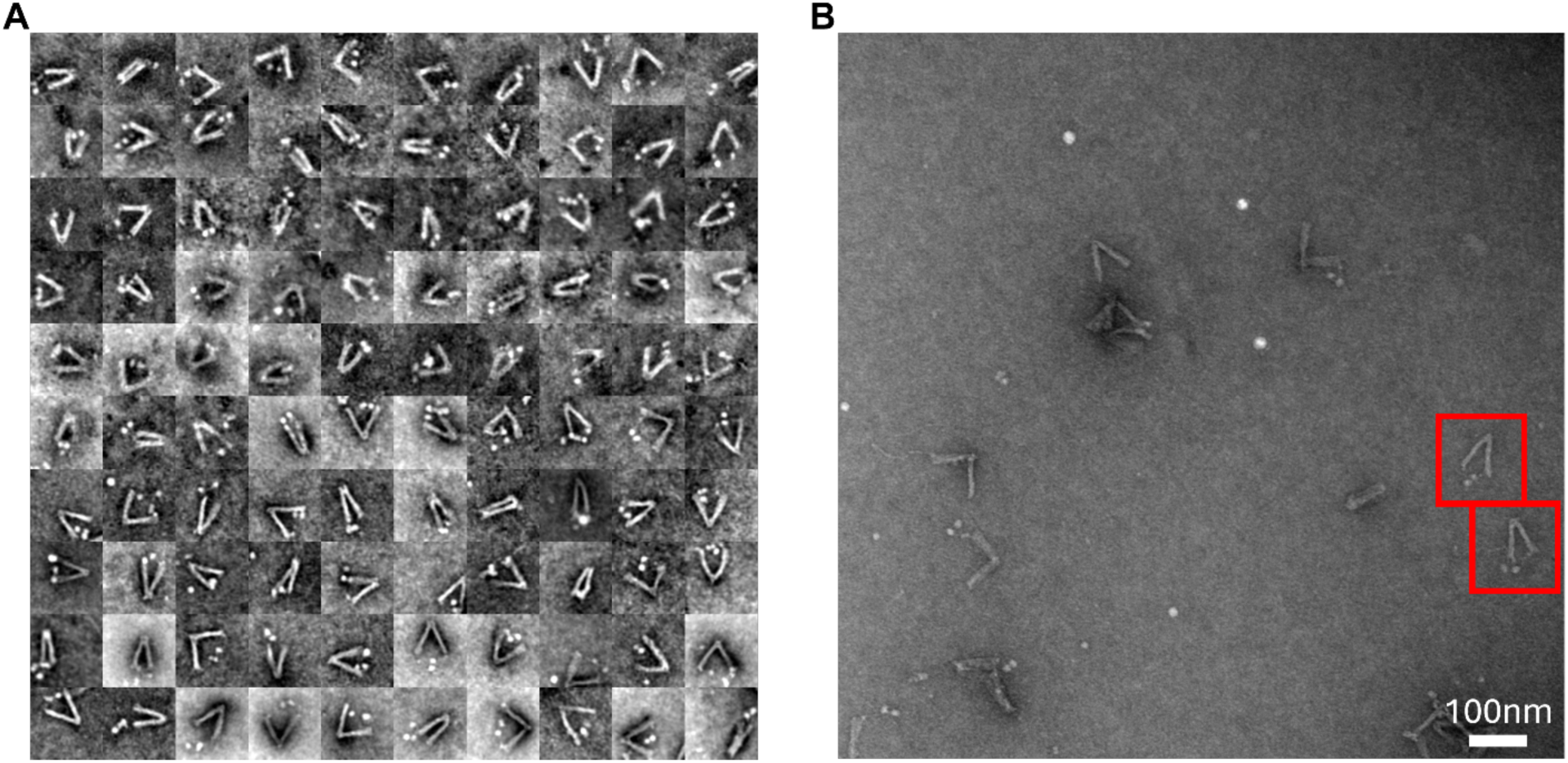
(**A**) Image gallery of 100 nDFS.B_TNuc,H1=40nM_ nanocalipers with 40 nM H1.0 that contain properly integrated tetranucleosomes. They were selected at random from the full dataset of 630 particles. (**B**) Example TEM image with example nDFS.B_TNuc,H1=40nM_ in red boxes.

**Figure S27:**
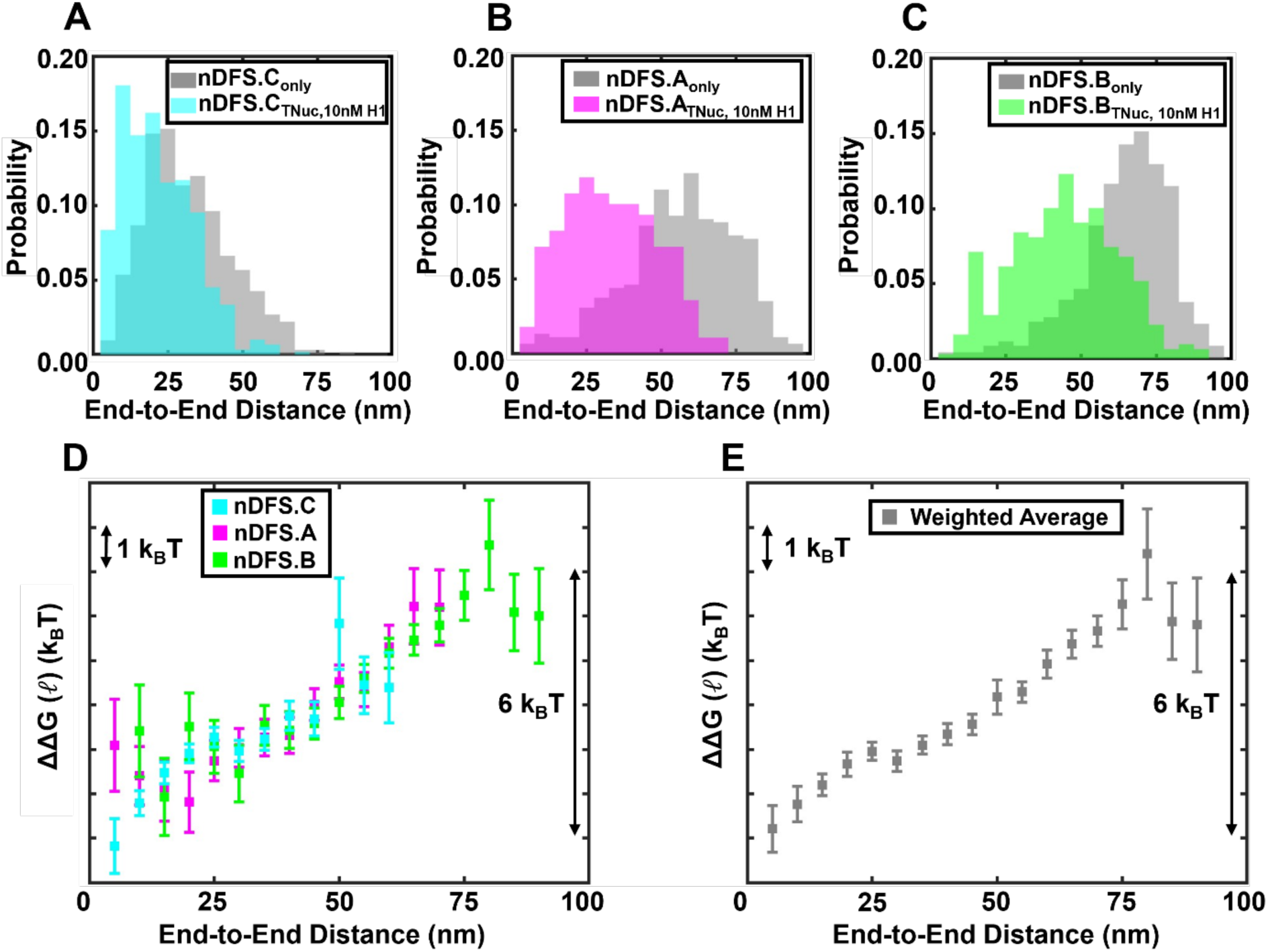
End-to-end free energy at 10nM H1.0. (**A**-**C**) End-to-end distance probability distributions for nanocalipers are shown in grey for each nanocaliper design (nDFS.C_only_, nDFS.A_only_, nDFS.B_only_). End-to-end distance probability distributions of nanocalipers with tetranucleosomes (nDFS.C_TNuc_, nDFS.A_TNuc_, nDFS.B_TNuc_) with 10nM H1.0 are shown in (A) cyan, (B) magenta, and (C) green, respectively. (**D**) End-to-end distance tetranucleosome free energy landscapes from each nanocaliper design. The error bars of the free energy values were propagated from the uncertainties of the probability measurements, which were estimated by the square root of the number of events in each bin. (**E**) End-to-end distance tetranucleosome free energy landscapes determined from the weighted average of the three independent free energy measurements using the three different nanocalipers with 10 nM (closed grey squares). The error bars of the free energies were determined as described in the Methods section.

**Figure S28:**
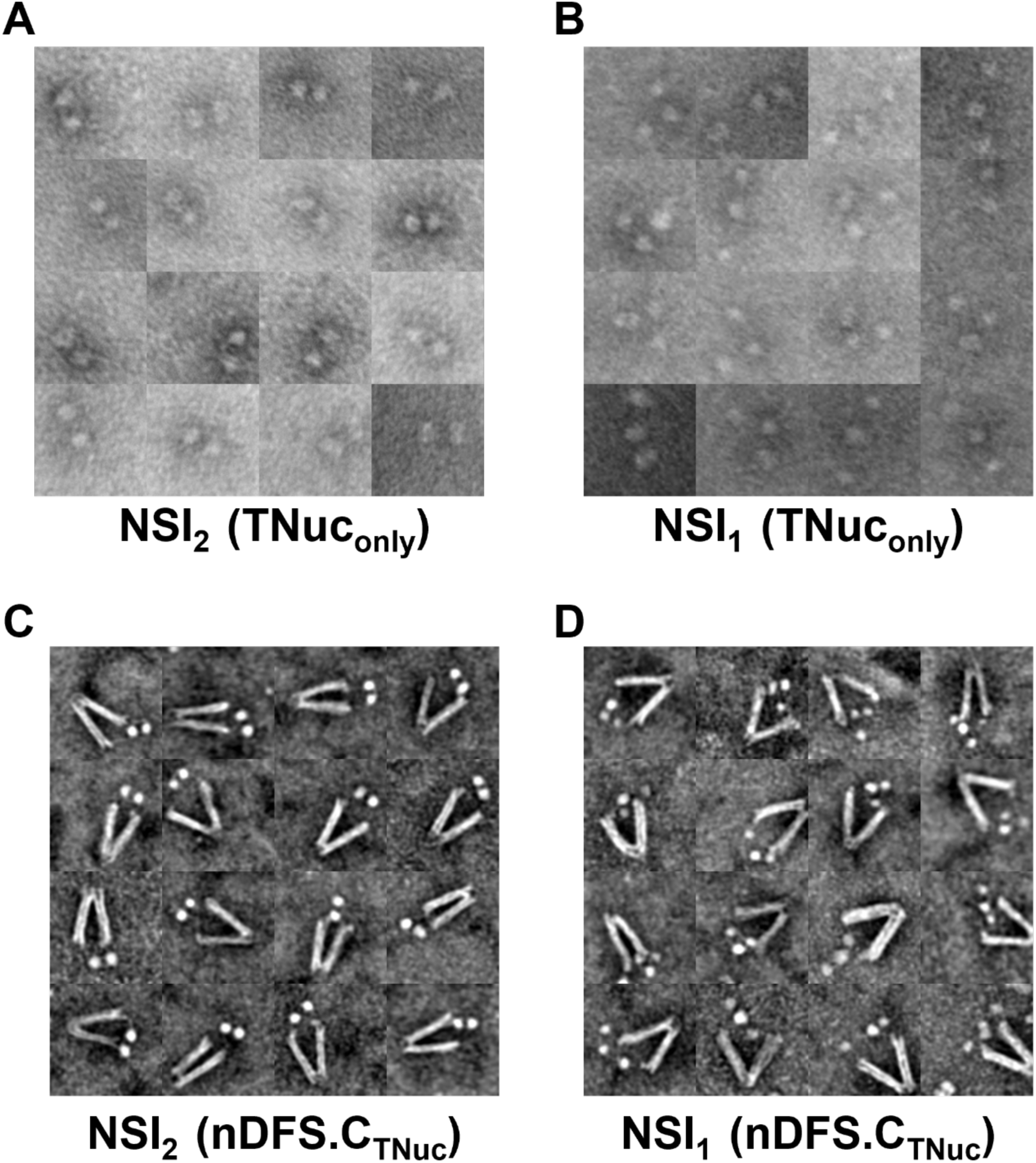
Example images of fully (NSI_2_) and partially (NSI_1_) compacted tetranucleosomes. (**A**-**B**) Image gallery of tetranucleosomes (TNuc_only_) in 200 mM NaCl and 1 mM MgCl_2_, that are in the NSI_2_ (A) and NSI_1_ (B) states. (**C**-**D**) Image gallery of tetranucleosomes incorporated into nDFS.C in 200 mM NaCl and 1 mM MgCl_2_, which are in the NSI_2_ (C) and NSI_1_ (D) states.

**Figure S29:**
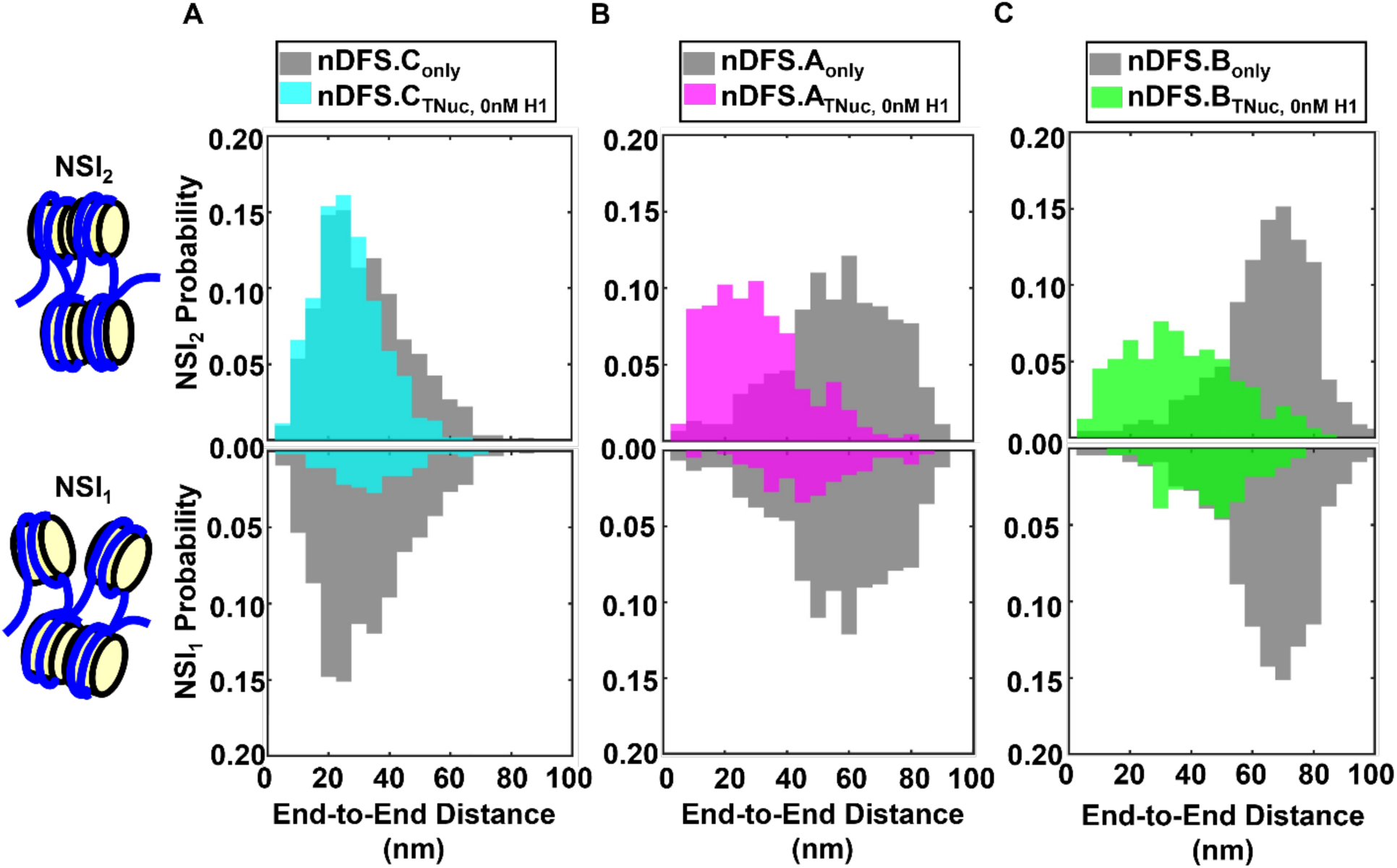
End-to-end distance distributions for the NSI_2_ and NSI_1_ tetranucleosomes states without H1. (**A**-**C**) End-to-end distance distributions for NSI_2_ (top upright) and NSI_1_ (bottom inverted) tetranucleosome states in nDFS.C (cyan), nDFS.A (magenta), and nDFS.B (green), respectively. The end-to-end distance distributions of nDFS.C_only_, nDFS.A_only_, and nDFS.B_only_ are plotted in grey in each plot for reference. The probability distributions for NSI_1_ vs. NSI_2_ states are from the same dataset, so the relative area of the top (NSI_1_) and bottom (NSI_2_) are the relative probabilities of the NSI_2_ and NSI_1_ states. Also, the sum of the NSI_2_ and NSI_1_ probabilities is 1.

**Figure S30:**
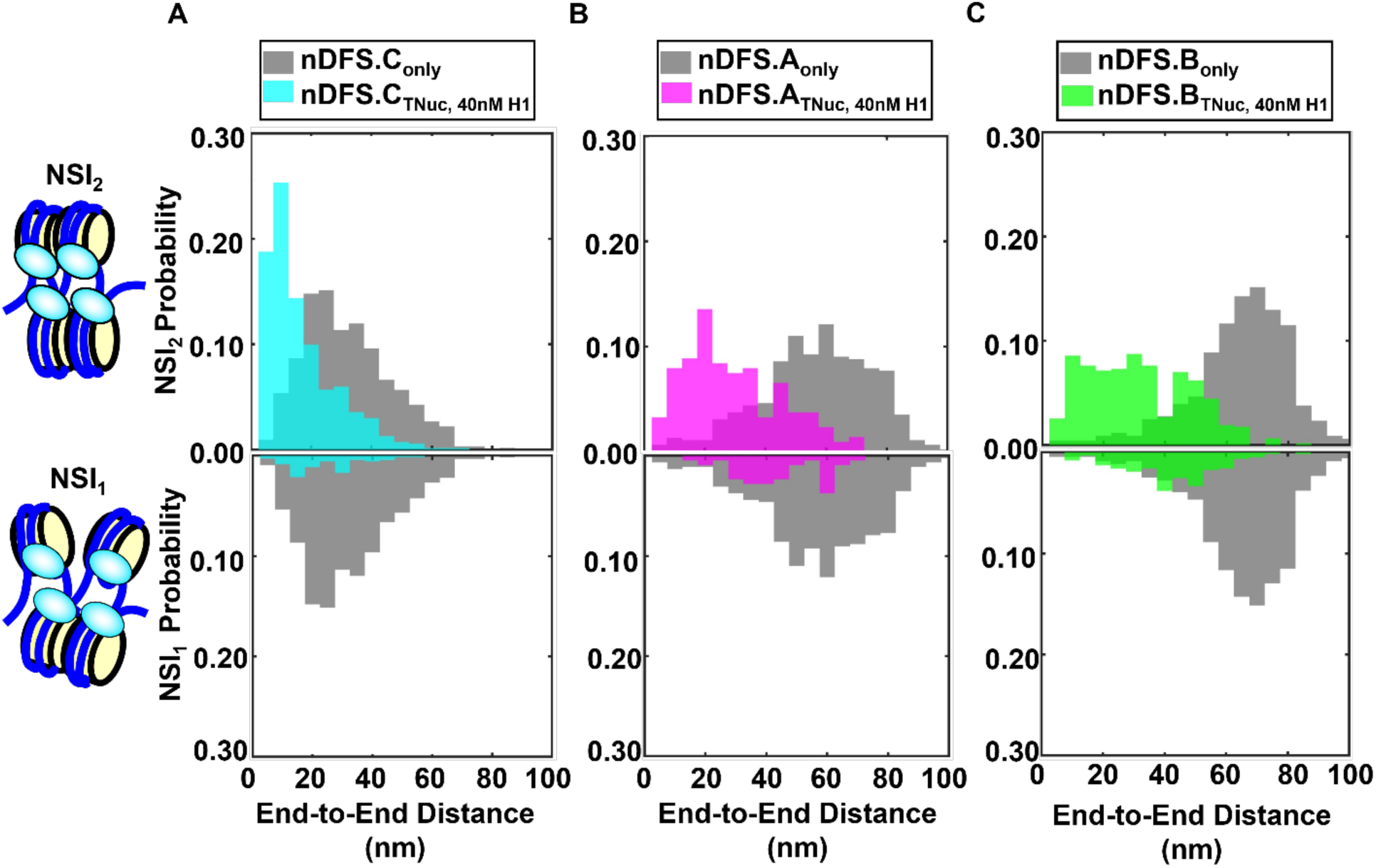
End-to-end distance distributions for the NSI_2_ and NSI_1_ tetranucleosomes states with 40 nM H1. (**A**-**C**) End-to-end distance distributions for NSI_2_ (top upright) and NSI_1_ (bottom inverted) tetranucleosome states in nDFS.C (cyan), nDFS.A (magenta), and nDFS.B (green), respectively, with 40 nM H1.0. The end-to-end distance distributions of nDFS.C_only_, nDFS.A_only_, and nDFS.B_only_ are plotted in grey in each plot for reference. The probability distributions for NSI_1_ vs. NSI_2_ states are from the same dataset, so the relative area of the top (NSI_1_) and bottom (NSI_2_) are the relative probabilities of the NSI_2_ and NSI_1_ states. Also, the sum of the NSI_2_ and NSI_1_ probabilities is 1.

**Figure S31:**
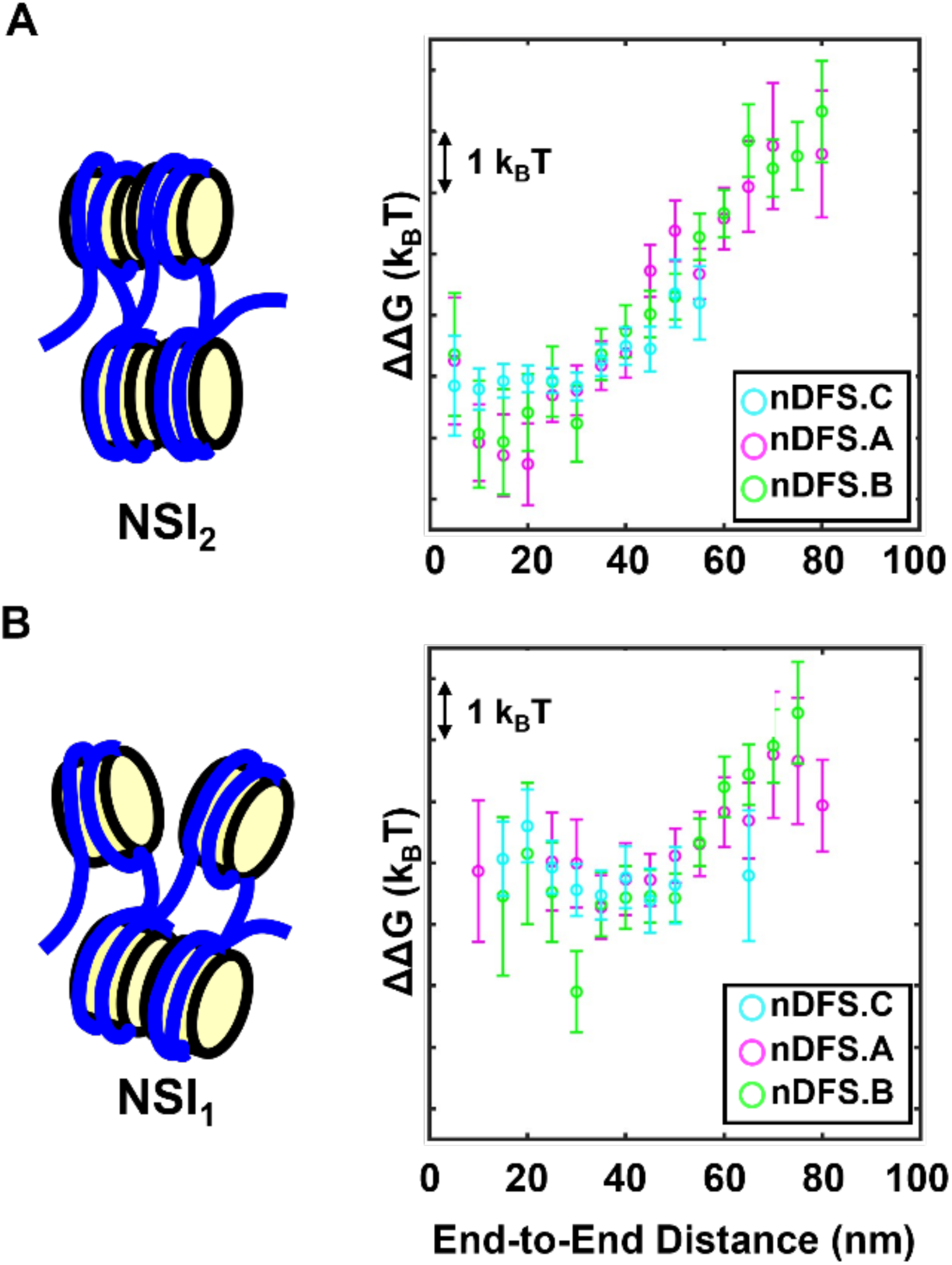
End-to-end distance free energy landscapes for the NSI_2_ and NSI_1_ tetranucleosome states in the absence of H1.0 measured using individual nanocalipers. (**A**) End-to-end distance free energy landscape of the NSI_2_ tetranucleosome state determined with nDFS.C (cyan), nDFS.A (magenta), and nDFS.B (green). The error bars of the free energy values were propagated from the uncertainty of the probability measurements, which were estimated by the square root of the number of events in each bin. (**B**) End-to-end distance free energy landscape of the NSI_1_ tetranucleosome state determined with nDFS.C (cyan), nDFS.A (magenta), and nDFS.B (green). The error bars of the free energy values were propagated from the uncertainty of the probability measurements, which were estimated by the square root of the number of events in each bin.

**Figure S32:**
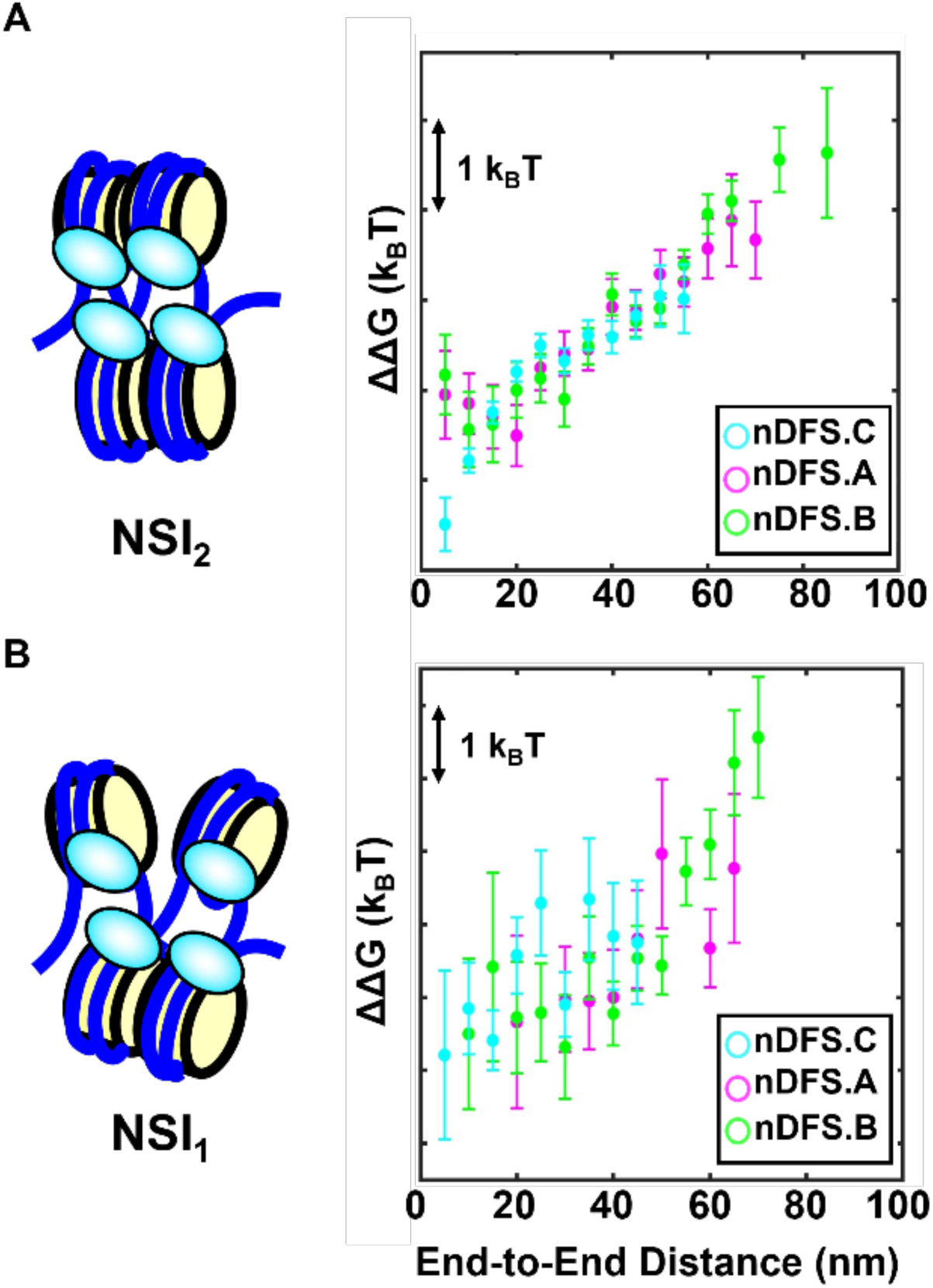
End-to-end distance free energy landscapes for NSI_2_ and NSI_1_ tetranucleosome states in the presence of 40nM H1 measured using individual nanocalipers. (**A**) End-to-end distance free energy landscape of the NSI_2_ tetranucleosome state with 40 nM H1.0 determined with nDFS.C (cyan), nDFS.A (magenta), and nDFS.B (green). The error bars of the free energy values were propagated from the uncertainty of the probability measurements, which were estimated by the square root of the number of events in each bin. (**B**) End-to-end distance free energy landscape of the NSI_1_ tetranucleosome state with 40 nM H1.0 determined with nDFS.C (cyan), nDFS.A (magenta), and nDFS.B (green). The error bars of the free energy values were propagated from the uncertainty of the probability measurements, which were estimated by the square root of the number of events in each bin.

**Figure S33:**
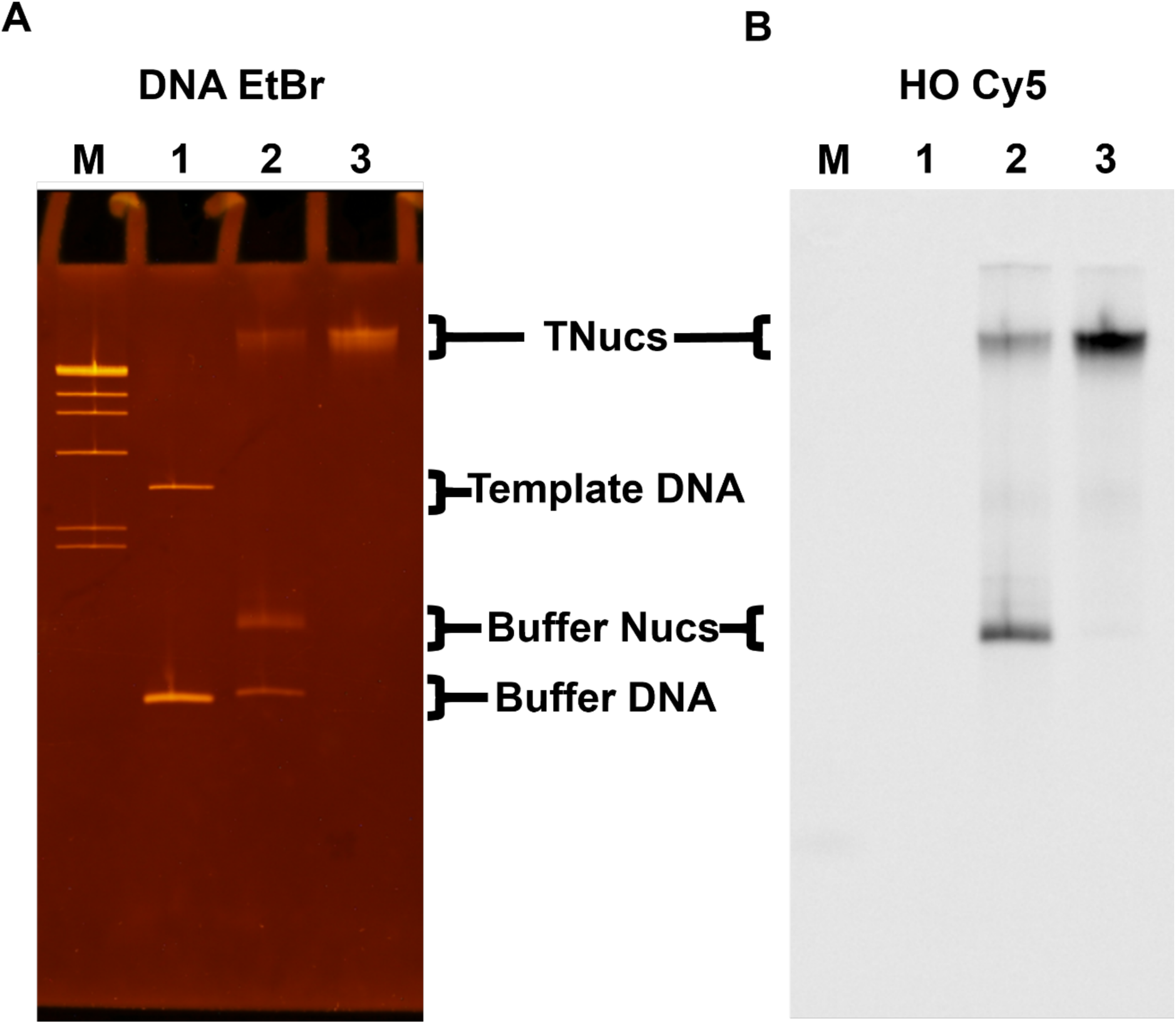
Native polyacrylamide gel electrophoresis analysis of tetranuclesomes. Native 4% polyacrylamide native gel with 1kB DNA ladder (M), DNA only (1), pre-purified tetranuclesomes (2), and sucrose gradient purified tetranucleosomes (3). (**A**) Ethidium bromide (EtBr) fluorescence image of the gel post-stained with EtBr. (**B**) Cy5 fluorescence image of the same gel in (A). H2AK119C is Cy5-labeled, so only nucleosomes are detected in the image.

**Figure S34:**
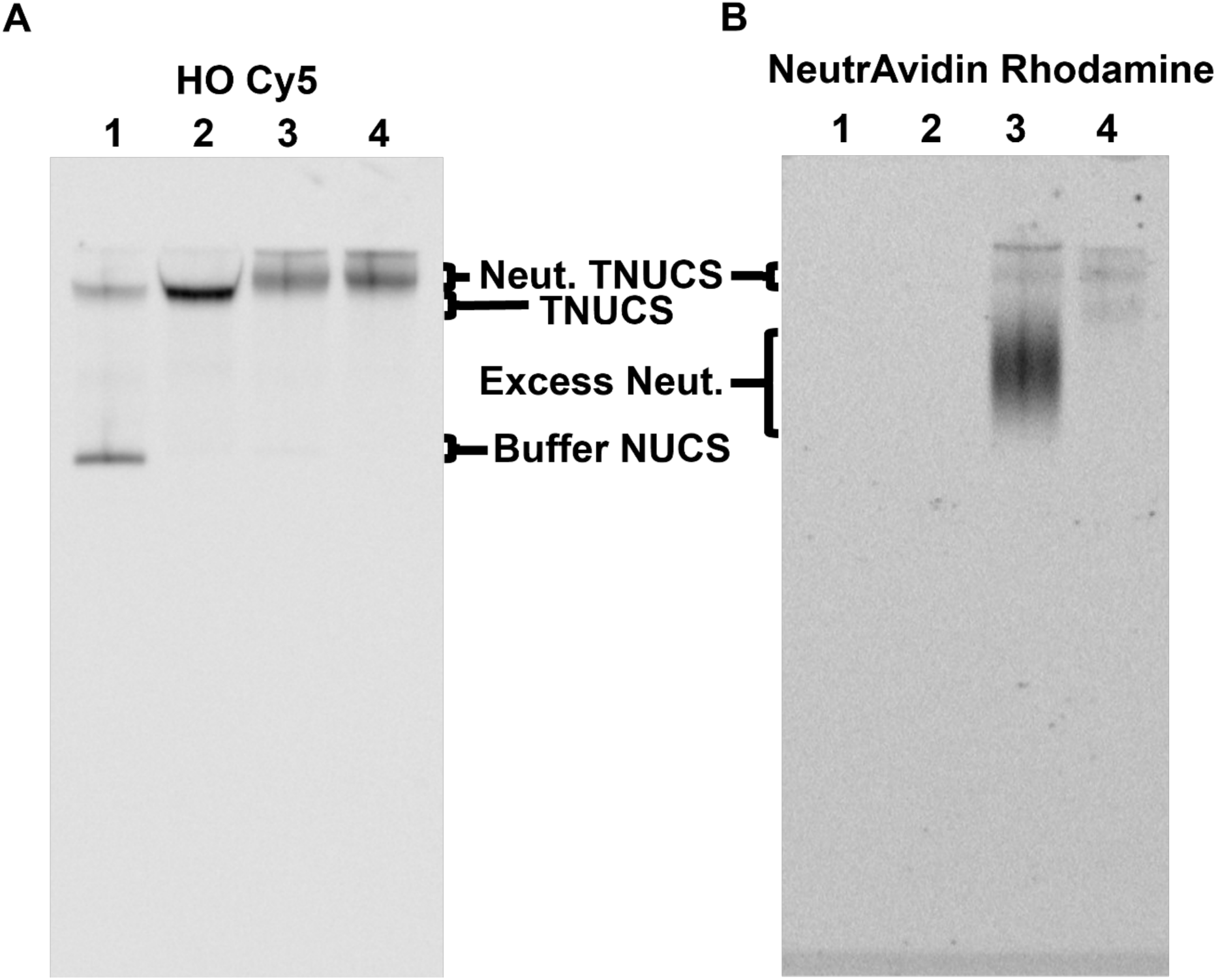
Native polyacrylamide gel electrophoresis analysis of tetranucleosomes labeled with NeutrAvidin. Native 4% polyacrylamide gel with pre-purified tetranucleosomes (1), and sucrose gradient purified tetranucleosomes (2), NeutrAvidin labeled tetranucleosomes before the second sucrose gradient (3), and NeutrAvidin labeled tetranucleosomes following the second sucrose gradient, which removed excess unbound NeutrAvidin (4). (**A**) Cy5 fluorescence image, which visualizes Cy5-labeled H2AK119C. (**B**) Rhodamine fluorescence image, which visualizes Rhodamine-labeled NeutrAvidin.

### Supplementary Tables

**Table S1:** Number of TEM images and total devices analyzed for nanocalipers alone (nDFS_only_) and nanocalipers with tetranucleosomes (nDFS_TNuc_) in 1 or 0.25 mM Mg^2+^.

|  | [Mg<sup>2+</sup>] | nDFS.C |  | nDFS.A |  | nDFS.B |  |
| --- | --- | --- | --- | --- | --- | --- | --- |
|  |  | Images | Devices | Images | Devices | Images | Devices |
| nDFS<sub>only</sub> | 1 mM | 628 | 635 | 150 | 545 | 227 | 687 |
| nDFS<sub>only</sub> | 0.25 mM | 220 | 408 | 215 | 443 | 284 | 419 |
| nDFS<sub>TNuc</sub> | 1 mM | 346 | 546 | 280 | 439 | 421 | 486 |
| nDFS<sub>TNuc</sub> | 0.25 mM | 209 | 209 | 215 | 246 | 284 | 266 |

**Table S2:** Number of TEM images and total devices analyzed for nanocalipers with tetranucleosomes at different concentrations of H1.0.

| [H1.0] | nDFS.C<sub>TNuc</sub> |  | nDFS.A<sub>TNuc</sub> |  | nDFS.B<sub>TNuc</sub> |  |
| --- | --- | --- | --- | --- | --- | --- |
|  | Images | Devices | Images | Devices | Images | Devices |
| 5 nM | 263 | 332 | NA | NA | NA | NA |
| 10 nM | 290 | 590 | 140 | 279 | 272 | 309 |
| 20 nM | 400 | 637 | NA | NA | NA | NA |
| 40 nM | 405 | 711 | 107 | 214 | 586 | 630 |
| 80 nM | 359 | 521 | NA | NA | NA | NA |

**Table S3:** Nucleosome stacking interaction frequencies for tetranucleosomes alone and integrated into each nDFS device with and without 40nM H1.0. These values are plotted in Figure 5B.

| NSI | [H1.0] | Free TNucs | nDFS.C<sub>TNuc</sub> | nDFS.A<sub>TNuc</sub> | nDFS.B<sub>TNuc</sub> |
| --- | --- | --- | --- | --- | --- |
| NSI<sub>3</sub> | 0 | 0 | 0 | 0 | 0.012±0.005 |
| NSI<sub>3</sub> | 40 | 0 | 0.024±0.006 | 0.014±0.008 | 0.021±0.006 |
| NSI<sub>2</sub> | 0 | 0.83±0.04 | 0.84±0.04 | 0.78±0.04 | 0.67±0.04 |
| NSI<sub>2</sub> | 40 | 0.90±0.03 | 0.89±0.04 | 0.79±0.06 | 0.74±0.03 |
| NSI<sub>1</sub> | 0 | 0.16±0.02 | 0.15±0.02 | 0.21±0.02 | 0.29±0.02 |
| NSI<sub>1</sub> | 40 | 0.08±0.01 | 0.08±0.01 | 0.19±0.03 | 0.22±0.02 |
| NSI<sub>0</sub> | 0 | 0.010±0.004 | 0.006±0.003 | 0.016±0.006 | 0.037±0.009 |
| NSI<sub>0</sub> | 40 | 0.024±0.005 | 0.001±0.001 | 0.005±0.005 | 0.018±0.005 |

**Table S4:**
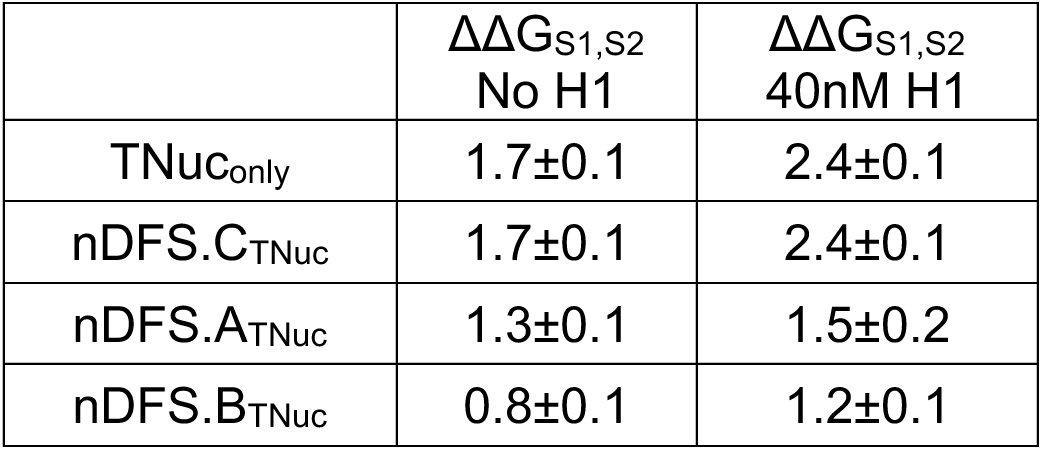
Free energy differences (ΔG_S2-S1_) between NSI_2_ and NSI_1_ from Figure 5C, which were calculated from the measured probabilities in Figure 5B.

**Table S5:**
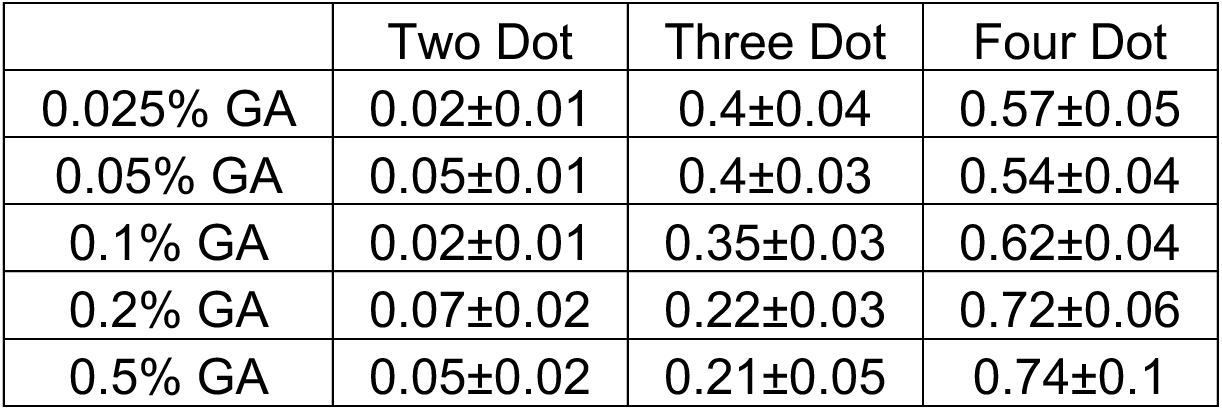
Dot number probability for open array conditions (2mM NaCl) for each Glutaraldehyde (GA) concentration in the bar plot in Figure S4.

|  | Two Dot | Three Dot | Four Dot |
| --- | --- | --- | --- |
| 0.025% GA | 0.02±0.01 | 0.4±0.04 | 0.57±0.05 |
| 0.05% GA | 0.05±0.01 | 0.4±0.03 | 0.54±0.04 |
| 0.1% GA | 0.02±0.01 | 0.35±0.03 | 0.62±0.04 |
| 0.2% GA | 0.07±0.02 | 0.22±0.03 | 0.72±0.06 |
| 0.5% GA | 0.05±0.02 | 0.21±0.05 | 0.74±0.1 |

**Table S6:** Dot number probability for compacted array conditions (200mM NaCl, 1mM MgCl_2_, 40nM H1) for each Glutaraldehyde (GA) concentration in the bar plot in Figure S5.

|  | One Dot | Two Dot | Three Dot | Four Dot |
| --- | --- | --- | --- | --- |
| No GA | 0.01±0.01 | 0.01±0.01 | 0.18±0.04 | 0.80±0.07 |
| 0.02% GA | 0 | 0.05±0.02 | 0.24±0.04 | 0.71±0.06 |
| 0.04% GA | 0 | 0.26±0.04 | 0.39±0.05 | 0.35±0.04 |
| 0.06% GA | 0 | 0.62±0.06 | 0.30±0.04 | 0.08±0.02 |
| 0.08% GA | 0 | 0.74±0.06 | 0.24±0.03 | 0.02±0.01 |
| 0.1% GA | 0 | 0.75±0.06 | 0.25±0.04 | 0.01±0.01 |
| 0.2% GA | 0 | 0.85±0.06 | 0.14±0.02 | 0.02±0.01 |
| 0.4% GA | 0 | 0.88±0.1 | 0.13±0.04 | 0 |

**Table S7:** Incorporation efficiency, which is plotted in Figure 3 for each nanocaliper, as determined by selecting isolated devices from TEM images.

|  | nDFS.C | nDFS.B | nDFS.A |
| --- | --- | --- | --- |
| Properly Incorporated | 0.68±0.03 | 0.45±0.02 | 0.27±0.02 |
| Free Hinge | 0.07±0.01 | 0.11±0.01 | 0.28±0.02 |
| 1 Partially Bound | 0.18±0.01 | 0.30±0.02 | 0.41±0.03 |
| 2 Partially Bound | 0.06±0.01 | 0.13±0.01 | 0.03±0.01 |

## MAIN TEXT REFERENCES

1. Alberts, B. (2008). Molecular biology of the cell, 5th edition by B. Alberts, A. Johnson, J. Lewis, M. Raff, K. Roberts, and P. Walter. Biochemistry and Molecular Biology Education 36. 10.1002/bmb.20192.

2. Minami, K., Semeigazin, A., Nakazato, K., and Maeshima, K. (2025). Euchromatin and Heterochromatin: Implications for DNA Accessibility and Transcription. Preprint at Academic Press, 10.1016/j.jmb.2025.169270 https://doi.org/10.1016/j.jmb.2025.169270.

3. Hsieh, T.H.S., Weiner, A., Lajoie, B., Dekker, J., Friedman, N., and Rando, O.J. (2015). Mapping Nucleosome Resolution Chromosome Folding in Yeast by Micro-C. Cell 162. 10.1016/j.cell.2015.05.048.

4. Ricci, M.A., Manzo, C., García-Parajo, M.F., Lakadamyali, M., and Cosma, M.P. (2015). Chromatin fibers are formed by heterogeneous groups of nucleosomes in vivo. Cell 160. 10.1016/j.cell.2015.01.054.

5. Song, F., Chen, P., Sun, D., Wang, M., Dong, L., Liang, D., Xu, R.M., Zhu, P., and Li, G. (2014). Cryo-EM study of the chromatin fiber reveals a double helix twisted by tetranucleosomal units. Science (1979) 344, 376–380. 10.1126/science.1251413.

6. Kilic, S., Felekyan, S., Doroshenko, O., Boichenko, I., Dimura, M., Vardanyan, H., Bryan, L.C., Arya, G., Seidel, C.A.M., and Fierz, B. (2018). Single-molecule FRET reveals multiscale chromatin dynamics modulated by HP1α. Nat Commun 9. 10.1038/s41467-017-02619-5.

7. Chen, P., Dong, L., Hu, M., Wang, Y.Z., Xiao, X., Zhao, Z., Yan, J., Wang, P.Y., Reinberg, D., Li, M., et al. (2018). Functions of FACT in Breaking the Nucleosome and Maintaining Its Integrity at the Single-Nucleosome Level. Mol Cell 71. 10.1016/j.molcel.2018.06.020.

8. Yang, Z., and Hayes, J.J. (2011). The divalent cations Ca 2+ and Mg 2+ play specific roles in stabilizing histone-DNA interactions within nucleosomes that are partially redundant with the core histone tail domains. Biochemistry 50. 10.1021/bi201377x.

9. Hergeth, S.P., and Schneider, R. (2015). The H1 linker histones: multifunctional proteins beyond the nucleosomal core particle. EMBO Rep 16. 10.15252/embr.201540749.

10. Hao, F., Kale, S., Dimitrov, S., and Hayes, J.J. (2021). Unraveling linker histone interactions in nucleosomes. Preprint, 10.1016/j.sbi.2021.06.001 https://doi.org/10.1016/j.sbi.2021.06.001.

11. Zhou, B.R., and Bai, Y. (2019). Chromatin structures condensed by linker histones. Preprint, 10.1042/EBC20180056 https://doi.org/10.1042/EBC20180056.

12. Dorigo, B., Schalch, T., Kulangara, A., Duda, S., Schroeder, R.R., and Richmond, T.J. (2004). Nucleosome arrays reveal the two-start organization of the chromatin fiber. Science (1979) 306. 10.1126/science.1103124.

13. Chen, Q., Yang, R., Korolev, N., Liu, C.F., and Nordenskiöld, L. (2017). Regulation of Nucleosome Stacking and Chromatin Compaction by the Histone H4 N-Terminal Tail–H2A Acidic Patch Interaction. J Mol Biol 429. 10.1016/j.jmb.2017.03.016.

14. Schalch, T., Duda, S., Sargent, D.F., and Richmond, T.J. (2005). X-ray structure of a tetranucleosome and its implications for the chromatin fibre. Nature 436, 138–141. 10.1038/nature03686.

15. Fierz, B., Chatterjee, C., McGinty, R.K., Bar-Dagan, M., Raleigh, D.P., and Muir, T.W. (2011). Histone H2B ubiquitylation disrupts local and higher-order chromatin compaction. Nat Chem Biol 7. 10.1038/nchembio.501.

16. Poirier, M.G., Oh, E., Tims, H.S., and Widom, J. (2009). Dynamics and function of compact nucleosome arrays. Nat Struct Mol Biol 16, 938–944. 10.1038/nsmb.1650.

17. Cui, Y., and Bustamante, C. (2000). Pulling a single chromatin fiber reveals the forces that maintain its higher-order structure. Proc Natl Acad Sci U S A 97, 127–132. 10.1073/pnas.97.1.127.

18. Kruithof, M., Chien, F.-T., Routh, A., Logie, C., Rhodes, D., and van Noort, J. (2009). Single-molecule force spectroscopy reveals a highly compliant helical folding for the 30-nm chromatin fiber. Nat Struct Mol Biol 16, 534–540. 10.1038/nsmb.1590.

19. Li, W., Chen, P., Yu, J., Dong, L., Liang, D., Feng, J., Yan, J., Wang, P.Y., Li, Q., Zhang, Z., et al. (2016). FACT Remodels the Tetranucleosomal Unit of Chromatin Fibers for Gene Transcription. Mol Cell 64. 10.1016/j.molcel.2016.08.024.

20. Grigoryev, S.A., Arya, G., Correll, S., Woodcock, C.L., and Schlick, T. (2009). Evidence for heteromorphic chromatin fibers from analysis of nucleosome interactions. Proc Natl Acad Sci U S A 106. 10.1073/pnas.0903280106.

21. Ding, X., Lin, X., and Zhang, B. (2021). Stability and folding pathways of tetra-nucleosome from six-dimensional free energy surface. Nat Commun 12. 10.1038/s41467-021-21377-z.

22. Li, Z., Portillo-Ledesma, S., and Schlick, T. (2023). Brownian dynamics simulations of mesoscale chromatin fibers. Biophys J 122. 10.1016/j.bpj.2022.09.013.

23. Lin, X., and Zhang, B. (2024). Explicit ion modeling predicts physicochemical interactions for chromatin organization. Elife 12. 10.7554/elife.90073.3.

24. Qiu, Y., Liu, S., Lin, X., Unarta, I.C., Huang, X., and Zhang, B. (2024). Nucleosome condensate and linker DNA alter chromatin folding pathways and rates. Preprint, 10.1101/2024.11.15.623891 https://doi.org/10.1101/2024.11.15.623891.

25. Engelen, W., and Dietz, H. (2021). Advancing Biophysics Using DNA Origami. Preprint, 10.1146/annurev-biophys-110520-125739 https://doi.org/10.1146/annurev-biophys-110520-125739.

26. Knappe, G.A., Wamhoff, E.C., and Bathe, M. (2023). Functionalizing DNA origami to investigate and interact with biological systems. Preprint, 10.1038/s41578-022-00517-x https://doi.org/10.1038/s41578-022-00517-x.

27. Rothemund, P.W.K. (2006). Folding DNA to create nanoscale shapes and patterns. Preprint, 10.1038/nature04586 https://doi.org/10.1038/nature04586.

28. Castro, C.E., Kilchherr, F., Kim, D.N., Shiao, E.L., Wauer, T., Wortmann, P., Bathe, M., and Dietz, H. (2011). A primer to scaffolded DNA origami. Nat Methods 8, 221–229. 10.1038/nmeth.1570.

29. Zhan, P., Peil, A., Jiang, Q., Wang, D., Mousavi, S., Xiong, Q., Shen, Q., Shang, Y., Ding, B., Lin, C., et al. (2023). Recent Advances in DNA Origami-Engineered Nanomaterials and Applications. Preprint, 10.1021/acs.chemrev.3c00028 https://doi.org/10.1021/acs.chemrev.3c00028.

30. Le, J. V., Luo, Y., Darcy, M.A., Lucas, C.R., Goodwin, M.F., Poirier, M.G., and Castro, C.E. (2016). Probing Nucleosome Stability with a DNA Origami Nanocaliper. ACS Nano 10, 7073–7084. 10.1021/acsnano.6b03218.

31. Wang, Y., Le, J. V., Crocker, K., Darcy, M.A., Halley, P.D., Zhao, D., Andrioff, N., Croy, C., Poirier, M.G., Bundschuh, R., et al. (2021). A nanoscale DNA force spectrometer capable of applying tension and compression on biomolecules. Nucleic Acids Res 49, 8987–8999. 10.1093/nar/gkab656.

32. Darcy, M., Crocker, K., Wang, Y., Le, J. V., Mohammadiroozbahani, G., Abdelhamid, M.A.S., Craggs, T.D., Castro, C.E., Bundschuh, R., and Poirier, M.G. (2022). High-Force Application by a Nanoscale DNA Force Spectrometer. ACS Nano 16, 5682–5695. 10.1021/acsnano.1c10698.

33. Kramm, K., Schröder, T., Gouge, J., Vera, A.M., Gupta, K., Heiss, F.B., Liedl, T., Engel, C., Berger, I., Vannini, A., et al. (2020). DNA origami-based single-molecule force spectroscopy elucidates RNA Polymerase III pre-initiation complex stability. Nat Commun 11. 10.1038/s41467-020-16702-x.

34. Funke, J.J., Ketterer, P., Lieleg, C., Schunter, S., Korber, P., and Dietz, H. (2016). Uncovering the forces between nucleosomes using DNA origami. Sci Adv 2. 10.1126/sciadv.1600974.

35. Funke, J.J., Ketterer, P., Lieleg, C., Korber, P., and Dietz, H. (2016). Exploring Nucleosome Unwrapping Using DNA Origami. Nano Lett 16, 7891–7898. 10.1021/acs.nanolett.6b04169.

36. Poirier, M.G., Bussiek, M., Langowski, J., and Widom, J. (2008). Spontaneous Access to DNA Target Sites in Folded Chromatin Fibers. J Mol Biol 379, 772–786. 10.1016/j.jmb.2008.04.025.

37. Lowary, P.T., and Widom, J. (1998). New DNA sequence rules for high affinity binding to histone octamer and sequence-directed nucleosome positioning. J Mol Biol 276. 10.1006/jmbi.1997.1494.

38. Zhou, B.R., Jiang, J., Ghirlando, R., Norouzi, D., Sathish Yadav, K.N., Feng, H., Wang, R., Zhang, P., Zhurkin, V., and Bai, Y. (2018). Revisit of Reconstituted 30-nm Nucleosome Arrays Reveals an Ensemble of Dynamic Structures. J Mol Biol 430, 3093–3110. 10.1016/j.jmb.2018.06.020.

39. Kästner, J. (2011). Umbrella sampling. Preprint, 10.1002/wcms.66 https://doi.org/10.1002/wcms.66.

40. Kumar, S., Rosenberg, J.M., Bouzida, D., Swendsen, R.H., and Kollman, P.A. (1992). THE weighted histogram analysis method for free-energy calculations on biomolecules. I. The method. J Comput Chem 13, 1011–1021. 10.1002/jcc.540130812.

41. Freeman, G.S., Hinckley, D.M., Lequieu, J.P., Whitmer, J.K., and De Pablo, J.J. (2014). Coarse-grained modeling of DNA curvature. Journal of Chemical Physics 141. 10.1063/1.4897649.

42. Noel, J.K., Levi, M., Raghunathan, M., Lammert, H., Hayes, R.L., Onuchic, J.N., and Whitford, P.C. (2016). SMOG 2: A Versatile Software Package for Generating Structure-Based Models. PLoS Comput Biol 12. 10.1371/journal.pcbi.1004794.

43. Parsons, T., and Zhang, B. (2019). Critical role of histone tail entropy in nucleosome unwinding. Journal of Chemical Physics 150. 10.1063/1.5085663.

44. Liu, S., Lin, X., and Zhang, B. (2022). Chromatin fiber breaks into clutches under tension and crowding. Nucleic Acids Res 50, 9738–9747. 10.1093/nar/gkac725.

45. Happel, N., and Doenecke, D. (2009). Histone H1 and its isoforms: Contribution to chromatin structure and function. Preprint, 10.1016/j.gene.2008.11.003 https://doi.org/10.1016/j.gene.2008.11.003.

46. Cutter, A.R., and Hayes, J.J. (2017). Linker histones: Novel insights into structure-specific recognition of the nucleosome. Biochemistry and Cell Biology 95. 10.1139/bcb-2016-0097.

47. Zhou, B.R., Feng, H., Kale, S., Fox, T., Khant, H., de Val, N., Ghirlando, R., Panchenko, A.R., and Bai, Y. (2021). Distinct Structures and Dynamics of Chromatosomes with Different Human Linker Histone Isoforms. Mol Cell 81. 10.1016/j.molcel.2020.10.038.

48. Allan, J., Hartman, P.G., Crane-robinson, C., and Aviles, F.X. (1980). The structure of histone H1 and its location in chromatin. Nature 288. 10.1038/288675a0.

49. Woodcock, C.L., and Dimitrov, S. (2001). Higher-order structure of chromatin and chromosomes. Preprint, 10.1016/S0959-437X(00)00169-6 https://doi.org/10.1016/S0959-437X(00)00169-6.

50. Grigoryev, S.A., Bascom, G., Buckwalter, J.M., Schubert, M.B., Woodcock, C.L., and Schlick, T. (2016). Hierarchical looping of zigzag nucleosome chains in metaphase chromosomes. Proc Natl Acad Sci U S A 113. 10.1073/pnas.1518280113.

51. Bernier, M., Luo, Y., Nwokelo, K.C., Goodwin, M., Dreher, S.J., Zhang, P., Parthun, M.R., Fondufe-Mittendorf, Y., Ottesen, J.J., and Poirier, M.G. (2015). Linker histone H1 and H3K56 acetylation are antagonistic regulators of nucleosome dynamics. Nat Commun 6. 10.1038/ncomms10152.

52. Burge, N.L., Thuma, J.L., Hong, Z.Z., Jamison, K.B., Ottesen, J.J., and Poirier, M.G. (2022). H1.0 C Terminal Domain Is Integral for Altering Transcription Factor Binding within Nucleosomes. Biochemistry 61. 10.1021/acs.biochem.2c00001.

53. Yue, H., Fang, H., Wei, S., Hayes, J.J., and Lee, T.H. (2016). Single-Molecule Studies of the Linker Histone H1 Binding to DNA and the Nucleosome. Biochemistry 55. 10.1021/acs.biochem.5b01247.

54. Luger, K., Mäder, A.W., Richmond, R.K., Sargent, D.F., and Richmond, T.J. (1997). Crystal structure of the nucleosome core particle at 2.8 Å resolution. Nature 389, 251–260. 10.1038/38444.

55. McGinty, R.K., and Tan, S. (2016). Recognition of the nucleosome by chromatin factors and enzymes. Preprint, 10.1016/j.sbi.2015.11.014 https://doi.org/10.1016/j.sbi.2015.11.014.

56. Dorigo, B., Schalch, T., Bystricky, K., and Richmond, T.J. (2003). Chromatin fiber folding: Requirement for the histone H4 N-terminal tail. J Mol Biol 327. 10.1016/S0022-2836(03)00025-1.

57. Shogren-Knaak, M., Ishii, H., Sun, J.M., Pazin, M.J., Davie, J.R., and Peterson, C.L. (2006). Histone H4-K16 acetylation controls chromatin structure and protein interactions. Science (1979) 311. 10.1126/science.1124000.

58. Meng, H., Andresen, K., and Van Noort, J. (2015). Quantitative analysis of single-molecule force spectroscopy on folded chromatin fibers. Nucleic Acids Res 43, 3578–3590. 10.1093/nar/gkv215.

59. Brouwer, T., Pham, C., Kaczmarczyk, A., De Voogd, W.J., Botto, M., Vizjak, P., Mueller-Planitz, F., and Van Noort, J. (2021). A critical role for linker DNA in higher-order folding of chromatin fibers. Nucleic Acids Res 49, 2537–2551. 10.1093/nar/gkab058.

60. Moller, J., Lequieu, J., and De Pablo, J.J. (2019). The Free Energy Landscape of Internucleosome Interactions and Its Relation to Chromatin Fiber Structure. ACS Cent Sci 5. 10.1021/acscentsci.8b00836.

## METHOD REFERENCES

1. Wang, Y., Le, J. V., Crocker, K., Darcy, M.A., Halley, P.D., Zhao, D., Andrioff, N., Croy, C., Poirier, M.G., Bundschuh, R., et al. (2021). A nanoscale DNA force spectrometer capable of applying tension and compression on biomolecules. Nucleic Acids Res 49, 8987–8999. 10.1093/nar/gkab656.

2. Darcy, M., Crocker, K., Wang, Y., Le, J. V., Mohammadiroozbahani, G., Abdelhamid, M.A.S., Craggs, T.D., Castro, C.E., Bundschuh, R., and Poirier, M.G. (2022). High-Force Application by a Nanoscale DNA Force Spectrometer. ACS Nano 16, 5682–5695. 10.1021/acsnano.1c10698.

3. Douglas, S.M., Marblestone, A.H., Teerapittayanon, S., Vazquez, A., Church, G.M., and Shih, W.M. (2009). Rapid prototyping of 3D DNA-origami shapes with caDNAno. Nucleic Acids Res 37, 5001–5006. 10.1093/nar/gkp436.

4. Poirier, M.G., Bussiek, M., Langowski, J., and Widom, J. (2008). Spontaneous Access to DNA Target Sites in Folded Chromatin Fibers. J Mol Biol 379, 772–786. 10.1016/j.jmb.2008.04.025.

5. Luo, Y., North, J.A., and Poirier, M.G. (2014). Single molecule fluorescence methodologies for investigating transcription factor binding kinetics to nucleosomes and DNA. Methods 70, 108–118. 10.1016/j.ymeth.2014.09.011.

6. Gibson, M.D., Brehove, M., Luo, Y., North, J., and Poirier, M.G. (2016). Methods for Investigating DNA Accessibility with Single Nucleosomes. In Methods in Enzymology, pp. 379–415. 10.1016/bs.mie.2016.08.014.

7. Poirier, M.G., Oh, E., Tims, H.S., and Widom, J. (2009). Dynamics and function of compact nucleosome arrays. Nat Struct Mol Biol 16, 938–944. 10.1038/nsmb.1650.

8. Caravaca, J.M., Caño, S., Gállego, I., and Daban, J.R. (2005). Structural elements of bulk chromatin within metaphase chromosomes. Chromosome Research 13. 10.1007/s10577-005-1008-3.

9. Song, F., Chen, P., Sun, D., Wang, M., Dong, L., Liang, D., Xu, R.M., Zhu, P., and Li, G. (2014). Cryo-EM study of the chromatin fiber reveals a double helix twisted by tetranucleosomal units. Science (1979) 344, 376–380. 10.1126/science.1251413.

10. Jentink, N., Purnell, C., Kable, B., Swulius, M.T., and Grigoryev, S.A. (2023). Cryoelectron tomography reveals the multiplex anatomy of condensed native chromatin and its unfolding by histone citrullination. Mol Cell 83. 10.1016/j.molcel.2023.08.017.

11. Zhou, B.R., Jiang, J., Ghirlando, R., Norouzi, D., Sathish Yadav, K.N., Feng, H., Wang, R., Zhang, P., Zhurkin, V., and Bai, Y. (2018). Revisit of Reconstituted 30-nm Nucleosome Arrays Reveals an Ensemble of Dynamic Structures. J Mol Biol 430, 3093–3110. 10.1016/j.jmb.2018.06.020.

12. Ding, X., Lin, X., and Zhang, B. (2021). Stability and folding pathways of tetra-nucleosome from six-dimensional free energy surface. Nat Commun 12. 10.1038/s41467-021-21377-z.

13. Eastman, P., Galvelis, R., Peláez, R.P., Abreu, C.R.A., Farr, S.E., Gallicchio, E., Gorenko, A., Henry, M.M., Hu, F., Huang, J., et al. (2024). OpenMM 8: Molecular Dynamics Simulation with Machine Learning Potentials. Journal of Physical Chemistry B 128. 10.1021/acs.jpcb.3c06662.

14. Liu, S., Wang, C., Latham, A.P., Ding, X., and Zhang, B. (2023). OpenABC enables flexible, simplified, and efficient GPU accelerated simulations of biomolecular condensates. PLoS Comput Biol 19. 10.1371/journal.pcbi.1011442.

15. Ding, X., Vilseck, J.Z., and Brooks, C.L. (2019). Fast Solver for Large Scale Multistate Bennett Acceptance Ratio Equations. J Chem Theory Comput 15. 10.1021/acs.jctc.8b01010.

